# Deep Evolutionary History of the Phox and Bem1 (PB1) Domain Across Eukaryotes

**DOI:** 10.1101/2020.01.13.903906

**Authors:** Sumanth Kumar Mutte, Dolf Weijers

## Abstract

Protein oligomerization is a fundamental process to build complex functional modules. Domains that facilitate the oligomerization process are diverse and widespread in nature across all kingdoms of life. One such domain is the Phox and Bem1 (PB1) domain, which is functionally (relatively) well understood in the animal kingdom. However, beyond animals, neither the origin nor the evolutionary patterns of PB1-containing proteins are understood. While PB1 domain proteins have been found in other kingdoms, including plants, it is unclear how these relate to animal PB1 proteins.

To address this question, we utilized large transcriptome datasets along with the proteomes of a broad range of species. We discovered eight PB1 domain-containing protein families in plants, along with three each in Protozoa and Chromista and four families in Fungi. Studying the deep evolutionary history of PB1 domains throughout eukaryotes revealed the presence of at least two, but likely three, ancestral PB1 copies in the Last Eukaryotic Common Ancestor (LECA). These three ancestral copies gave rise to multiple orthologues later in evolution. Tertiary structural models of these plant PB1 families, combined with Random Forest based classification, indicated family-specific differences attributed to the length of PB1 domain and the proportion of β-sheets.

This study identifies novel PB1 families and reveals considerable complexity in the protein oligomerization potential at the origin of eukaryotes. The newly identified relationships provide an evolutionary basis to understand the diverse functional interactions of key regulatory proteins carrying PB1 domains across eukaryotic life.

## BACKGROUND

Protein-protein interaction is a basic and important mechanism that brings proteins together in a functional module and thus allows the development of higher-order functionalities. One of the versatile interaction domains that brings this modularity through either dimerization or oligomerization is the PB1 domain. Initially, the two animal proteins, p40^Phox^ and p67^Phox^, were shown to interact through a novel motif that contains a stretch of negatively charged amino acids [1]. In the same study, it was also shown that the yeast CELL DIVISION CONTROL 24 (Cdc24) protein contains the same motif as found in p40^Phox^, and hence named as PC motif (for p40^Phox^ and Cdc24; Nakamura et al. 1998). Later, the BUD EMERGENCE 1 (Bem1) protein in yeast was also found to have this motif, after which it has been renamed as PB1 domain (for Phox and Bem1). The PB1 domain of Bem1 in yeast is required for the interaction with Cdc24 to maintain cell polarity [2]. Later, in mammals, many protein families were identified that contain a PB1 domain [3]. In plants, the PB1 domain was initially recognised as domain III/IV in the auxin repressor proteins AUXIN/INDOLE-3-ACETIC-ACID (Aux/IAAs) [4]. Later, the domains III/IV were found to form a fold similar to (and hence renamed as) the PB1 domain, with a wider presence in multiple gene families in plants [5–7].

The PB1 domain ranges from 80-100 amino acids in length and exhibits a Ubiquitin β-grasp fold with five β-sheets and two α-helices [8,9]. The first half of the domain represents a positively charged face, with a conserved Lysine (K) in β1. The latter half of the domain represents a negatively charged face, with D-x-D/E-x-D/E as core (OPCA motif; Müller et al.2006). Based on the presence/absence of these important residues/motifs, the PB1 domains are divided into three types. If the PB1 domain contains only the conserved OPCA motif but not the Lysine, it is considered as a type-1 (or type-A) PB1 domain. If there is only Lysine but not OPCA motif, it is a type-2 (or type-B) domain. If the PB1 domain contains both Lysine and the OPCA motifs, it is referred as type-1/2 (or type-AB). Various proteins that harbour a PB1 domain undergo dimerization or oligomerization, where the positive face of one PB1 domain interacts with the negative face of another in a head-to-tail fashion [3,8]. Hence, depending on the type of PB1 domain they interact with, there can be either homotypic or heterotypic PB1 interactions.

All the eukaryotes are divided into five kingdoms: Protozoa, Chromista, Fungi, Animalia and Plantae. PB1 domain-containing proteins have been relatively well studied in Animalia, when compared to the other kingdoms. At least nine gene families have been shown to encode a PB1 domain [10]. Animal genomes encode proteins that contain all three types of PB1 domains: type-1 - NEUTROPHIL CYTOSOL FACTOR 4 (NCF4/p40^Phox^), MITOGEN-ACTIVATED PROTEIN KINASE KINASE 5 (M2K5) and NEXT TO BRCA 1 (NBR1); type-2 - NEUTROPHIL CYTOSOL FACTOR 2 (NCF2/p67^Phox^), PARTITIONING DEFECTIVE 6 (Par6) and MITOGEN-ACTIVATED PROTEIN KINASE KINASE KINASE 2/3 (M3K2/3); type-1/2 - SEQUESTOSOME-1 (SQSTM1/p62), ATYPICAL PROTEIN KINASE C (aPKC) and TRK-FUSED GENE (TFG). A systematic analysis through yeast two-hybrid and pull-down assays revealed various homotypic and heterotypic interactions among these PB1 domains [3]. The p67^Phox^ upon its interaction with p40^Phox^ activates the phagocyte NADPH oxidase that is important for innate immunity in mammals [11]. The Par6-aPKC complex establishment through PB1 is essential for cell polarity in mammals and insects [12]. This complex, along with Par3, also regulates the formation of junctions through apical-basal polarity in mammalian epithelial cells [13]. p62 acts as a crucial scaffolding protein playing important roles in autophagy, apoptosis and inflammation [14].

The PB1 domain of M3K2/3 interacts with M2K5 to activate ERK5 mediated signalling in response to growth factors and osmotic stress [15]. TFG PB1 domain is involved in transforming activity by forming the TFG-TrkA (Tyrosine Kinase A) fusion [16]. NBR1 interacts with p62 through PB1 which is required for targeting p62 to sarcomeres [3]. Few non-canonical PB1 interactions were also observed, for example, in p40^Phox^ PB1 and PX domains undergo intramolecular interaction, disruption of which is required to activate the NADPH oxidase [17]. In yeast, interaction of both the PB1 domain containing proteins, Bem1 and Cdc24 is critical for the cell polarity establishment at both budding and mating [2]. The NADPH OXIDASE REGULATOR (NoxR) plays a central role fungal morphogenesis, growth and development through NADPH oxidation pathway [18].

The best-studied PB1 domains in plants are from the AUXIN RESPONSE FACTOR (ARF) transcription factors and their AUXIN/INDOLE-3-ACETIC-ACID (Aux/IAA) inhibitors. Both the homotypic and heterotypic interactions among and between these gene families is relatively well established [19]. The structural basis for these interactions has also been scrutinized in detail [8,20]. Both ARFs and Aux/IAAs are involved in auxin-dependent gene regulation through the Nuclear Auxin Pathway, that controls various growth and developmental processes (reviewed in Weijers and Wagner 2016). Another PB1 domain containing protein, AtNBR1, an Arabidopsis ortholog of animal NBR1, is involved in autophagy and was shown to homo-polymerize through its PB1 domain [5]. Joka2, an AtNBR1 orthologue of tobacco, can also homodimerize through its PB1 domain [7]. Moreover, this study also revealed non-canonical interaction of the PB1 domain with the C-terminal UBA domain within the same protein [7]. Homotypic interactions through PB1 domains of NIN-LIKE PROTEINS OF PLANTS (NLPs) are required to induce nitrate-dependent gene expression [22,23]. Interestingly, like AtNBR1/Joka2, the PB1 domain of NLP also undergoes non-canonical interaction with the HQ domain of TEOSINTE BRANCHED 1, CYCLOIDEA, PCF DOMAINS CONTAINING PROTEIN 20 (TCP20) [22]. Another study identified a novel unclassified PB1 domain-containing protein PAL OF QUIRKY (POQ) that undergoes non-canonical interaction with QUIRKY (QKY) [24]. However, the structural or mechanistic basis of these non-canonical interactions are yet to be elucidated.

Even though PB1 domain proteins are well defined and their mechanical basis is relatively well established in animals (reviewed in Sumimoto et al. 2007; Burke and Berk 2015), their evolutionary histories are essentially unknown. Moreover, it is unclear how many PB1 domain-containing gene families are present in other kingdoms. Deep evolution has been relatively well studied for ARF and Aux/IAA gene families [26] and to a certain extent for NLPs [27], but the presence and the evolution of other PB1 domains, if any, in plants and unicellular eukaryotes is obscure. Hence, the current study is designed to address several important questions related to the distribution and ancestry of PB1 domains in the eukaryotic tree of life: (1) How many PB1 domain-containing gene families are present in the kingdoms Protozoa, Chromista, Fungi and Plantae? (2) What is the origin of the PB1 domain? How many copies of PB1 were present in the Last Eukaryotic Common Ancestor (LECA)? (3) How have PB1-containing proteins diversified/multiplied in evolution across multiple kingdoms? (4) What are the sequence/structural patterns specific to each family of PB1s and how to classify them?

To answer these questions, we have utilized the large transcriptome datasets in Chromista and Plantae and the (almost) complete proteomes from Fungi and Animalia. We found that the PB1 domains have a deep evolutionary origin with at least two copies in LECA. Moreover, we find that the PB1 domain is associated with a variety of domains, ranging from DNA-binding domains to Kinases and membrane-binding domains. Further, a detailed sequence analysis of PB1 domains in Plantae revealed that these are poorly conserved among various families in general, with few residues being specific to each family. Taken together, this study provides the first evolutionary framework of the PB1 domains across the eukaryotes.

## RESULTS

### Identification and evolution of PB1 domain-containing proteins in various kingdoms

#### Animalia

Based on literature, we extracted protein sequences of all PB1 domain-containing proteins in the human genome from the Uniprot database. Nine gene families were found to encode the PB1 domains as a part of their protein architecture (Fig. 1a). aPKC and M3K2/3 both contain PB1 and Kinase domains in their N- and C-terminus, respectively. Whereas, aPKC contains an extra diacylglycerol-binding (kDAG) domain in the middle. NCF2/p67^Phox^ and NCF4/p40^Phox^ both contains SRC Homology 3 (SH3) and PB1 domains in the C-terminus, where NCF2 contains Tetratricopeptides and NCF4 contains Phox homologous domain (PX) in their N-terminus (Fig. 1a). The other three protein families, Par6, TFG and p62/SQSTM1, are in general shorter than other PB1 domain-containing proteins, with a PB1 domain in the amino-end. p62 contains a Ubiquitin-associated domain (UBA), whereas Par6 contains a PSD95-Dlg1-Zo1 (PDZ) domain in the carboxy-end (Fig. 1a). The full name or description of all the domains along with a link to the InterPro domain database are provided online (see Additional file 1: Table S1).

**Figure 1:**
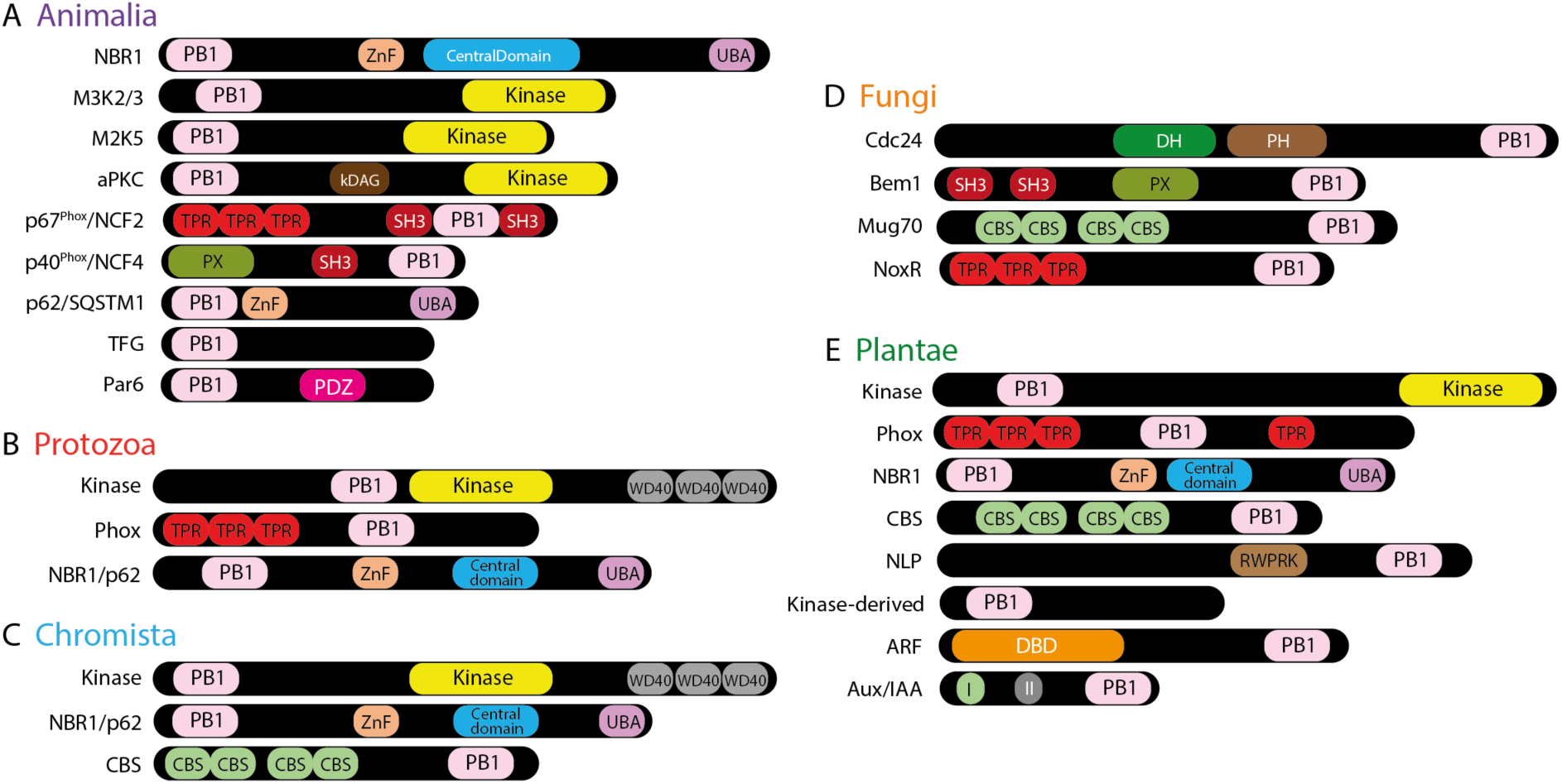
Domain architecture of various PB1 domain containing proteins across the five eukaryotic kingdoms. Panels show he complement of PB1-containing proteins in Animalia (A), Protozoa (B), Chromista (C), Fungi (D) and Plantae (E). NBR1/p62 in (A) and (B) indicate the presence of both NBR1 and p62 with all the domains as their orthologs along with multiple partial domain combinations. DBD in (C) represents the DNA binding domain, which is a combination of both B3 and dimerization domain (DD). Abbreviation and the corresponding InterPro database link of all the domains are provided in Additional file 1: Table S1. PB1 domains that are identified in only one sequence and/or one species are provided in the Additional file 1: Fig. S1.

PB1 sequences from all the above-mentioned proteins were used as queries to retrieve orthologues from ten species across various phyla in Animalia (see Additional file 4 for the list of species used). Retrieved orthologous sequences were used in a phylogenetic analysis along with the respective human counterparts. The PB1 domain-based phylogeny reflected the monophyly of each gene family (Fig. 2 and Additional file 1: Fig. S2). PB1 domains of M3K2-M3K3 and aPKC-M2K5 form paralogous pairs, indicating the common ancestry of PB1 for each pair at the emergence of the kingdom Animalia. Interestingly, the paralog ous pairs M3K2-M3K3 and aPKC-M2K5 PB1 domains are closer to the respective orthologues from other kingdoms than the other PB1 domains in the same kingdom, Animalia. A similar trend is observed with NBR1, however, surprisingly NCF2/p67^Phox^ is placed as the sister clade to the NBR1. p62 do not show any close relationship with other PB1 domains, neither paralogous nor orthologous, from the same kingdom or the other kingdoms (Fig. 2 and Additional file 1: Fig. S2). In a similar way, Par6 and TFG also appear to be Animalia-specific clades (Fig. 2 and Fig. 3).

**Figure 2:**
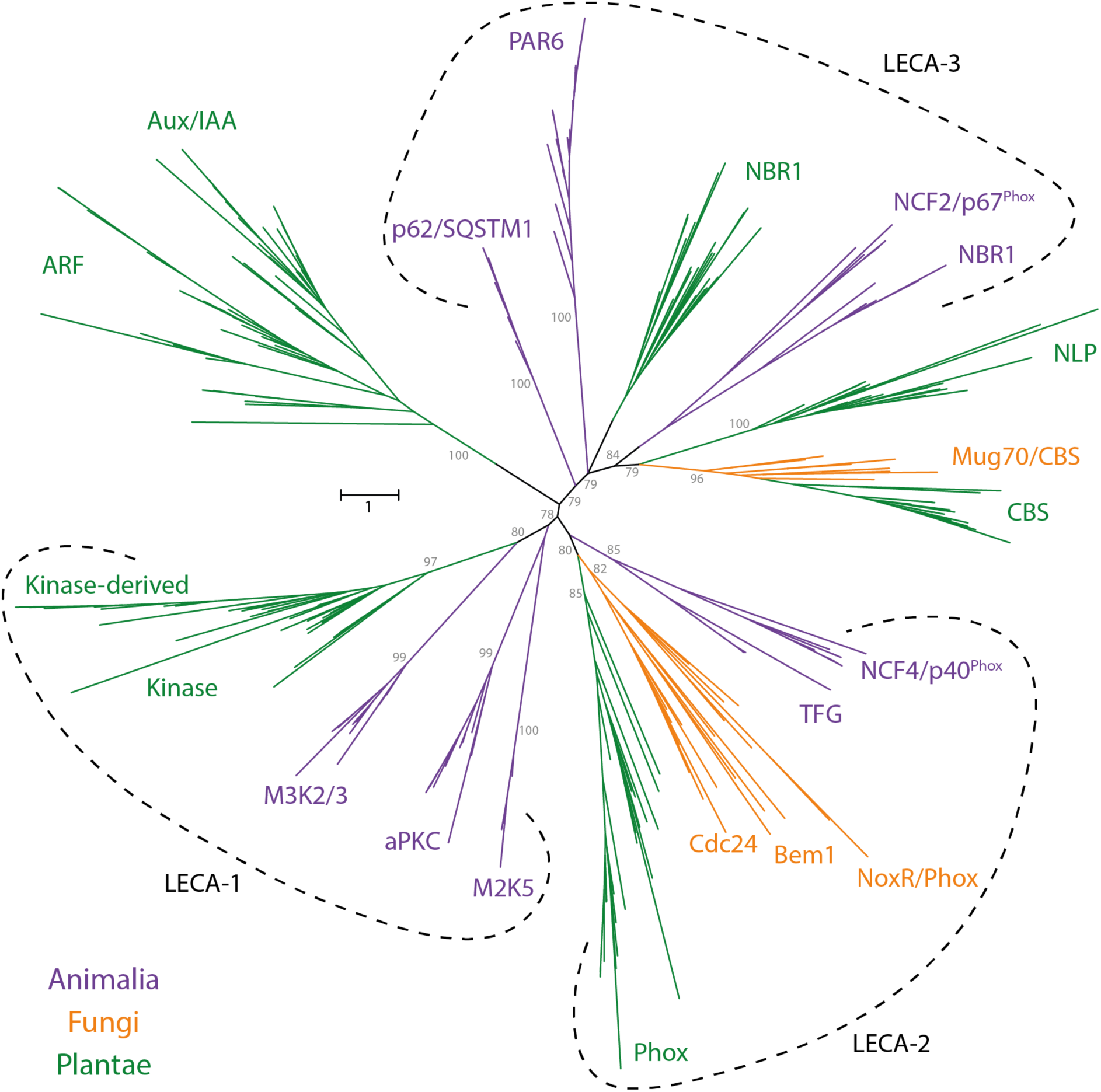
Unrooted tree with representative Fungi, Animalia and Plantae PB1 domains. Early branches that are well-supported (bootstrap >75) are indicated in grey. Orthologs from each kingdom are represented with each colour as indicated: Fungi in ‘orange’, Animalia in ‘purple’ and Plantae in ‘green’. The groups outlined with dotted lines indicated as LECA-1, LECA-2 and LECA-3 represent the probable ancestral copies in LECA corresponding to Kinase, Phox and NBR1 groups respectively. Another phylogenetic tree with all the five kingdoms is presented in the Additional file 1: Fig. S2 as a schematic and full version with taxa names and domain information of both the trees are available online (Additional file 2 and Additional file 3).

**Figure 3:**
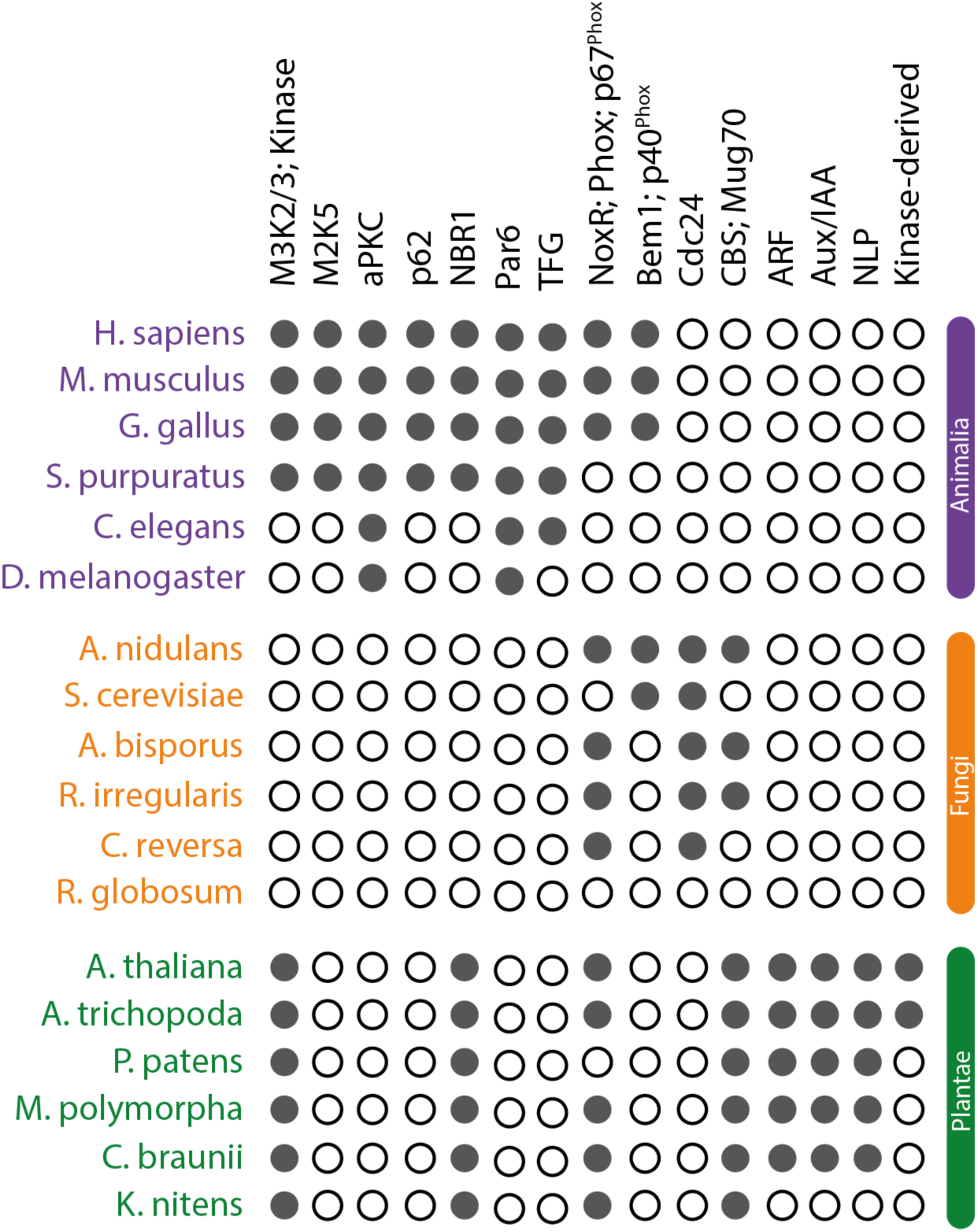
Summary of various PB1 domain-containing proteins across key species in Animalia, Fungi and Plantae. Filled and empty circles represent presence and absence, respectively, of the orthologous genes for the family mentioned on the top in the corresponding species.

#### Protozoa

From UniProt, at least six reference proteomes and other individual Protozoan sequences from various species across different phyla in Protozoa were used to identify the PB1 domains (see Additional file 4). Few PB1 domain-containing proteins were identified along with a large number of partial (or truncated) proteins with either only a PB1 domain or a large unknown flanking sequence (Fig. 1 and Additional file 1: Fig. S1). Among the (full length) PB1 domain-containing proteins, orthologs of Animalia M3K2/3 as well as Plantae Phox were identified, and named as Kinase and Phox respectively (Fig. 1b and Additional file 1: Fig. S2). Unlike the animals, the Protozoan kinases also contain WD40 repeats at their C-terminus. Moreover, the PB1 domain is also adjacent to kinase domain (Fig. 1b). Orthologues of the NBR1 with all the four (known) domains were also found, along with the sequences of various domain combinations i.e. either PB1 with Zn-finger, with the NBR1 Central domain, or with the UBiquitin Associated (UBA) domains (Fig. 1). We also identified many PB1 domain-containing proteins either associated with a Sterile Alpha Motif (SAM), Per-Arnt-Sim (PAS), EF-hand or Cystathionine Beta Synthase (CBS) domains (Additional file 1: Fig. S1). However, the majority of these are identified in only one sequence or one species, and also represented in polyphyletic groups spread across the phylogenetic tree, making it difficult to classify them into a certain clade (Additional file 1: Fig. S2).

#### Chromista

To identify the PB1 domains in Chromista, all the transcriptomes from the MMETSP database were used (see Additional file 4). Well-annotated fungal (Bem1 and Cdc24), plant (Arabidopsis and Marchantia) and animal (Human and Mouse) PB1 sequences were used to query the database, and processed the data with the pipeline developed previously [26]. Two gene families, Kinase and NBR1 were identified in Chromista, with a similar domain architecture, being orthologous to the respective gene families in Animalia (Fig. 1c and Additional file 1: Fig. S2). Interestingly, like Protozoan kinase proteins, these carry WD40 repeats too, however, the PB1 domain is far to the N-terminus as in Plantae or Animalia (Fig. 1c). Orthologues of NBR1 are identified as multiple (partial) proteins represented as polyphyletic groups, similar to Protozoa (Additional file 1: Fig. S2). However, as a third gene family, we identified the CBS domain-containing proteins, where the PB1 domain is associated in their carboxy terminus (Fig. 1c). Few other PB1 domain proteins were identified as single copies in only one species that host either a Tetratricopeptide (TPR) repeat or an EF-hand domain, which were also represented in polyphyletic groups (Additional file 1: Fig. S1 and Additional file 1: Fig. S2).

#### Fungi

For Fungi, we have selected the 12 reference proteomes from MycoCosm database (see Additional file 4). Well-annotated plant and animal PB1 sequences were used as query sequences. Four PB1 domain containing protein families were identified (Fig. 1d). The widely known Bem1 and Cdc24, were identified as a monophyletic paralogous pair in our study (Fig. 2). Along with the PB1 domain, Bem1 contains SH3 and PX domains, whereas Cdc24 contains Dbl homology (DH) and Plextrin homology (PH) domains. Interestingly, NCF4/p40^Phox^, the animal ortholog of Bem1, contains PX in N-terminus, unlike in the middle as in Bem1 (Fig. 1d). CBS domain containing proteins (referred as Mug70) were also identified, with a similar domain architecture like in other kingdoms (Fig. 1d). Further, NoxR, an ortholog of Animalia and Plantae Phox, was also identified as a sister clade to this pair (Fig. 2 and Additional file 1: Fig. S2). In summary, all the four gene families form a respective individual monophyletic group with all the paralogs, indicating their presence across major phyla in Fungi (Fig. 2 and Fig. 3). It is worth noting that an ortholog of p62 and a PB1 domain associated with a SAM domain were identified. However, each was found in only one species and a single copy (Additional file 1: Fig. S1). Hence, we discarded them for further analysis as they are considered of low confidence and may not represent any phylum or the kingdom itself.

#### Plantae

To identify all the PB1 domains in the kingdom Plantae (Fig. 1e), we have adapted a similar pipeline as mentioned above [26], using 485 transcriptomes, that belong to multiple phyla in the kingdom Plantae, from the OneKP database (Matasci et al. 2014; see Additional file 4). We identified eight gene families that encode for PB1 domain-containing proteins in plants (Fig. 1e). Among these, NBR1 and Kinase orthologues are placed in the same clade as their counterparts from other kingdoms, and also contain the same domain architecture as their animal orthologs (Fig. 1e and Fig. 2). ARF and Aux/IAA families form a distinct monophyletic clade indicating a common ancestry at the base of the Streptophytes. ARFs contain a B3 and Dimerization domains (together referred as DNA-binding domain (DBD)) at the N-terminus and a PB1 domain at the C-terminus, like the Aux/IAAs. In addition to a PB1 domain, Aux/IAAs also contain an EAR-motif and a degron motif (Domain-I and −II respectively; Fig. 1e). Phox proteins, having the same domains as animal counterparts, form a sister clade to the respective orthologous proteins from other kingdoms (Fig. 2). CBS domain-containing proteins were also identified in plants, placed in the same clade as fungal Mug70. Kinase-derived, ARF, Aux/IAA and NLP are Plantae specific families that are not identified in any other kingdom (Fig. 3). All these plant-specific gene families were discovered before, except Kinase-derived, which has only a PB1 domain in its N-terminus with a large flanking sequence without any known domains. It is worth mentioning that the Kinase-derived PB1 domains, resemble the Kinase PB1 domains and appears to have been duplicated in the phylum ‘Magnoliophyta’ (Angiosperms; Fig. 2 and Fig. 3). NLPs contain an RWP-RK domain, in association with the PB1 domain and they are placed as sister clade to the CBS domain containing proteins (Fig. 1e and Fig. 2). Interestingly, among all the families identified so far across all the kingdoms, there do not appear to be any constraints on either the position of the PB1 domain in the protein, or the category of domains it is associated with (DNA binding, Oligomerization, Phosphorylation etc.; Fig. 1). An overview of all the identified gene families and their existence across the major phyla in the kingdoms Animalia, Fungi and Plantae is summarized in Figure 3.

### Ancestral copy number in LECA

To better understand the origin and evolutionary patterns of all the PB1 domains across five kingdoms in eukaryotes, two phylogenetic trees were constructed using only the PB1 domain protein sequences. One is based on the PB1 domains from only three kingdoms (Animalia, Fungi and Plantae; Fig. 2), whereas another one is constructed based on all the sequences from five kingdoms (Additional file 1: Fig. S2). The detailed versions of both the phylogenetic trees are available online (Additional file 2 and Additional file 3). All the previously mentioned pairs that form the monophyletic groups of individual families in each kingdom are well supported with good bootstrap values (>75), especially in Fungi, Animalia and Plantae (Fig. 2). The branches representing PB1 domains in Protozoa and Chromista are highly unreliable due to the polyphyletic nature and their random distribution across the phylogenetic tree (Additional file 1: Fig. S2). Overall, the recently evolved clades in the phylogeny that are either gene family-specific or kingdom-specific, are generally monophyletic in nature. We have observed a decrease in the support of the split of early branches (with poor bootstraps) in the phylogeny based on all the five kingdoms (Additional file 1: Fig. S2 and Additional file 3). Monophyletic grouping, as well as the presence in multiple kingdoms, support the notion that there would have been at least two common ancestral copies of PB1 domains, each corresponding to Kinase and Phox orthologues across eukaryotes. Even though the Plantae NBR1 PB1 domains, along with the animal orthologues (and the similar proteins p62) are not monophyletic in origin, they are still placed in the phylogeny as sister clades (Fig. 2). This distribution of orthologues from the various kingdoms hint at a third common ancestor of PB1 in LECA. This analysis has failed to predict the order of evolution because of the lack of sufficient phylogenetic signal due to poorly conserved sequences, a relatively small domain (in general) and poor bootstraps in the early branches in the tree with all five kingdoms. The use of bacterial outgroup sequences could not improve resolution, leading to mixing in the phylogeny along with ingroup sequences. Hence, no outgroup was used and the tree is unrooted. Because of these drawbacks, this study could not identify the order of events, but could predict the copy number in LECA, based on both the monophyletic nature of Kinase and Phox groups as well as presence of NBR1 orthologous sister clades across multiple kingdoms.

### (Dis)similarities in the plant PB1 domains

After identifying the majority, if not all, of the PB1 domain-containing proteins and understanding their evolution patterns across major phyla in all five kingdoms in eukaryotes, we further investigated the plant PB1 domains in detail at the amino acid level. To achieve this, we gathered the PB1 domain sequences from four whole genome-sequenced land plants, one species each from liverworts (*Marchantia polymorpha*), mosses (*Physcomitrella patens*), basal angiosperms (*Amborella trichopoda*) and a core eudicot (*Arabidopsis thaliana*). All PB1 domain protein sequences that belongs to the eight families identified were aligned, and an individual sequence logo was derived for each family (Fig. 4). The well-conserved (group of) residues across the majority of the families are the positive residues Lysine (K) in β1 and Arginine (R) in β2 that together represent the positive surface. However, the Lysine of β1 that makes contact with the OPCA motif on the negative face of another PB1, is not conserved in Kinase and Kinase-derived PB1 domains, indicating that these could be type-1 PB1 domains with only a conserved negative face (Fig. 4). In general, the negative face represented by the OPCA motif is relatively well-conserved in all the gene families, despite a strong conservation of three amino acids (QLP) just before the OPCA motif in Kinase and Kinase-derived PB1 domains (Fig. 4). Interestingly, the Tyrosine (Y) in β3 is relatively well conserved in all the gene families. Apart from these generally conserved residues across multiple families, there are various single amino acids that are specifically conserved in each gene family. For example, Tyrosine (Y) and Phenylalanine (F) of the α1, Glycine (G) in β4 and Phenylalanine (F) in α2 are specific to ARF and Aux/IAA PB1 domains. In a similar way, two Phenylalanine (F) in and before the β1 and a G-x-L-x-L-x-L motif in β5 are specific to PB1 domains associated with CBS domains (Fig. 4). The Tryptophan (W) in β4 is specific to NLPs. Despite NBR1 being a single-copy gene in the kingdom Plantae, there do not seem to be any constrains on the domain itself, as there is less than 20% identity among them. This provides a basic understanding of relaxed evolutionary pressure in the PB1 domain, providing opportunities for many gene family-specific changes. This makes it difficult not only to predict general sequence patterns that are important for function, but also to estimate the domain properties specific to each family purely based on the primary sequence and its poorly conserved amino acids.

**Figure 4:**
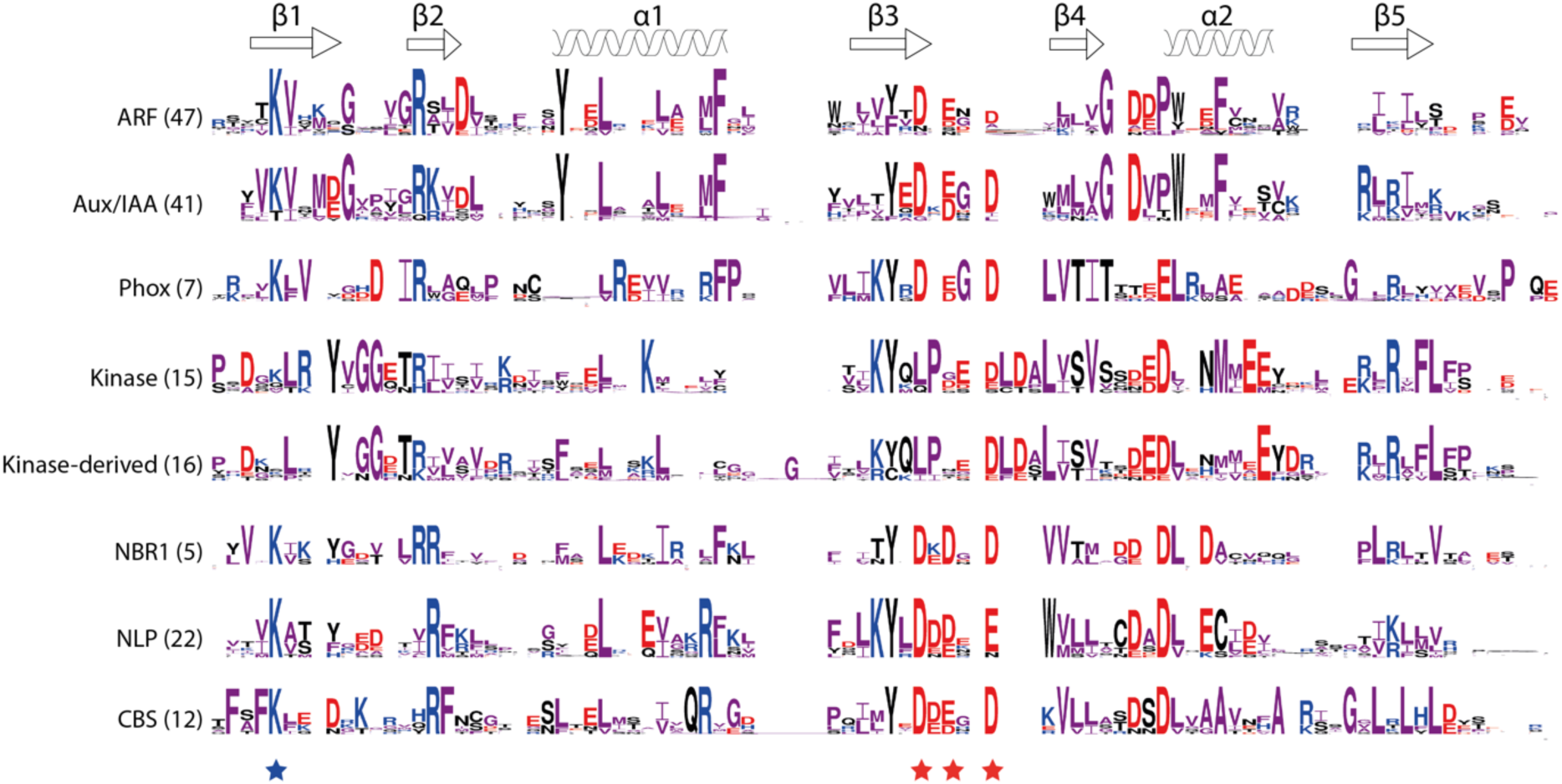
Sequence logos based on the alignment of PB1 domains from the representative land plants. (Marchantia, Physco, Amborella and Arabidopsis). Secondary structures (α-helices and β-sheets) represented on the top are based on the ARF5 structure (PDB ID: 4CHK). Numbers represented in the braces next to the name of the gene family, shows the number of sequences present in all these four species together, and also the number of sequences used for that particular alignment logo. Amino acids are coloured according to the group: ‘PAGFLIMV’, ‘KRH’ and ‘DE’ are shown in ‘purple’, ‘blue’ and ‘red’ colours respectively. All other amino acids are shown in ‘black’. Stars at the bottom represent the key residues on positive (blue) and negative (red) faces, corresponding to Lysine and OPCA motif (D-x-D/E-x-D/E core) respectively.

### Classification using Random Forests

Since there are no clear patterns to identify the gene family to which each PB1 belongs to and because it is also not possible to identify important features of a specific PB1 domain based on the sequence alignment, one might detect patterns based on the secondary structure composition along with the amino acid properties. Random forest (RF) based classification was performed with 28 amino acid descriptors as variables. After bootstrap aggregating (bagging) all the decision trees from the RF, the mean out-of-bag (OOB) error rate is only 6% which indicates the high reliability of the RF model (Fig. 5). The classification error rate is the highest (∼14%) for Kinase-derived and the least (∼2%) for ARF PB1 domains (Fig. 5a and Additional file 1: Table S3). On an average, the majority of PB1 families were resolved well, indicating the high reliability of classification using these descriptors.

**Figure 5:**
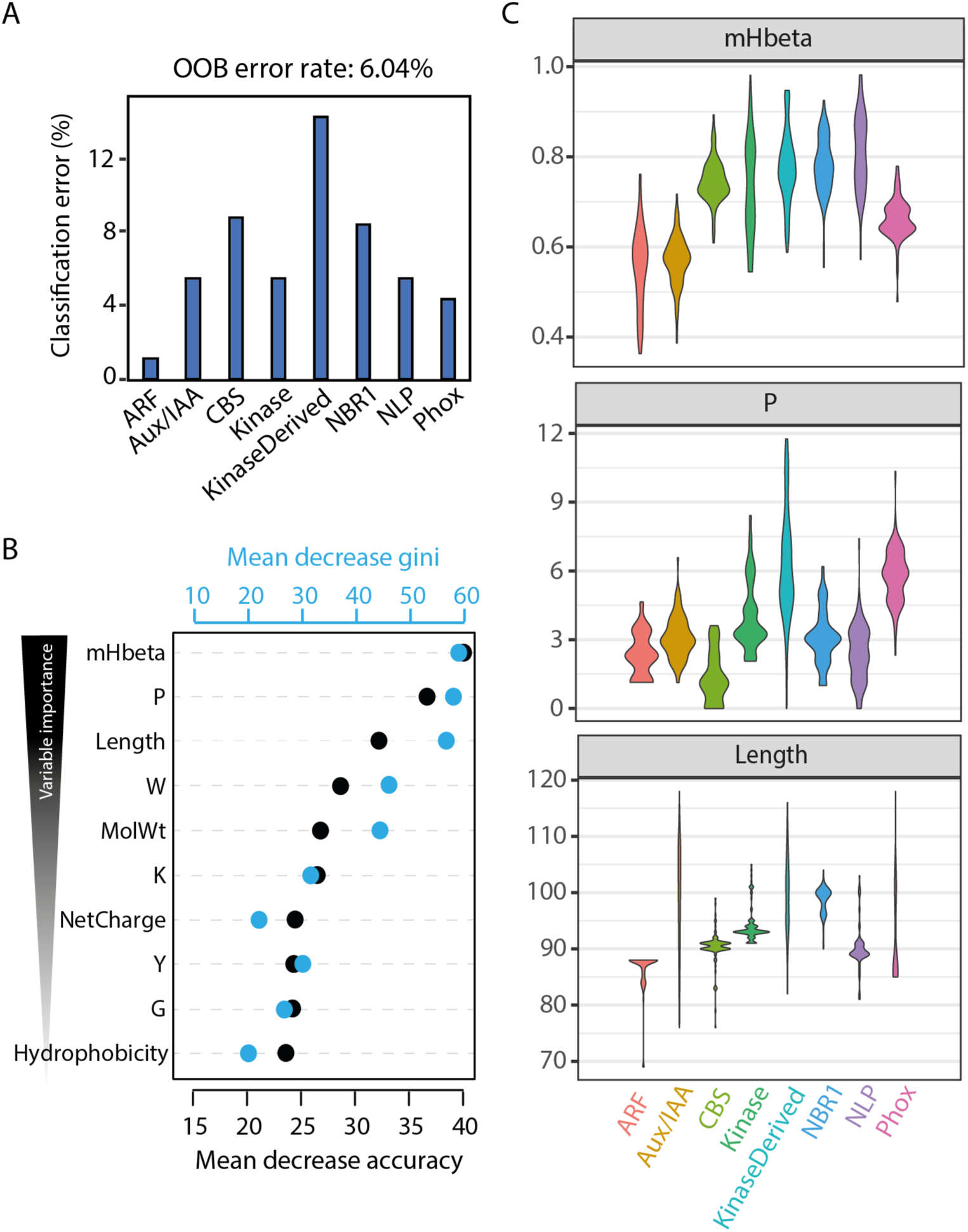
Random forest (RF) classification of the plant PB1 families. (A) Mean out-of-bag (OOB) error rate of 6% is reported for the classification of eight plant PB1 families with individual classification error % as shown in bar chart (B) Importance plots of 10 most important descriptors (variables). The predictive value of each variable was expressed as the mean decrease in accuracy (black dots with scale at bottom) and the mean decrease in Gini (blue dots at top), arranged from most important (top) to the less important (bottom) variables. (C) Violin plots showing the actual distribution of three most important variables across eight families. mHbeta has no units; Composition of Proline is indicated as percentage; and Length is the number of amino acids in the PB1 domain.

The importance of each variable is evaluated through the mean decrease in accuracy (MDA) and the mean decrease in Gini (MDG). Higher values of both MDA and MDG indicate the most important variables. In this case, the top 10 important variables are shown in Fig. 5b, with mHbeta being the most important variable to differentiate the different classes of PB1 domains. Hydrophobic moment of β-sheets, mHbeta, indicates the strength of periodicity in the hydrophobicity of the β-sheets, also indicating the formation of more β-sheets (Eisenberg 1984). The next most important variables are composition of Proline (P) and length of the PB1 domain (Fig. 5b). We further analysed how these three important variables differ between the gene families (Fig. 5c). mHbeta is low for ARFs and Aux/IAAs, slightly higher for Phox, but even higher for the rest of the gene families. On the other hand, the composition of Proline (P) is lowest in CBS, but shows a very broad distribution in the Kinase-derived family. However, the length of the PB1 domain is very constrained for majority of the families (>90 for NBR1 and <90 for ARFs), except Aux/IAA and Kinase-derived, and to a certain extent for Phox (Fig. 5c). To correlate the contribution of mHbeta to the β-sheets in the secondary structure, we have performed homology modelling of at least one randomly selected Arabidopsis orthologue and we indeed found that higher mHbeta represents secondary structures with more β-sheets (Fig. 6). For example, IAA17 shows ∼18% of the residues in β-sheets, whereas CBS36500 has ∼34%, correlating with lower and higher mHbeta values observed for Aux/IAA and CBS PB1s, respectively (Fig. 6; Additional file 1: Fig. S3). Taken together, these results clearly indicate that there is a difference between the gene families that can be explained from the mHbeta, the composition of Proline and by keeping the length unique/constrained for that respective family.

**Figure 6:**
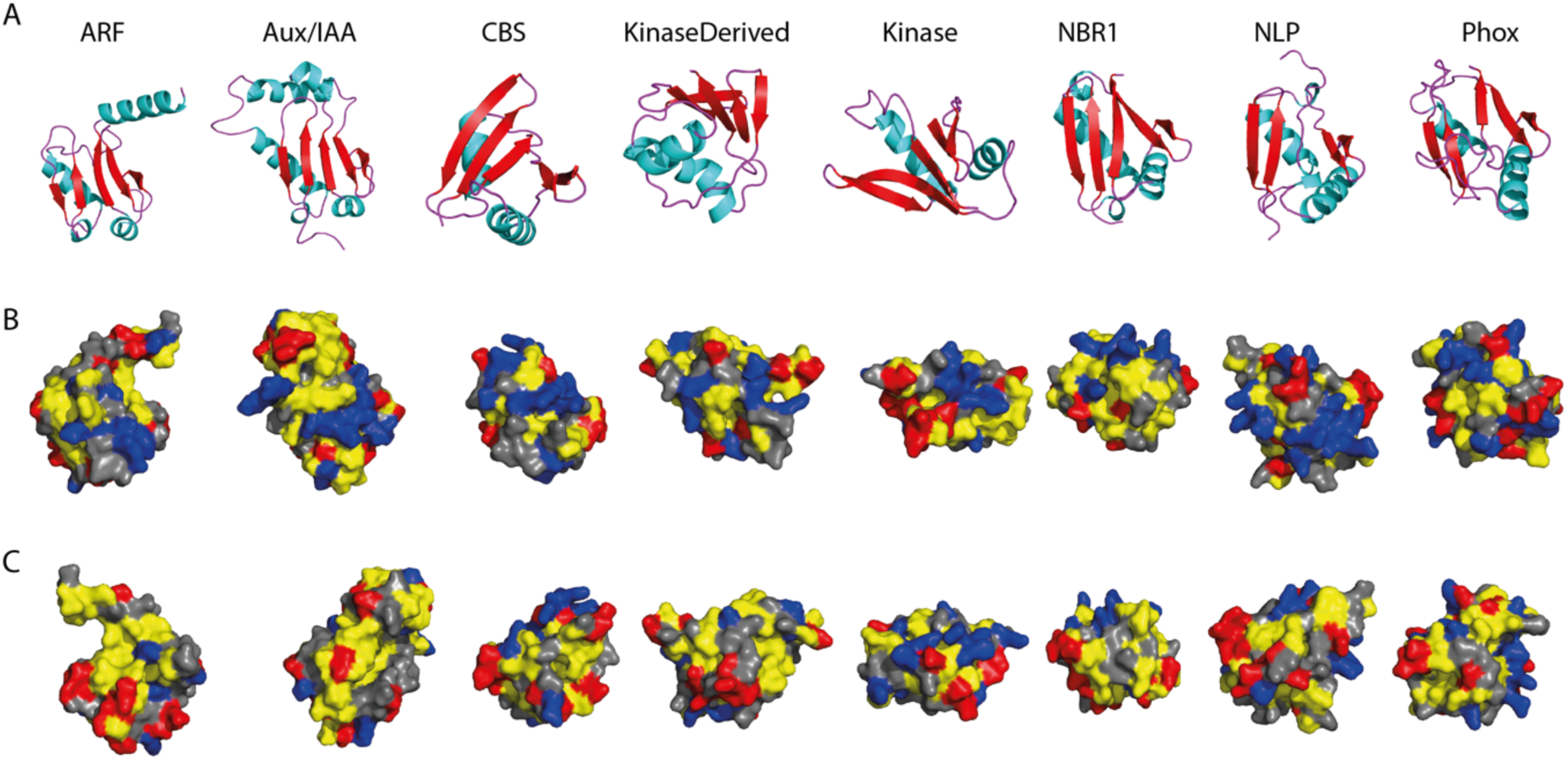
Representative (homology model) structures of one member from each family in Arabidopsis. The identifiers were: ARF5 (PDB: 4CHK), IAA17 (PDB: 2MUK), At2G36500 (CBS), At1G04700 (Kinase), At2G01190 (KinaseDerived), NBR1, NLP9 and Phox2. (A) Secondary structures shown in various colours: α-helices in ‘Cyan’; β-sheets in ‘red’ and turns in ‘purple’. Surface representation of the positive and negative faces shown in (B) and (C) respectively. Hydrophobic amino acids ‘AGVILFMP’ are in ‘Yellow’; Polar residues ‘NQTSCYW’ are in ‘Grey’; Positively charged ‘RKH’ are in ‘Blue’ and Negatively charged ‘DE’ are in ‘Red’.

## DISCUSSION

The PB1 domain is widespread in nature, throughout all the kingdoms in the eukaryotic tree of life. It is diversified to a great extent in organisms with complex body plans like animals and even more in land plants (Fig. 1). As a result, the human genome encodes 13 PB1 domain-containing proteins, whereas a simple model Angiosperm, *Arabidopsis thaliana*, encodes more than 80 PB1 domain copies grouped into eight families (Fig. 3). The proteins with a PB1 domain also feature various domains, representing a manifold association ranging from DNA/protein binding, catalytic function, scaffolding to membrane association (Fig. 1). However, the PB1 domain is (mostly) found at either terminus of the protein, preferably facilitating these to perform their native function, scaffolding or oligomerization, without the hindrance of other domains.

The evolutionary patterns of the PB1 domain showed that there are multiple families shared across multiple kingdoms (Fig. 2). Kinase and NBR1 are present in all the five kingdoms, while Phox is found in four kingdoms (except Chromista) with a similar domain architecture (Fig. 1). The phylogenetic placement of Kinase and Phox PB1 domains and their orthologs indicate the presence of two ancestral copies in LECA and presumably a third copy might be represented by the NBR1 and/or p62 group. It is known that orthologous proteins may perform similar functions by interacting with similar proteins across kingdoms. An example of common functionality of orthologous domains across multiple kingdoms is the recently studied DIX domain in plant cell polarity protein SOSEKI (van Dop et al., in revision). The DIX head-to-tail oligomerization domain is conserved across multiple kingdoms (e.g. DISHEVELLED in animals; DIX-like in protozoans), and functions in cell polarity by forming oligomers in plants and animals. In a similar way, the PB1 orthologs across multiple kingdoms may share common functionality in similar pathways. One such is the PB1 domain containing protein NBR1, which serves as an autophagy cargo receptor in both plants and animals, where homodimerization of the PB1 domain is also conserved as a part of its function [5,29]. Phox orthologues in animals (p67^Phox^) and fungi (NoxR) play a key role in NADPH oxidation pathway by interacting with the membrane associated proteins (gp91^Phox^ and NoxA/B respectively) as well as through PB1 domain with p40^Phox^ and Cdc24 respectively [18,30]. Similarly, in Arabidopsis, Phox4 was shown to interact with membrane-associated proteins KNOLLE, SYP22 and PEN1, which belong to the SNARE family [31]. However, it is unknown which PB1 domain protein interacts with Phox4 PB1 in plants. Discovering these unknown interactions may provide a link to the existence of an NADPH oxidation pathway in plants controlled by PB1-dependent interactions.

Apart from the proteins that are shared across multiple kingdoms, some are specific to each kingdom (Fig. 3). ARF, Aux/IAA and NLP families are specific to plants, whereas TFG, Par6, M2K5 and aPKC are specific to animals. ARFs and Aux/IAAs are involved the Nuclear Auxin Pathway, controlling transcriptional regulation of downstream targets with multiple functions in response to the phytohormone auxin (reviewed in Weijers and Wagner 2016). NLPs are master regulators of nitrate-inducible gene regulation in higher plants [23]. On the other hand, animal Par6 and aPKC PB1 domains are known to interact with each other playing a key role in cell polarity [32]. Thus, as far as can be inferred, these kingdom-specific PB1 domain-containing proteins appear to regulate processes that are specific to that kingdom.

It is interesting to see that the key interacting PB1 domains have also evolved in pairs. Some such pairs are: aPKC-M2K5 (Animalia), Bem1-Cdc24 (Yeast), ARF-Aux/IAA (Plantae) (Fig. 2). The interacting pairs (for example ARF-Aux/IAA) seem to maintain pairs of amino acids specific to those classes (Fig. 4). Hence, based on this ‘paired’ conservation pattern, it is enticing to speculate that the Kinase and Kinase-derived PB1’s might form interacting pairs (Fig. 4). Despite the overall poor sequence conservation, it is clear that PB1 domains are maintaining a flexible (global β-grasp fold) yet specific (local conserved residues) sequence context in each family may provide specificity in function. Adding to the complexity in specificity of each interaction, the PB1 domains can also undergo non-canonical interactions. In plants, PAL OF QUIRKY (POQ), a Kinase-derived PB1 domain, interacts with QUIRKY [24]. The PB1 domain of NLP interacts with HQ domain of TCP20 [22]. In animals, the M2K5 PB1 interacts with ERK5, among many others [10]. However, the structural and/or mechanistic basis of any of these interactions is currently unknown.

In various kingdoms, the PB1 domain-containing proteins have expanded to various complexities/copies. For example, NBR1 in plants is (mostly) a single copy gene, where ARFs and Aux/IAAs are represented by large gene families with more than 20 copies (Additional file 1: Table S2). This clearly shows varying duplication rates in different gene families. However, whether it is a single- or a multi-copy gene family, there is hardly any conservation in the PB1 domain among the members of the same gene family outside of key residues: Lysine in β1, Tyrosine in β3 and the OPCA motif (Fig. 4). Despite their low conservation, all the PB1 domain families identified in plants can potentially form a β-grasp ubiquitin fold (Fig. 6). Thus, for the PB1 domain it is evident that sequence conservation seems to be a less important factor than maintaining the overall β-grasp structure itself.

This poor sequence conservation is never a bottleneck to identify the most important features, as there are efficient machine learning based classification programs like Random Forests (RF). RF has been very successful in classification with highly correlated variables at low error rate [33]. The classification error rate is as low as 2% (in ARFs), but up to 14% in Kinase-derived, which could be due to the broader distribution of all three most important variables. This clearly defines that the more specific the variables are, the lower the error rate is. RF also provides the relative importance of each variable with the precision. Hydrophobic moment of β-sheets, mHbeta, the most important variable in our case, is low for Aux/IAA but high for CBS, correlating with the increased β-sheets in CBS (Fig. 6). How this increase of β-sheets could bring a change in function needs to be elucidated. Another interesting observation is that there is a clear difference between some variables being very constrained for each family. For example, the length of the PB1 domain is always above 90 amino acids in the NBR1 family, where as it is always below 90 for ARFs. Hence, it is evident that PB1 domains are constrained in different ways to maintain the uniqueness of that family. Moreover, using more (specific) parameters in future, one should be able to distinguish PB1 domains to a much broader extent, even across multiple kingdoms, and including homo- and heterotypic interactions.

Apart from DIX and PB1 domains, the SAM domain also undergoes head-to-tail oligomerization, but this domain is structurally different from both others [34]. It is unclear why the PB1 domains are much more widespread and preferred for (head-to-tail) oligomerization, compared to DIX or SAM domains. The latter are only limited to few families and few members in each family. One reason could be that, as discussed above, the PB1 domain is highly flexible in nature contributed by a wide range of (non)-canonical interactions and its diverse evolutionary patterns.

## MATERIALS AND METHODS

### Search for PB1 Domains in Animalia, Protozoa and Fungi

To study the PB1 domains in the kingdom Animalia, based on the literature, we first extracted all the PB1 domain sequences of Human proteome from the UniProt database (https://www.uniprot.org/proteomes/). To find other PB1 domains, we then used ten proteomes from the kingdom across various phyla (see Additional file 4). A protein database has been created with all these proteomes and queried this database with the PB1 domain sequences from already known plant (Arabidopsis and Marchantia) and animal (Human) species. BLASTP module in NCBI BLAST 2.7.1+ [35] was employed for this search and InterPro domain database v5.30-69.0 (https://www.ebi.ac.uk/interpro/) was used for domain identification in the BLAST hits. All the sequences that have a PB1 domain have been used for further phylogenetic analysis.

A similar procedure was used to obtain the PB1 sequences from Protozoa and Fungi. However, the proteomes of twelve fungi across multiple phyla have been obtained from MycoCosm database at JGI (see Additional file 4; https://mycocosm.jgi.doe.gov). For the Protozoa, we have used the six reference proteomes from UniProt (see Additional file 4).

### Identification of the PB1 domains in Chromista and Plantae

To identify the PB1 domains in the kingdom Plantae, we employed a large transcriptome resource, 1000 plant transcriptomes (OneKP) database [28,36]. Out of nearly 1300 transcriptomes in the database, we have used 485 transcriptomes in this study, covering all the phyla in the kingdom Plantae. We have adapted a protocol that was developed earlier [26]. In brief, the query PB1 sequences from Arabidopsis were searched against each transcriptome, where the resulting scaffold hits were translated using TransDecoder (v2.0.1; https://transdecoder.github.io). All these translated sequences were checked for the presence of PB1 domains using InterProScan [37] and only those protein sequences with a PB1 domain identified were used for further analysis. In a similar way, for Chromista, we have employed another transcriptome dataset, Marine Micro Eukaryote Transcriptome Sequencing Project (MMETSP) database [38]. We have used all the available transcriptomes and adapted a similar protocol as mentioned above (see Additional file 4).

### Phylogeny construction and visualization

Using all the PB1 sequences that were identified in all the five kingdoms of eukaryotes, we performed the phylogenetic analysis (see Additional file 5). The protein sequences were aligned with MAFFT G-INS-i algorithm using default parameters (v7; [39]). Alignment was cleaned up further, where the positions with more than 20% gaps were removed with trimAl, prior to phylogeny construction [40]. ModelFinder (accessed through IQtree) indicated ‘LG’ as the best model of evolution, of all the 462 models tested [41]. Further, the Maximum Likelihood (ML) method, employed in the IQtree program was used for the phylogenetic tree construction, with 1000 rapid bootstrap replicates and tree branches tested by SH-aLRT method [42]. The resulting tree was manually curated further for some misplaced taxa. In a similar way, another phylogenetic tree was generated using the PB1 sequences only from three kingdoms (Animalia, Fungi and Plantae). An unrooted version of both the trees were visualized in Additional file 2 and Additional file 3.

### Alignment of the plant PB1 domains

To understand the PB1 domains in the plant kingdom further, we have taken the PB1 sequences of all the eight families from four species (Marchantia, Physcomitrella, Amborella and Arabidopsis), aligned them using ClustalOmega [43]. After the alignment, the domains from each family were separated and a sequence logo was generated using All the gene identifiers from these four species are available in Additional file 1: Table S2. LogOddsLogo server was used for logo generation, with the colour codes for specific amino acids (https://www.ncbi.nlm.nih.gov/CBBresearch/Yu/logoddslogo/proteins.cgi). Amino acid groups ‘PAGFLIMV’, ‘KRH’ and ‘DE’ were shown in purple, blue and red colours respectively. All other amino acids were shown in black colour.

### Random forest (RF) based plant PB1 classification

The Random Forest (RF) method was used to identify key amino acid descriptors to differentiate and classify each of the PB1 domains into eight plant PB1 families [33]. To make this classification an extensive one, we extracted all the PB1 domains from Plaza Monocots database v4.5, that includes species from all the major phyla in Embryophytes [44]. Since the size of each family is different, and to make the analysis uniform and comparable, we have extracted 100 PB1 sequences randomly for each gene family (except 78 for NBR1 as it is a single copy gene). We have used 28 amino acid descriptors (variables) calculated either with ‘protr’ or ‘peptides’ R packages [45,46]. Among these, 20 variables correspond to the composition of 20 amino acids, and the remaining eight correspond to the general parameters such as length, molecular weight, hydrophobicity, net charge, isoelectric point (pI), aliphatic index, hydrophobic moment of alpha and beta sheets. We used ‘RandomForest’ R package to build a maximum of 500 decision trees with 5 variables being tried at each step (www.r-project.org; [47]). Confusion matrix and variable importance plots showing mean decrease in accuracy and gini were obtained. Descriptive plots and other graphs shown were obtained using ‘ggplot2’ R package. Additional file 6 provides the complete R script that has been used for the RF analysis.

### Homology modelling

Homology modelling for the eight selected members in Arabidopsis, one each from each plant PB1 family were performed using Phyre2 webserver ‘normal’ mode [48]. The identifiers of the PB1 sequences used were: ARF5 (PDB: 4CHK), IAA17 (PDB: 2MUK), At2G36500 (CBS), At1G04700 (Kinase), At2G01190 (Kinase-derived), NBR1, NLP9 and Phox2. Obtained homology models were visualized in PyMol software (Schrodinger Inc., USA).

## DECLARATIONS

### Ethics approval and consent to participate

Not applicable

### Consent for publication

Not applicable

### Availability of data and materials

All data generated or analyzed during this study are included in this published article and its supplementary information files.

### Authors’ contributions

S.K.M. and D.W. conceived the research. S.K.M. performed analysis. D.W supervised the research and acquired funding. S.K.M. and D.W. discussed and interpreted the results and wrote the manuscript.

## Acknowledgements

The authors would like to thank the 1000 plant transcriptomes (OneKP) and Marine Micro Eukaryotic Transcriptome Sequencing Project (MMETSP) consortiums for providing essential data resources for the scientific community, and Francois Parcy and Mariann Bienz for their comments on the manuscript.

## ADDITIONAL FILES

**Additional file 1:** Supplementary figures and Supplementary tables

**Additional file 2:** Full phylogenetic tree based on species from three kingdoms, related to Fig. 2

**Additional file 3:** Full phylogenetic tree based on species from three kingdoms, related to Additional file 1: Fig. S2

**Additional file 4:** List of species used for the phylogenetic tree construction in all the five kingdoms

**Additional file 5:** FASTA file of all the PB1 domain sequences used for phylogenetic tree construction

**Additional file 6:** R script used for RandomForest classification

