## Additional File 5 for "Deep Evolutionary History of the Phox and Bem1 (PB1) Domain Across Eukaryotes"

```
>GA_AKCR_2002629_Parachlorella_kessleri.p1|229_317
VVATSIPVEVEARPSAKAQPLGELPLSVESSYGELLTAIQAAFGERLPRKAGMRLVYQDADGDWLLLL
PDVPWQLFAGSVRRLLVTYK
>GA_ALZF_2020169_Halochlorococcum_marinum.p1|284_378
SRPTSQVVECSIEDAAGKEQATRARSLDLAGFDSLRHLWMGLRAMFESQLPQLADAKLVYQDADGDWL
KYTPDKPWSSFVTSAHRLSVRPADQP
>CA_APTP_2015690_Ishige_okamurai.p1|1_84
HLLKLRYDGVLRITEIDAPRFEVLVKAVETAYGIDNIEHYKFTYLDSDGDQVSFDTDAELELALRTTP
APLRISVSAKSPLSS
>CA_ASZK_2000638_Punctaria_latifolia.p1|39_111
TKKKIKLKLEFNGSKRVAAVYEGYTFARLHQRLCEDFGFDVFLEYEDSDGDKVMLNCENDLTELLENE
QGTV
>GA_AZZW_2004419_Chlorokybus_atmophyticus.p1|921_980
QTPLLTFQSLVNEVVKLFRLNTAQQLKLKYFDGEEWVLITNDKDLHEASRVAVKANANS
>GA_AZZW_2021372_Chlorokybus_atmophyticus.p1|458_554
DVMASERFTYKIEDSVGRVHRFQAAANDYEEVVGAVTQRLEVSEDEVRRHLKYVDDEGDEVSISCDAD
LTAAVALARSLSSKILRLQVFKKPTRTA
>GA_AZZW_2021616_Chlorokybus_atmophyticus.p1|666_763
TMHEQGFCKVFKNGSGVGRLVDLKRITSYTQLEDEVAQLFKLGKNEYRDSEGRLHMVYEDRDGDTMLI
EGGPAPWSMFLSSAQRIHLLPPENSMKAD
>GA_AZZW_2021745_Chlorokybus_atmophyticus.p1|632_708
KDSTVPRVKCFVQGTPVGVSIDLSRFNSTEKLKDTLLATTGGTNVVYQDADDDMILLGDESWEYFLKT
VKKMYIRT
>GA_BAZF_2072625_Chaetopeltis_orbicularis.p1|44_126
TNLWKYDGGESHNFSMPMNVTFQEFLQQVGKRWPTVGLRYADARDNPDPSELISLQTEDDYEEMKDEY
RRAVDLGLQSKNIP
>GA_BFIK_2004609_Entransia_fimbriat.p1|311_407
PRDGDGKLSYHGGETRNMTVARDIKYEELIFKFCEXGTDLTIKYQLPNWGLDTLVSVTNDDDVANMMD
EYDQLEARGARDGSQRLRLFLFSAEEQQ
>CA_BOGT_2014263_Mallomonas_sp..p1|2_95
DSSTLAIKLTYNGIIKRISFRICEFSFENLSQTTFRLFPSICSSRILFRYTDADNDEITLSTDAELDD
ALSFFQCMTKLTCVKFVVVESAVLF
>GA_BTFM_2005445_Monomastix_opisthostigma.p1|27_111
RMASTVLFKVTHATDTRRLTVGSDSTDFAEFCNKISALFGVAPGQLVFTYTDADGDTVTLATQSDMDE
LMLQQLNPVRLTMRLS
>RA_CKXF_2002994_Gymnogongrus_ftabelliformis.p1|1_89
SSLSFKASYKNSKRRFAVESTTSFADFQARIASMFSLTPPFTIAYEDDEKDIVSVSSDAELHELFCIA
STAEISLLRLHIYDSEEELP
>RA_CKXF_2004893_Gymnogongrus_ftabelliformis.p1|449_540
NIDTLIGVKVEFRGDIRRLRIGVKSSFQSFASELCDLFELQGPLTIKYRDEENDFVTIANERDMKEMI
YMVKEHRLVPLRVQLEERPVQNE
>CA_DBYD_2059002_Synura_petersenii.p1|1_88
ISIKLSYKGEIRRAAISKSSFNYESLNNFARRLFPLLNEVRKFSFSWTDDDGDKITMSSNEELLEAFR
IGEGESTGLLRFEVSLNEN
>GA_DFDS_2041262_Desmidium_aptogonum.p1|1_94
DNLQVIKVKFSGSMCRISARKSTDENVISLTALETRLRASFKLPTAAYLALTYKDSDGDVITISSDQD
IVDALVYQELNPLRIDLDVQLPEGV
>GA_DUMA_2036503_Tetraselmis_cordiformis.p1|2_96
RTRRLPVFKVSLGDDTRLWNPGEALRYDELHCKVIEMFCYDLEQSSTFLIRYTDADGDKITVASDGDL
NLLLRSFRTYESQVRLTVEKRKKAKE
>GA_FMRU_3048952_Zygnema_sp._2_samples_combined.p1|1_88
ATHLVKIKLDDDLRIMEIPSPPRFEGLVKAVMEAYAVPPRKEKQLSFTYRDNDGDEVRFDTDAELDLA
IRTSSVPLRIMVQKNARSR
>GA_FMVB_2002844_Scherffelia_dubia.p1|1_90
```

AGRLLFKLQCGGDLRRWTLEGPPEALTWTALHEKLRELFGDAADEAALQYTDADGDRVTLGCQSDVVE
LLRQRLEGVYLTVAVKDKAKD
>GA_FMVB_2004451_Scherffelia_dubia.p1|45_139
DGLYKYIGGESYLESVPRSWRYHELMFRLTEKVQQGVSVKYQVPGEELDPHALISVNDDNDLQEMFDE
YYRGLHRPGTPVKTFRLRVFLFKAAE
>GA_FMVB_2038112_Scherffelia_dubia.p1|161_258
YGKVYKGGETRLVTLELDQNTKRTQLIRQLMGVGSSNDVSALEAAQVKYELPSEPGIFVAVTEDEDVQ
NMFEEWQSAQAPNGAAKKLRLFIEKVAPE
>CA_FOMH_2016910_Sargassum_integerrimum.p1|447_533
GGGSVSIKVEDEAGRVHKISASAGSLGSVLQAVAERTGIPSDAVRLTYKDESGDIIVLASDDTLRVAV
DLAKASNSRSLKLAASRM
>GA_FPCO_2028016_Interfilum_paradoxum.p1|382_472
LQTSFTFKIEDRQGRNHRFTCGCQSMAELVVALSTRLDLPPAAIPPISYIDDEGDRVLLSGDGDLSSA
VNVARSAGLKNLRLYLDFSSGG
>GA_FPCO_2028362_Interfilum_paradoxum.p1|499_591
EMDVGLTVKATLGQDTARFKLSPEMGWQEVVSEVAKRLKVEPSTVKLKYLDDEQEWMLLSTDQDLAEC
IDIVRSTGNSIIKLMVTGGEGVGQ
>GA_FPCO_2029246_Interfilum_paradoxum.p1|1_89
AFIVKATYGGVMKRLTYAEEGPTAFQNLVKTLKESFSIPEAATLRIQYMDQDKDKVLLSDKKDLFDAV
TVQKLNPLYLEVTVISEDQP
>GA_FPCO_2029587_Interfilum_paradoxum.p1|294_377
LERPAIPLKLVYQGHDIRKTELPANGGLRTLREVVQKRYPGSKNVLIKFQDEDADWITITSNEELRGA
LELAGYAGKSAGAEN
>GA_FPCO_2030410_Interfilum_paradoxum.p1|339_437
GEIKFTDGKLRYVGGETRIFTVSRDITYSELMFKLTEYYSEALSLKYQLPDMDLDTLLSVSSDEDLAS
MMEEYDEIERKAAEGKAQRLRLFLFSAEDK
>GA_FQLP_2009864_Klebsormidium_subtile.p1|709_785
EEEQRSQVKCFVEESPYGVVVDLSQYKSTQQLKNALLDATRGRLVVYQDKDGDMILLGDEEWSFFLKT
VSRIFIRK
>GA_GGWH_2009874_Onychonema_laeve.p1|1326_1402
VRGESQRVKCFVEGSPFGVSVDLSRFHSTQELREALLSVTEGTSVVYQDRDGDMILLGDESWDFFLKS
VKRMYIRR
>GA_GYRP_2005594_Euastrum_affine.p1|393_484
PDQPIMFKMEDKKGRVHRFHCKSDSLTELVCAIASRLGGDFDPDNPPSIMYDDEEGDKVVISVDDDLS
AAVKFAKASNMKALKLSLDYHGD
>GA_HAOX_2000118_Spirogyra_sp..p1|141_233
VKPFNVSTKVIKSGAIGRMINLSKFSGYEELKVEIGTMFKLSAHDFEEEWHLTYVDHDGDILLLGDAS
WQEFLSAVRAIKVLSSNEVSLLSA
>GA_HAOX_2002346_Spirogyra_sp..p1|1_91
EGLQVIKIEHNGTLRRFTYPEKESISFSSLDSKIRELFKFNSETYFSLSYSDSDNDIVVMSQDEELKD
AISNQKLNPLRIKVAKSLKAAD
>CA_HFIK_2069821_Sargassum_vachellianum.p1|1040_1122
FVYKVNDEAGNLRKFRASASNLERVRQAVANQIRVPMEELVLKVEDDDGDEILLTGDDILHEAVSLAR
TSSSGALKILASRK
>GA_HVNO_2008550_Tetraselmis_chui.p1|10_97
SPLCAVPFKFTLRKDTRRWTPSTRPTHGELLARVRMTYELPEETTINLKYTDADGDQVTLASDSDVQE
LFRQSLPVIRVAVTAPEWA
>GA_HYHN_2002361_Prasinoderma_coloniale.p1|755_852
GDSGDIVIKAKYGGETVRVRMPAAQAALATVIARVATAIGCAPPAALKMRYLDREGDLIRLETQQDFD
ELVAATLEARPAEVPQRPLTVRLDVVDAL
>RA_IEHF_2001841_Dumontia_simplex.p1|480_572
PLASFKFKDINGEFRRIKVPMTLEPGDFDQFVVDVRRRFGGSTGVGPIKIKYVDEDRDEVLISNDEDL
ASCFEDFTGIRQGTIQLKVYEVER
>GA_IJMT_2016123_Aphanochaete_repens.p1|2_92

TMSLVVKIQFGQDTRRVSVSSGLPNFQALADLLLRLFPNLNLHDYAIRYTDPENDLITVSSDIELREA
IDLCTSERAPLRLLITPKDQIS
>GA_IJMT_2089881_Aphanochaete_repens.p1|159_237
KALCGDEVRTLKTKAGVRFEELSTEIANMFSKKRFRLKYKDSDGDEVTISRDEDLAEAIKSLPSGVRT
LKIFVYERRE
>GA_IJMT_2091548_Aphanochaete_repens.p1|322_407
HVRVFKCFLDDDVRLLTVETKQLSVQYLIAQVTEEYNRERLVIKFMDEDGDLIRITKQQDLDYFLALY
GHSKALKLFVFEDKGLV
>GA_IRYH_2006286_Heterochlamydomonas_inaequalis.p1|656_742
LVECLMEVAGHALSQMQPQIVDLAKIRSFKDLWSHLAELFRDDMPDKMDAKLIYLDDEGDWIMVTPEE
HWSLFVASALKILVTSRC
>CA_IRZA_2008892_Proteomonas_sulcata.p1|859_934
DALQEKRIYKLGDNFTYGELCDKVMELWKGPLTIKYEDEDREWTTMVTNEDLDEAKASCYNHEVRKMT
LRVMGEP
>CA_IRZA_2009647_Proteomonas_sulcata.p1|365_438
EGSQEKRMFKLPSDFTYKNLYEKLLTIYSSRFSISFEDEDQEWIMLTNDEDVETARESGSQSYKQKLI
LKVSF
>CA_IRZA_2010067_Proteomonas_sulcata.p1|155_258
SNVFEYKVKIEFDRTKRVVSASIKWEWSEFLAHISKALELDVDHFEGPEPQLHLHYIDKEGDVIVVTN
EEGYEEMTTQYTTSAPWRDLPPLRIVVLQGDVDPI
>GA_ISHC_2045732_Staurastrum_sebaldi.p1|341_436
ALEPGLDGGVEYRGGEKRMIGFPRSMAYQALCDKLAQVFGQVPRVKYQLMHDRDLVISVSNDEDVENM
LAQCQCPGAGQSYCRELMLVLEFQEQQ
>RA_JEBK_2023167_Eucheuma_denticulatum.p1|346_430
DEGKMTVKVEKDGDLRRLRIDTDWTFEGLREELVGMYGLVGEFSIRYRDEEGDFVTVASEKDMKELFQ
LVKEHKLVPLRMKVVM
>RA_JJZR_2000281_Rhodochaete_parvula.p1|494_588
TEVVFKLKDLNDDVRRIPMTFQSHDFNESFDQFLSRIRAGFGVSSKQTIKVKYIDDEGDEVVITNDTD
LAECITVVSEMKSKTVQLRVSVVPDP
>GL_JKHA_2011089_Cyanoptyche_gloeocystis.p1|5_101
SVPQGLRLKCHIGGHFEETPHGSRRYAGGEIRVITLPPDISYSELMFKLCEDYGGVTLKYELGDGDLV
TVRSQEDMLELAREYRYLLLKSQKRFLE
>GL_JKHA_2019758_Cyanoptyche_gloeocystis.p1|1_85
SFKSNYKGALRRFALTPPTWETLQGTLRHLYGAPAYASVTVQYKDEDGDDIFIGSDMELEEAIRIGAK
KAVLPINVEFNPAIEN
>GL_JKHA_2031828_Cyanoptyche_gloeocystis.p2|455_548
APGVEFTVKASLGDEIARFKLDSHRNVVDLYREVRLAFSHVIPRECDIAVKYKDDDGDMVGLLREEDL
RECLALAHNQGTKRIAISVNIVSLR
>GA_JOJQ_2007390_Cylindrocystis_cushleckae.p1|6_94
NKNLVVKIKYGNTLRRLTVPRSTELSYSWLEGKVRSLFQLPEDAPLSVTYVDNEQDVVTMGDDMELKD
AVSLQGLNPLRLTVSVFNSE
>GA_JOJQ_2038013_Cylindrocystis_cushleckae.p1|191_285
GTASLQYIKVNKYPSSITRKLEVSRLVNYDELRSQITNLFGLGNEDPLENYNVTYKDKDGDIILVGDE
PWGDFLLNVRSIELFTNLPEGEDGKG
>CA_JQFK_2080261_Nannochloropsis_oculata.p1|152_241
NRQITVPVKLTMDSDTRAFQLAPGITYSELMEHARQLFPNAGPFVLKVLDKEGDLVTIASRADINRAI
QESIDAAGKSGRIQQGSLQAI
>GA_JTIG_2004837_Bryopsis_plumosa.p1|168_241
LYFSAKCTLDSETRVIHLSHNTSYAELLDVLKAKFPTSGPFVAKYLDKEGDLVTITEQSDISNAIAEV
LAAYE
>GA_JWGT_2063704_Volvox_aureus_M1028.p1|37_131
PRQWRYVGGEVYNESFPLDSKYAEVMKRLNDKFGDTVSFKYLCPGDDIDPDNLVQVQGDDDLQEMCDE
YHNALQKSSTPVKTVRIKVFVFRAVI
>GA_KADG_2047154_Ignatius_tetrasporus.p1|104_189

VNGTVVYKDGEIRLTTFRPPYTWENLAAVLPDYCTNTEAIKATWQLKFALPDDESTLVDVKTDADVYN
MVEEVMDYKQTKPTYKV
>GA_KEYW_2002865_Gonatozygon_kinahanii.p1|331_409
MVVLKLLMNGHDLRVGQMPITYGFKQLQDISKTTFSLSPESSILLKYVDDDGDLITMTSRSDIQAAIR
IAAAIAPNPS
>GA_KMNX_2004693_Nucleotaenium_eifelense.p1|3_91
RLHTVVKVKYNGSLRRFNFDETKDCSFALFSKKIRNIFTIPVEVAIVVTYIDDDDDQITMACDDDLVD
AISVQSLNPLRVTVSHASLV
>GA_KYIO_2003156_Mesostigma_viride.p1|333_437
EHEGEPRFVFKMERPASSAVAMSPQRSPSENPNILKFFSSSSSLRELRLNVIKRLGWSDPEALEGVLK
YMDEDGDAVTIGCDADVQAAVRGARLHGAKFVMLAL
>CA_LIRF_2015667_Dictyopteris_undulata.p1|758_850
GWDEKFVYKVSDDSGHTYKFKASAEKLESVLVAVAEKLKKPKESLLLKYADDDGDQIVLSGDDSLHEA
IDMARASSAPALKLVASLKLGTVE
>RA_LJPN_2003218_Gracilaria_blodgettii.p1|395_487
PVASFKFRDINNEYRRIKVPMNLGPGAYDQFVLDVRRRFAGSNSVGAIKIKYLDEDKDEILISNDEDL
ESCFEDYLEMRSKTILLRVYEAEH
>GA_MCHJ_2016284_Micrasterias_fimbriata.p1|625_718
QPGGLVTVKATFGIDTVRFKVPLNGMAFRTVEEEVASRVKLRPGEFSLKYLDEDEEWVVLASEQDMME
CIDITRAQGNNVIKLVVKAEIDASH
>GA_MCHJ_2017763_Micrasterias_fimbriata.p2|10_100
SMEDGRVRVRGGETRMLALPRSITFQELATKLMREYGVLPSIKYQLRSEEDALITVASDEDVENMLEE
CEREAAAHRKKLRLFLERPQVA
>GA_MCPK_2014984_Bathycoccus_prasinos.p1|903_975
VPQLVVKFRVGADTIRFRLLHKWSVQMLLDRIRKLIVFPDSAKLKYMDDEDDMCILQSEHDLDECIKF
TEHR
>CA_MJMQ_2004620_Hemiselmis_virescens.p1|9_88
DEEGEETRWVFRLDTDNFHFSDIYTRLLEKYGERLVITYQDEDEQWISLQNDEDLAEAKKSGKERAKH
KLSITVAQEVP
>GA_MMKU_2001882_Nephroselmis_olivace.p1|1_89
AGTVLKIFFQDAIRRMTVDLDDVETFREFQDLVAEVLALDDASNYRFSYIDADGDTIALVNDCDVDEL
TTQGLKTIRVNVVRAAEPVY
>GA_MMKU_2049414_Nephroselmis_olivace.p1|189_263
MPPLSVKFVKGDDIRLGQLPATVNYVELLSAAQSKYPGAGSFSLKYKDRDGDLITITSRADIRTAGVE
AQAGVD
>GA_MWXT_2012711_Chara_vulgaris.p1|153_241
TIEEHFFCKIFRGGELVGRAVDLGKFSNYDQLCAELSVMYGIDRHELQKNMAYRDSEGHWVLVGDEPY
RHFAGTARKAMIMSTGGDHP
>GA_NBYP_2052036_Mesotaenium_kramstei.p1|212_301
DHRASPRIKVYLPGSIGRKLDIRRLTGYADLKHEVALLFGLLEDELMSKYRVAYMDLEGDVLLVGDEP
WGEFLMNVKNLQVVPMQEIEN
>GA_NBYP_2055233_Mesotaenium_kramstei.p1|399_491
APAGTFQFKMEDKQGRLHRFNCGTESLTELSASIATRCASDWDHEHPPAIMYLDEDGDKVLLSNDEDL
TAAVQFARTSQSKSLKLHLGFTPM
>GA_NNHQ_2003350_Spirotaenia_minuta.p1|291_387
PPADWFVYKVQDANEGPIHRIHAPANDFSKLLQAIALRVGSASPARLRAAVRYIDDEGDEVSITCDSD
LAAAVLVAKQARSQVVRIKLVAGRSFGK
>GA_NNHQ_2007429_Spirotaenia_minuta.p1|1_88
ATTVVKVKFGEDLRRLTIPTSVDFNFVDSRVRELFGIPAAATLLLTYHDTDSDVVTLASDNDVADALR
VQRLNPLRLFAEVKGNDEA
>GA_OAEZ_2005814_Persursaria_percursa.p1|175_263
QSNNQFVCKVTMETETKILHLPFGVTYYTLQQAIKEKWTGLHNFKIFYQDRDQDWVLVTCARDVQKAQ
QDIISYAQRVLSHRQRQGLD
>GA_OFUE_2047682_Lobochlamys_segnis.p1|187_268

LANTTFAAKLSLGDETKLLHITATTSYAELLSAAQAKFPNAGPIVLRYVDKEGDLVTLTCRADLQQAI
GELLLANRNHISA
>GA_OFUE_2047741_Lobochlamys_segnis.p1|23_137
EMAGAIKLRLHYQGQFYKENGSWAYSGGEVFNESVPRSFKYSDLCRKLNDKFCDTVSFKYLAPGEDLD
PSNLIAVGDDDDLQELYDEYDTALRRPGTPLKTFRIKVYVFPACQF
>GL_POOW_2009423_Glaucocystis_cf._nostochinearum.p1|11_112
NEPMAIRLKCHVGGRFVVDGNTRRYEGGEXXXXXXXHFGITYSELMFKLAEEYGAVTLKYELGDGDLV
TVRSQEDVMELCHEYRDMRSVKRYLEVYLFEAM
>GL_POOW_2012277_Glaucocystis_cf._nostochinearum.p1|1_93
LTVKVCLGEDIRRFQAERTIRYSELDETLKQLYFPSCSPPFFVKYVDEDGDLVTVSSDNELEEAIRLA
IRSGEDESKAMLRLYIDVPTYSAS
>GL_PQED_2012992_Gloeochaete_wittrockiana.p1|88_170
GGQRRYEGGEIRLMTIPFNISYSELMFRLAEEYGAVTLKYEIGDNTGELVTVRSQEDVAELCSEYAEL
RRTSKLRYLDVFLF
>GL_PQED_2052415_Gloeochaete_wittrockiana.p1|351_426
VEALYEGGSFVFSTRFGQSFADLKSLLRHLLYVDASEEGDIAVKYKDEDSDWVWMVRENDYQLCQQLG
KEYMDNK
>GL_PQED_2053825_Gloeochaete_wittrockiana.p1|595_687
QNEVEYTMKVSFNDELARFKFGGKRSLNDLVHEVRAAFEHVIKPEQTMSLKYKDEEGDFISLVRDQDF
AECLQVARYTNNRCNIVVTLKQKS
>RA_PYDB_2000614_Sinotubimorpha_guangdongensis.p1|1_85
SQGVKATYGSTKRRIVIARDVSYGAFREQLIRLFGITNTDEGQPIPADDITFSYRDSDGDVITVSSEA
ELREMFRITNEPTLLM
>RA_PYDB_2025587_Sinotubimorpha_guangdongensis.p1|29_104
IMSVIIKATQGAVIRRLPFQAPETLESLRTQLSTHFTISADARLTYKDEDGDEITFSTDAELVDQLRT
AALNPVK
>GA_QPDY_2002632_Coleochaete_irregularis.p1|332_409
RRPPIPLKLVYEKGSDLRRATLPPRCGPRGLRDIVQQRFPSSKAVIIKFRDMEGDLVTITNAEEFQLA
LAGYVERPR
>GA_QPDY_2005537_Coleochaete_irregularis.p1|159_239
CSSGAEFCKVWWHQEPVGRTVDLSRFGSYQELFQTLAEMFHVDSFELHYQDKGGHWVPVTAVKPYTHF
ATTAKRARIVTI
>GA_RAWF_2006144_Uronema_belka.p1|31_134
GEFKVQDSAYKYDGGQSFMESVPRTCTYSELIHKLNEKFDHDLSIKYLAPGDVLDPQNLVSVQDDDDL
KELYREYFAWHKRNPKSEGTFRIRLFLFNAVEDED
>GA_RHVC_2038843_Dunaliella_salina.p1|165_273
GPNLQLTIKATLGNESKLVTAPLTVSYYELLQAVKAKYPDAGPFRLQWTDKDGDNVTVTSRQDIQTAL
SELFLAFQKQHGGAHAPRLLQQNGLPPLKLQVVPVEKQSQ
>GA_RPGL_2023221_Cylindrocystis_brebissonii_M2853.p1|297_394
AGAPPGSFAFKMEDRRGRVHRFTCGTESLTELTASVAARCGKDYDPNDPPAITYVDEDGDKVLLSSDA
DLAAAVHFARTSQTKNLRLHLDYDHTLAA
>GA_RPQV_2004229_Phymatodocis_nordstedtiana.p1|457_533
GDGESQRVKCFVEGSPFGVSVDLSRFHSTQELREALLSVTEGTSVVYQDRDGDMILLGDESWDFFLKS
VRRMYIRR
>GA_RPRU_2001832_Staurodesmus_omearii.p1|56_146
TMSNGKIVYKGGETRMMAFPRTISFHDLAAKLTHEYGRLPCIKYQLPNEEDALITVRNDEDLENMLEE
CDREATHSKKLRLLLEKADVGT
>GA_RQFE_2037907_Cosmocladium_cf._constrictum.p1|595_688
GGDPVVTVKATYGSDTVRFKVPLKGAAFRTIEDEVASRVKLRPGEFSLKYLDEDDEWVVLASEQDMVE
CVEISKAQGHSVIKLVVKVETGNSS
>CA_RWXW_2003827_Sargassum_horneri.p1|24_115
INMATTLVKIKFDEDLRMVEIPSQPCFSEFVKTVAEAYDLALGEAQSLTFNYKDTDGDEVRFDTDAEL
ELAIRSSPSPLRVTIQSSYARGK
>GA_SNOX_2006283_Planotaenium_ohtanii.p1|290_386

TVGPPLSFKMEDRKGRKHRFSMAVGSESMSELICAVASRLSSDFNPDNSPEILFEDEEGDRVVLSSDE
DLKSAMDFARAIKAKSLRLHFDFHESEP
>GA_SNOX_2007773_Planotaenium_ohtanii.p1|370_456
SNSEKLKIKLIYNGVDIRRAEFPEWRKRNACRFLREIASQTFRRSKHVLFKYTDEEGDLVTITGIEEL
RHALLLHLSLFQKTGSNN
>GA_TGNL_2000716_Picocystis_salinarum.p1|725_817
GEHSGISIKATYGNERLRFQYREGMNYESVLYEVSAYFKIAPQALRILYLDDQQDWITLARDADLQEC
IEVSAELTPAGRRVVKLTVQKLAE
>GA_TGNL_2002226_Picocystis_salinarum.p1|191_263
NPNLVKFRCSLEEDIRMLTVPAGITYHGLLKELRNLYIGHYPFCVKYKAKDASMVTITCKEDVAAAYA
VKQD
>GA_TNAW_2000454_Pyramimonas_parkeae.p1|1_89
GSTITVKSDLQGDKRRFQLDVRGGFDALRLAVETAYGECNLLMKYEDDEKDLVTLVSDDDLQEALRVL
HSMNLSSPLRLSVFNSAVAV
>GA_TPHT_2022039_Spirotaenia_sp..p1|226_323
GAQVDHVFSFKMEDLKGRVHRFSCGTESLTELVVAVAGRLGSDFDPDNPPSVLYEDDEKDKVLLNSDE
DLAAAVSFARTTGLKSLKLVFDFSEAPTE
>GA_TSBQ_2005923_Chlamydomonas_sp._M2762.p1|178_281
QYPYYITVKCNHNGETKLVHTQLLVSYADLYDLVKQKFPKAGPFLLRYTNAEGTTVTIASRLELQAAL
GEAIALFQKQAPSTPKGMAPQLPPLKVEVVACTEA
>RA_UGPM_2003717_Chondrus_crispus.p1|1_89
DQSIIMKVAFEGTIRRLPVTRDISFADFHQQLAERFSITAPFVTQYEDLDKDIITFDSDAELQDLFST
IEDMSKPLRISLFTLEEANE
>RA_UGPM_2024126_Chondrus_crispus.p1|54_141
IMAPLSVKACYKHTKRRFSLEPSSTFADFQAKLAAIFCIPAPQTILYKDDEDDLVAVSSDSELAELFA
IAASANITPLRVFLYDTAE
>RA_UGPM_2025075_Chondrus_crispus.p1|295_393
GDVALASFKFKDLNHEYRRIKMPMPPAPGAFDQFVVDIRRRFAGSANVGPIKIKYVDEDGDEVLLSND
EDLASCYDDHLESKNRTIHLRVYDTERPTS
>EZ_UNBZ_2019015_Euglena_sp..p1|76_166
DQQEIKLKIKCGQEVRVLTVMATTNYAHLLKTLTSDYGEIESMTYEDAEGDTLSIRSQHDLTQAFDYY
LEYLQDKGTALRVTVVKSQLAR
>EZ_UNBZ_2033230_Euglena_sp..p1|162_257
GEHMPMTLKCHMGDTAVLLSVARSITYADLQRVLTEEYKQDVCIQYHDFNGDLVTVASQWSLDTAKAQ
HEQRCKDLPAHHRVFDLYLSTPPEPVL
>GA_VALZ_2000486_Chlamydomonas_noctigama.p1|1_118
DKERVRLRLHWGGSFQSSSALPEWKYVGGEVFNESVPAESKYFDLCRKLNEKVGYTTSIKYQAPGEEL
SPNELISITGDDDLQELYDEYFQALLRPGTHVKTFRVKIYLFPALQLEL
>GA_VAZE_2052983_Cylindrocystis_sp..p1|354_436
NDGPQVRVKCFVEGSPYGVSVDLGRFNSTSQLRGALLTVTEGTSVVYQDKDGDMVLLGDESWEFFLHA
VRKMYIRRDDPPKP
>GA_VJDZ_2041089_Botryococcus_sudeticus.p1|180_282
NPVAYFNAKVQLGDELRLIHLDPRISYVTLMDQVRARFPEAPEHFVLKYVDKDSDLVTLASRMDVQAA
LAEAVSNMERKGGLGQLSVPTLRITVTPVASASD
>GA_VQBJ_2006004_Coleochaete_scutata.p1|1170_1259
SDGLITVKATYNQDTVRFKLSAEADYGNLRHDVAKRLQLVEELTHLKYKDDDDEWMLLTNDQDLQECV
ELVRKSGGSMIKLMVRYDGAP
>GA_VQBJ_2008956_Coleochaete_scutata.p1|2_97
LLKETLVVKVNVEGVSDELHRFTFGKGVTLCFTDLEKKICARFGLPVASSLRMTYKDEEDEKVVMTDD
DDLKDAINLQSLNPLRLYVVVKKDAVK
>GA_VQBJ_2009355_Coleochaete_scutata.p1|927_1003
SGQGGNRVKCFVEGSPYGVSVDLSRFGSTDELRLALLSATEGTTVVYQDAEDDMILLGDETWEYFLQK
VRKMFIRR
>CA_VYER_2086858_Sargassum_hemiphyllum.p1|750_825

EAGNLRKFKASASNLERVRQAVASQIRVPEEELVLKVEDDDGDQILLTGDDILHEAVSLARTSSSGAL
KILASRK
>GA_WCQU_2015062_Staurodesmus_convergens.p1|41_120
SMQVIKVKYEGKNCRISAAKSANADVITFDTLEDRVRASFKLSPSAALVFTYTDCDGDVINLASNQDI
VDALLYQQLNP
>GA_WDGV_2054867_Cosmarium_subtumidum.p1|387_480
GDDAPFTFKMEDRKGRVHRFSCASDSMTELVCAIANRLGGDFDPDNPPSIMYDDEDGDKVVISVDDDL
SAAVKFARASNMKGLKLTLDYHDDD
>GA_WSJO_2003854_Mesotaenium_braunii.p1|836_912
GAGESLRVKCFVEGSPFGVSVDLSRFNTTQELREALLSVTEGTSVVYQDKDSDMILLGDESWDFFLKS
VRRMYIRR
>GA_WSJO_2008687_Mesotaenium_braunii.p1|2_90
ESIIVKVTCNGSLRRISLAKANEPSFGALESKIRDVFKLAAIAPLTVTYLDEDADEVLVDDDAGLRDA
LVYQKLNPLRITVKVKDASA
>GA_WSJO_2039692_Mesotaenium_braunii.p1|40_134
SSRQRAFTKVYKRGHLTRSLSVSALGGYGQLREAVERLFGLPKGSLADAESGWTLAYEDLNGVSLSVG
DDAWGHWLTRVKHLRVWSQLEALPVG
>GA_XIVI_2011751_Cymbomonas_sp..p1|25_109
EWTRRTLKIRYGEDLRRHSSSEGDLGYPGLLKLITELFGVTRFVLKYRDEDGEEVTIAREEDWRECER
CPLRPIRLFVRSCEEQ
>GA_XIVI_2095815_Cymbomonas_sp..p1|6_103
ASAEPTSCKVVFGSHGSGSQAPHDVRRFKLQPGGGFAYLEDKLKAAYPEAGELTIKYVDDEGDYVTIG
DDEDYEVALSLCTESIFRVFVSESPAMPQ
>GA_XIVI_2096822_Cymbomonas_sp..p1|896_991
DGLLDIVVKARYGKDTFRLKFSLQGQKADYQPLLNKLAARLQLPADEIKLEYTDDQDDMIILGCDEDL
LSCLQLARATNKSCTLLIKLVVSRIIR
>GA_XMCL_2000747_Prasinococcus_capsulatus.p1|827_896
HATTGDVVKFKFDSYQTVEKMRNKVAVAFGMDAQQVRFRYKDEEGDSCLIEDEDDIQEFVGNLTRGLP
N
>GA_XMCL_2016423_Prasinococcus_capsulatus.p1|142_236
EKGWTYDGGETRIIEVHLSDSLDDVVKKIGLSPPGEELRYELPGQCDGPAPLVTLNSEDDLRFMFDEF
EDALASVPEGKRRSLKLHLFVVEKSM
>GA_YOXI_2048426_Cylindrocystis_brebissonii_M2213.p1|161_229
SSGNSAFIKVHKEGCISRKICLTNFSNYEELNQGVASLFVEEGLSKATEGYRLTYVDLEGDVVLVGDE
>GA_YOXI_2056567_Cylindrocystis_brebissonii_M2213.p1|1_89
AQVIKVKYQNVLRRVLISEPEQLSFRGLKERIAGIFGLPSDANLKITYTDCDNDVVALVDDTDVRDAI
MLQRLNPLRLVVEEVLLSTF
>CA_YRMA_2101013_Sargassum_thunbergii.p2|11_95
DGEVKGEIRRFQVDLPSGSTELYPCLSSKVADIFALAPNSFRLYWQDADGDFVTFTSTMELMEAVKHS
PDGVLRVFVKRTSGKS
>RA_YSBD_2000902_Heterosiphonia_pulchra.p1|419_517
PLASFKFKDINGDFRRIKVPMELQQGDFDQFVLDVRRRFLGQALSPSSLGGVKIKYMDEDGDDVLIAN
DDDLASCFEDVGDVKGKTIQLKVEEAEGAG
>GA_ZRMT_2007919_Mougeotia_sp..p1|772_848
GEKEVHRVKCFVEGSPFGVSVDLSRFHSTQELRESLLSVTEGTSVVYQDKDGDMILLGDESWAFFLQS
VRRMYIRR
>GA_ZRMT_2009027_Mougeotia_sp..p1|1_92
AADMLVVKIKYEEVLRRFSLPKSPSPTYGHLVKKVHDLFSLSNSVNLAITYVDDEGDVVTMSDDQDVH
DAVTLQGLNPLRLTVNKTSNHAT
>RA_ZULJ_2003006_Porphyra_yezoensis.p1|15_87
LINGTFEGEITRTFNTVGEDFKAFKANVEKAEKAEVIFQYKDDEGDVCLITSNDEFQEALRLYPAGLS
LILS

>kfl00094_0070_RAV
PESEEEQRPQVKCFVEESPYGVVVDLSQYKSTQQLKNALLDATHGRLVVYQDKDGDMILLGDEEWSFF
LKTVSRIFIRK
>MpRAV_Mapoly0072s0102.1
LQWSPEKFLNISLANFHSVEGLKRELLQFTHLDFAQGLDIVYKDKDGDIMLLSEHSWGLFKENVQEMW
IRKTEQPQQQRA

>AT1G04240_SHY2_IAA3|87_184
HEGQGIYVKVSMDGAPYLRKIDLSCYKGYSELLKALEVMFKFSVGEYFERDGYKGSDFVPTYEDKDGD
WMLIGDVPWEMFICTCKRLRIMKGSEAKG
>AT1G04250_IAA17_AXR3|105_216
GPEAAAFVKVSMDGAPYLRKIDLRMYKSYDELSNALSNMFSSFTMGKHGGEEGMIDFMNERKLMDLVN
SWDYVPSYEDKDGDWMLVGDVPWPMFVDTCKRLRLMKGSDAIG
>AT1G04550_BDL_IAA12|119_222
KVQGLGFVKVNMDGVGIGRKVDMRAHSSYENLAQTLEEMFFGMTGTTCREKVKPLRLLDGSSDFVLTY
EDKEGDWMLVGDVPWRMFINSVKRLRIMGTSEASG
>AT1G19220_IAA22_ARF19_ARF11|953_1056
TQRMRTYTKVQKRGSVGRSIDVTRYSGYDELRHDLARMFGIEGQLEDPLTSDWKLVYTDHENDILLVG
DDPWEEFVNCVQNIKILSSVEVQQMSLDGDLAAIP
>AT1G19850_IAA24_MP_ARF5|788_882
TPRVRTYTKVQKTGSVGRSIDVTSFKDYEELKSAIECMFGLEGLLTHPQSSGWKLVYVDYESDVLLVG
DDPWEEFVGCVRCIRILSPTEVQQMS
>AT1G30330_ARF6|791_885
NPQSNTFVKVYKSGSFGRSLDISKFSSYHELRSELARMFGLEGQLEDPVRSGWQLVFVDRENDVLLLG
DDPWPEFVSSVWCIKILSPQEVQQMG
>AT1G34170_ARF13|428_530
LLFGVDLTKVHMQGVAISRAVDLTAMHGYNQLIQKLEELFDLKDELRTRNQWEIVFTNNEGAEMLVGD
DPWPEFCNMAKRIFICSKEEIKKMKLKNKFFQPE
>AT1G62390_Phox2_CLMP1|285_387
KRIRWRPLKFVYDHDIRLGQMPVNCRFKELREIVSSRFPSSKAVLIKYKDNDGDLVTITSTAELKLAE
SAADCILTKEPDTDKSDSVGMLRLHVVDVSPEQE
>AT1G79570|188_281
PRPGDSKLRYVGGETHIISIRKDISWQELRQKILEIYYQTRVVKYQLPGEDLDALVSVSSEEDLQNML
EEYNEMENRGGSQKLRMFLFSISDM
>AT2G17150_NLP1|807_898
ARGAIKVKATFGEARIRFTLLPSWGFAELKQEIARRFNIDDISWFDLKYLDDDKEWVLLTCEADLVEC
IDIYRLTQTHTIKISLNEASQVK
>AT2G25290_Phox1|275_364
DATVTRTVKLVHGDDIRWAQLPLDSSVVLVRDVIKDRFPALKGFLIKYRDSEGDLVTITTTDELRLAA
STREKLGSFRLYIAEVSPNQE
>AT2G28350_ARF10|575_673
QGLETGHCKVFMESEDVGRTLDLSVIGSYQELYRKLAEMFHIEERSDLLTHVVYRDANGVIKRIGDEP
FSDFMKATKRLTIKMDIGGDNVRKTWITGI
>AT2G35050|187_281
IPRPRDQKLRYVGGETRIIRISKTISFQELMHKMKEIFPEARTIKYQLPGEDLDALVSVSSDEDLQNM
MEECIVFGNGGSEKPRMFLFSSSDIE
>AT2G36500|404_492
LVSSFAFKFEDRKGRVQRFNSTGESFEELMSVVMQRCEADSGLQIMYQDDEGDKVLISRDSDLVAAVT
FARSLGQKVLRLHLDFTETI
>AT2G43500_NLP8|829_921
SGSTTLIVKASYREDTVRFKFEPSVGCPQLYKEVGKRFKLQDGSFQLKYLDDEEEWVMLVTDSDLQEC
LEILHGMGKHSVKFLVRDLSAPLG
>AT2G46990_IAA20|79_175
NSVGSFYVKVNMEGVPIGRKIDLMSLNGYRDLIRTLDFMFNASILWAEEEDMCNEKSHVLTYADKEGD

WMMVGDVPWEMFLSTVRRLKISRANYHY
>AT3G24715|187_279
PRSTDEKLKYVGGETHIISIRKNLSWEELKKKTSAICQQLHSIKYQLPGDELDSLISVSSDEDLQNMI
EEYNGLERLEGSQRPRLFLIPIGE
>AT3G46920|18_117
IIPRPSDGMLRYVGGQTRIVSVKKNVRFDEFEQKMIQVYGHPVVVKYQLPDEDLDALVSVSSSEDIDN
MMEEFEKLVERSSDGSGKLRVFLFDASSSEV
>AT4G23980_ARF9|518_628
SSSTRSRTKVQMQGVPVGRAVDLNALKGYNELIDDIEKLFDIKGELRSRNQWEIVFTDDEGDMMLVGD
DPWPEFCNMVKRIFIWSKEEVKKMTPGNQLRMLLREVETTLT
>AT4G24020_NLP7|858_950
SEMRTVTIKASYKDDIIRFRISSGSGIMELKDEVAKRLKVDAGTFDIKYLDDDNEWVLIACDADLQEC
LEIPRSSRTKIVRLLVHDVTTNLG
>AT4G24690_NBR1|4_94
ANALVVKVSYGGVLRRFRVPVKANGQLDLEMAGLKEKIAALFNLSADAELSLTYSDEDGDVVALVDDN
DLFDVTNQRLKFLKINVNAGVS
>AT4G30080_ARF16|579_669
SGLETGHCKVFMESDDVGRTLDLSVLGSYEELSRKLSDMFGIKKSEMLSSVLYRDASGAIKYAGNEPF
SEFLKTARRLTILTEQGSESVV
>AT4G32070_Phox4|312_398
GATVTRTIKLVHGDDIRWAQLPLDSTVRLVRDVIRDRFPALRGFLIKYRDTEGDLVTITTTDELRLAA
STHDKLGSLRLYIAEVNP
>AT4G32280_IAA29|154_251
SSRSSMYVKVKMDGVAIARKVDIKLFNSYESLTNSLITMFTEYEDCDREDTNYTFTFQGKEGDWLLRG
DVTWKIFAESVHRISIIRDRPCAYTRCLF
>AT4G38340_NLP3|668_764
KAKDGMKVKAMFGDSMLRMSLLPHSRLTDLRREIAKRFGMDDVLRSNFSLKYLDDDQEWVLLTCDADL
EECIQVYKSSSLKETIRILVHHPLSRPS
>AT5G09620|49_152
IQPRPHDNQLTYVNGDTKIMSVDRGIRFPALVSKLSAVCSGGGDGGEISFKYQLPGEDLDALISVTND
EDLEHMMHEYDRLLRLSTKPARMRLFLFPSSPISG
>AT5G20360_Phox3|355_442
EDVNKDVKFVYSDDIRLAELPINCTLFKLREVVHERFPSLRAVHIKYRDQEGDLVTITTDEELRMSEV
SSRSQGTMRFYVVEVSPEQ
>AT5G20730_NPH4_MSG1_IAA21_IAA23_ARF7_IAA25_TIR5_BIP|1033_1136
TQRMRTYTKVQKRGSVGRSIDVNRYRGYDELRHDLARMFGIEGQLEDPQTSDWKLVYVDHENDILLVG
DDPWEEFVNCVQSIKILSSAEVQQMSLDGNFAGVP
>AT5G37020_ARF8|700_794
SNQTKNFVKVYKSGSVGRSLDISRFSSYHELREELGKMFAIEGLLEDPLRSGWQLVFVDKENDILLLG
DDPWESFVNNVWYIKILSPEDVHQMG
>AT5G49920|24_111
SDSVLKYVGGETRVVAVSPDISFSELMKKLTAITENDIVLKYQIIPEDLDALVSVKSDEDVKHMIEEY
NRHETPKLRTFLFPANPVV
>AT5G57420_IAA33|67_167
VSSVVPPVTVVLEGRSICQRISLDKHGSYQSLASALRQMFVDGADSTDDLDLSNAIPGHLIAYEDMEN
DLLLAGDLTWKDFVRVAKRIRILPVKGNTRQV
>AT5G57610|33_128
ILPRPQDGKLRYVGGETRIVSVNRDIRYEELMSKMRELYDGAAVLKYQQPDEDLDALVSVVNDDDVTN
MMEEYDKLGSGDGFTRLRIFLFSTPEQ
>AT5G60450_ARF4|660_752
SSSKRICTKVHKQGSQVGRAIDLSRLNGYDDLLMELERLFNMEGLLRDPEKGWRILYTDSENDMMVVG
DDPWHDFCNVVWKIHLYTKEEVEN
>AT5G62000_ARF2_ARF1_BP_ORE14_HSS|728_822
TNSSRSCTKVHKQGIALGRSVDLSKFQNYEELVAELDRLFEFNGELMAPKKDWLIVYTDEENDMMLVG

DDPWQEFCCMVRKIFIYTKEEVRKMN
>AT5G63130|24_117
ILPRSTDGKLRYVGGHTRVLSVDRSISFSELMKKLYEFCGYSVDLRCQLPNGDLETLISVKSEEELAE
IVEEYDRICGAKIRAVLSPPRSSHK
>Mapoly0011s0167.1|751_844
QGPVRSYTKIHKQGSFGRSIDVQSYDGYTDLLRKVENMFELNGELFDKKSGWQLVYTDHEDDVLLVGD
DPWMEFVSCVRTLRLLSPGEASSSG
>Mapoly0013s0010.1|658_770
GPNAPSPLTVVMEGRSVGRRLYLPDYDGYENFAEAMRSIFESYISSQEPRAHGESCTLANAIPGYVIA
YEDEEGDVLLAGDLPWREFVRVAKRIRIVSSSKISFKPPTGPSR
>Mapoly0013s0150.1|161_272
FPSSRVKFMCSFGGKILPRPSDQQLRYVGGQTRIIGINRDVNFSELRNKMRESFGQCYTFKYQLPDED
LDALVTVSSDEDLENMMEEYDKLEADGSSRLRVFLFPADQDAT
>Mapoly0019s0045.1|789_883
SSPQRSYTKVYKLGSVGRSLDVAQFTNYTDLRVHLARMFGLEGQLEDPQRSGWQLVFVDNEQDVLLVG
DDPWDEFVNCVRSIRILSPSEVMHMS
>Mapoly0034s0017.1|700_808
GQSNSWFVKVHMDGVPIGRKVDLRTNSSYEKLSQMLDEMFRTFVNGQNGSNRITLASDIKRNFLQGPD
YVLTYEDQDGDLMLVGDVPWTMFIDTVKRLRIMKGSEAIG
>Mapoly0039s0051.1|22_79
GEEGGLSFADLESSIRELFKISASSARLKVTYKDKDNDVVTMDSDQDLKDACVFQAL
>Mapoly0075s0050.1|487_579
PKPLRTYTKVYKLGSIGRALDVTRFSNYDELRCELARMFNLEGQLDQRSGWQLVFRDNENDILLVGDD
PWDEFVSSVRGIRILSPSEVNFYM
>Mapoly0083s0040.1|1042_1134
DESSSTIVKATYGSDTARFKLLPNSSYQDLREEVASRLKLPVQSLNLKYRDDDEEWVLLVCDADLDEC
IEVMRSTGGHAIKLMVRVETVNTS
>Mapoly0099s0038.1|274_364
AKHTPRPLKLVFGHDIRRAEVFGNCGFGDLREVVRKRFPGLKTVLIKYQDPDGDLVTITSREELRLAE
AAADEAGRSKKLDAGDTPVPDA
>Mapoly0100s0042.1|2_95
YVIKIKYEDTLRRLTYPGHRPYGDQYNDKLITFAKLEGNIRDLFKIPASSQIRVTYTDKDNDVVTMGD
DQDLEDACVLQGLNPLRLLVTLLDV
>Mapoly0179s0023.2|420_515
GMGNTFSFKLEDRKGRVHRFNCGTESLTELISAIVQRVGDDVDLNQLPHIMYEDDEGDRVLLGSDSDL
VAAVNFARISGWKGLRLYLDLPHFAPR

>Pp3c11_15160V3.1|406_507
SFGPGNTFSFKLKLSAYDGKPHHYRFNCGSESLTELMSTIAQRVGDDIDLTHLPRLMYIDDEGDKVLL
ATDSDVVAAVNVARISGWKALTLHLEESQEPRS
>Pp3c11_16970V3.1|1_95
ASYVFKIKHGDVVRRFTVQPSVPGGGPDMNFSQLEDTIRQAFKLPASSDLVINYNDKDGDVVTMAGDH
DLYDACVIQELNPLRLTVTAHESYPG
>Pp3c14_16990V3.10|723_826
PAPMRTFTKVHKLGSVGRSIDVQKFQNYSELRAELARLFNLEGLLEDPQTSGWQLVFVDNENDILLVG
DDLWEEFVSCVRSIKILSPNEILRMSREQLEILNP
>Pp3c15_9710V3.1|644_736
PKITRSYIKVYKLGSITRAVDVNRFKDYTELRCELARMFNLDGQLDPTVGWQLVFTDNEDDLLLVGDD
PWDEFVRNVRGIRILTPAEVYFYT
>Pp3c17_4370V3.1|1142_1234
VEPEAVTVKATFRADTVRFKLLVGSGYLDLRNEISRRLKVDEHGFDLKYLDDEEEWMLITCDADVKEC
IDVAQTLGRHTVKLMVRQSGSWTS
>Pp3c2_25890V3.1|646_740
APPVRSFTKVHKLGSVGRSLDINKFSNYVELRKELAHMFHLECLMEDSQQSSWKIVFVDNENDTLLLG

DEPWEEFVSCVRSIKILSPAEVAQMN
>Pp3c22_6370V3.1|1173_1262
EDSAAITVKVTYGLDTVRVKFAQNVSFVELKEEVGRRLKLAGQNFNLKYLDDDEEWMLLACDADLQES
IDLMRVSGRHAIKLMICSNML
>Pp3c24_6610V3.1|360_480
SSSSGNLVKIYMDGVPFGRKVDLKTNNSYEKLYYTLEDMFQHYINVHGCGGRSSSCGDSHSLASSRKL
NFLEGSEYVLIYEDHEGDSMLVGDVPWDWFIDAVKRLRIMKGSEQVNLAPKN
>Pp3c5_9420V3.1|655_749
VAPVRSGTKVYYSGKVGRTIDLKKCESYAALRRMLASLFGLEGQLDDVTKGWQLVYTDHENDVLLVGD
DPWEEFCNCVRSLKVLSPQDAAGQSV
>Pp3c9_25280V3.1|171_265
LPRPTDNQLRYVGGETRLITVSRDISYHELVLKMGELFSHFHTIKYKLPEEDLDALVSVSSNEDLANM
MEEYDRLQASEASPRLRIFLFSDMDH

>evm_27.model.AmTr_v1.0_scaffold00002.514|96_194
PEAPGFYVKVSMDGAPYLRKIDLKNYKGYQDLLKALENMFSCFTIGDFAKKEGSKTSEYVPTYEDKDG
DWMLVGDVPWEMFIFSCKRLRIMKGSDAIG
>evm_27.model.AmTr_v1.0_scaffold00007.258|75_171
IVPRPHDKALCYLGGDTRIVVVDRHTTLVELSSRLSRLLLNGRPFTLKYQLPNEDLDSLVSVTTDEDL
ENMVDEFDRCLRPARLRLFLFPTRPPDT
>evm_27.model.AmTr_v1.0_scaffold00007.382|837_930
QLCMRTYTKVQKRGSVGRLIDVTRYKGYDELRRGLACMFGMEGQLEDPHQSGWKLVYVDKENETLLVG
DDPWVLYEARTQLVLDLTLFPPPTY
>evm_27.model.AmTr_v1.0_scaffold00013.41|395_492
STGLASAFTFKLEDKKGRMHRFNCGTGSLSELLTSIVQRMGDDIDRNHLPQILYEDEERDKVLLATDS
DLTAAVDHARQAGWKGLKLYLDYSDSGSY
>evm_27.model.AmTr_v1.0_scaffold00017.72|401_497
SLGLGNTFSFKFEDRKGRVHRFNFGTENLGELSNAVLQRIGHSEEDRPPQLLYLDDEGDKVLLSSDSD
LVAATNHARIAGWKVLRLHLDYSDHQEK
>evm_27.model.AmTr_v1.0_scaffold00019.236|47_142
ILPRPLDGKLRYVGGETRIVSVQRDIIYDELMEKMRELFDGASSLKYQQPDEDLDALVSVVNNDDVTN
MMEEYEKLGAGDGFTRLRIFLFPHPDH
>evm_27.model.AmTr_v1.0_scaffold00019.282|51_158
SANMMPSVTVVLEGRAICQRISLQKHGSYQSLAKALRQMFLDISGDYSLEKSPENDCDLSNAIPGYFI
AYEDMEDDLLLAGDLNWKDFVRVAKRIRILPVKAKQKAY
>evm_27.model.AmTr_v1.0_scaffold00021.168|697_790
KTANRSCTKVHKQGSMVGRAINLSKFEGYDDLISELERLFNMEGLLNDPKKGWQVVYTDSDDDMMLIL
IYTHDEVEKMIPVVVASDDAQSCSE
>evm_27.model.AmTr_v1.0_scaffold00021.210|624_727
SISTRSCTKVLMQGIGVGRAVDLTKLNGYPDLIKELEERFDIKGELQSPNKKWEVVYMDDEGDMMLVG
DDPWAEFCNMVRKIAIYPCEEVKKMTNRTRLHCSE
>evm_27.model.AmTr_v1.0_scaffold00025.251|1043_1137
FPRMRTYTKVYKRGAVGRSIDITRYKGYEELRHDLALMFGIEGQLEDPLRTGWKLVYVDNEGDILLVG
DDPWDEFVSYVQSIRILSPAEVQQMS
>evm_27.model.AmTr_v1.0_scaffold00028.102|287_395
LSVRWRPLKLVYDHDIRLAQMPASCGYKVLREIVRKKFPCSKSVLMKYRDSEGDLVTITGTEELRLAE
LCADLLKKKDVVDGSIEGEVLEESLGVLRLHVVEVSPKQE
>evm_27.model.AmTr_v1.0_scaffold00029.187|703_798
KPHNPILVKIYKTGCVGRTLDISQFSSYEELRGKVADMFGLEGQLDDPLRSGWQLVFVDRENDALLLG
DGPWEAFVNNVWYIKILSPHDIQMMGT
>evm_27.model.AmTr_v1.0_scaffold00039.160|47_168
EKVITQFVKVNMDGLPIGRKVDLNAHGCYETLAQALEDMFQSPTATSIRQTGVHSALEKEHGGLSVES
KASKLLDGSSEFVLTYEDKEGDWMLVGDVPWGMFLSTVKRLRIMRTSEANGLA
>evm_27.model.AmTr_v1.0_scaffold00039.196|187_280

LPRPSDGKLRYVGGETRIVTVSKDISWHDLMQKTLSVHNQYHTIKYQLPGEDLDALVSVSSDEDLQNM
MEEYNGLENVDGSQRLRIFLISGNE
>evm_27.model.AmTr_v1.0_scaffold00049.221|29_123
PRHPDGKLRYVGGFTRVLAVDRSVHFSDLMEKLRKFCGEVAITLRCQLPTEDLDALVSVTSDEDLAHV
IEEYDRVNRERGSSLKVRAFLSPPKP
>evm_27.model.AmTr_v1.0_scaffold00049.238|1_94
ADDLVIKVKYGSTLRRFSIWAVGNGTSGFNMDMLKDKIRKLFNLKPIEDFAITYIDEDDDVVTLADDD
DLSDAIRQHLDPLRLTVSLTETPWE
>evm_27.model.AmTr_v1.0_scaffold00056.118|225_337
APKKGMFVKINMDGIPIGRKVDLNAYRSYEKLSGAVDELFRGLLAAQSDSSASSTQSGIGEKLRGLLD
GSGEYTLVYEDNEGDRMLVGDVPWEMFVSTVKRLRVLKSSELSG
>evm_27.model.AmTr_v1.0_scaffold00066.150|923_1013
GSSRVTVKATYKDDVVRFKFLPCLGYFQLFEEVGRRFKLSIGTFQLKYLDDEEEWVLLMNDSDLNECL
EILESYGAHSVKLMVRDLPCVM
>evm_27.model.AmTr_v1.0_scaffold00066.249|241_330
KVEPTKAVKLVLGEDIRWGQLPVNCNIRQLREIVRNRFPCSKAVLIKYRDVEGDLVTITTTEELRWAE
ESADSQGSLRLYIFDVGPDQE
>evm_27.model.AmTr_v1.0_scaffold00080.66|809_903
HGLEWVTIKTSYQDDMVIRFRLSSGSGIFDLHREVSKRFKMEVGAFGIKYFDNDKGNWVLLSCDADLH
KCIDASTSSGARVIRLSMTFSNQATR
>evm_27.model.AmTr_v1.0_scaffold00081.26|35_134
FIHSSQAGKTRYCGGENRLLCIDKNTSFSLFIFKLSKLCPFHSFSLKYMLPDSCSCTPLVSVSGEDDY
CNMIEMNLEFFAEGKTPRFRFFLFPQPLTRY
>evm_27.model.AmTr_v1.0_scaffold00109.93|40_140
IQPRPYDNLLSYYGGETKIVSFDRNVSFSAMKAKLLSLCNLPDIIYKYQLPGEDLDALISITDEEDLE
HLMSEYDRLQRSYGKKPVRLRLFLFPNYQSPI
>evm_27.model.AmTr_v1.0_scaffold00122.5|68_166
EQQKKNYVKVGMEGTPYLRKMDLKMFNEYSELSKALAAMFGCFSIGISNAKEERKSSEYFPIYEDDDG
DWMLVGDVPWEMFVESCKRLRIVKNDVKIR
>evm_27.model.AmTr_v1.0_scaffold00148.24|636_726
VEEGNGHCKVFMESEELGRSLDLSAFGSYEELYRKLATWFGIERSETLNHMLYRDESGTVKQTGDEPY
SEFMRTARRLTILSDSSSDNMG
>evm_27.model.AmTr_v1.0_scaffold00200.14|104_181
GRGSSFYVKVNMEGVGIGRKIDLNLHHSYSSLAHALEDMFGKHEKNKEDGTVCCPQCILTYEDREGDW
MLVGDVPWE
>Acanthoeca_like_sp_10tr_CAMPEP_0199716520|61_124
RRFQLPGHSAAELTRKLVSLFPQLEQGVFSVTWRDEDNDWVTVASDDDLAIAVESTAGQTLRL
>Acanthoeca_like_sp_10tr_CAMPEP_0199725100|2_83
ASIKVFYTRPGVPDDVRRFKLDEPLQFDALQAKLAAIYGVAYTNLTWKDEDGDAVTIGNAEDLDEAAR
SCSGTLKVYAAEG
>Alexandrium_monilatum_CCMP3105_CAMPEP_0200517146|8_90
LTLKVNLDGDVRRFKDSSPELQAIHEAIATLFSLSSEEAQKLALQYEDDEGDLCALTEVTLPDALELA
GRSKDVLHIYCSRG
>Alexandrium_monilatum_CCMP3105_CAMPEP_0200625392|1_86
KADLAGDIRRLKPWPSSGVEPSLAAARQAVGHLHSLAEEECGSLVLKYQDDEGDLCTLTDASLPDALH
LATENSAGVLRLSVLRE
>Alexandrium_temarense_CCMP1771_CAMPEP_0186390228|74_158
TLKFVHGEEIRRFRVEDHAAFSFGDLVSTIEAVFADASRTLPAKYSITYQDDEDDTITLSCDAELAEA
FRVALAESRKTLRLTI
>Amphidinium_carterae_CCMP1314_CAMPEP_0186415190|3_89
AILKVKHGSEVRRSQLEVDCLTYDRLAAAVTAMYPDCNNYLLKYRDPEGDLCTLTSTTYHDFKSLVEH
DIKGNPGQKIFRLELTPA
>Amphiprora_sp_CAMPEP_0186512182|19_123
TTIKLALEGPNAEKEGPFVKIPVNGPIGNQTLETLRQKSLDLWRATNGSFDDENLDEAKVLLTYRDSE

GDLCFVFSEGDLRCAIPMFPTSLKMFAKIELSQKKK
>Amphora_coffeaeformis_CCMP127_CAMPEP_0186530314|177_276
VRFKVKSTTSGTVLRFQCVPTYINLLSNISERLRRNSKTSSLWSTATNLITSSDSGLTIEFQDAESDW
CTLTNDEDLQDAVEASLGRKDDMVRLNIEEH
>Aplanochytrium_stocchinoi_GSBS06_CAMPEP_0200739792|553_633
KVKILSNGKMHRISLSPEMNWQVLKSEILSMAKMAASDVSFAYVDEEGDPVTISSDSALKEAQNMCQM
HNLGKLTILVTE
>Asterionellopsis_glacialis_CCMP134_CAMPEP_0199906488|521_604
ILFKVVDPNGKTHRIRSDLKLDKLLNQLLKKIKGVEDSSSIILNFSDDEGDEILITNDDDLIESVNVA
RSTGNKVVKLTATLQ
>Aurantiochytrium_limacinum_ATCCMYA1381_CAMPEP_0186637274|630_700
KVYRLSDKSVSSLQELLRQLERATGQDADEMQLVYLDEDKDRVQLANNLALDEARALAKQQGWKKIDV
FI
>Ceratium_fusus_PA161109_CAMPEP_0199433280|3_73
KVELRGEIRYFARPSSLNALLEDAQKRFGEALHPWLEHPDCDGDIVRISNTEELLIAMGISEQRGIPL
RL
>Chaetoceros_neogracile_CCMP1317_CAMPEP_0201016606|1209_1276
TYLVRSDNNYADFLKALSNKVKKELDPKKIQLKYIDDEGDTILIGSNDCLSEAIVNSRKKGNQAVKL
>Chattonella_subsalsa_CCMP2191_CAMPEP_0187147442|27_103
RLKCSYQENLRAVLVHKGFKFSDLKKRLEQDFGFEVMLQYRDEDGDLVLLASQDDLDDLFGYMEEMHP
ATSKFHVG
>Chattonella_subsalsa_CCMP2191_CAMPEP_0187161922|118_179
RVKLTLDDEVRAVRLPVQITLPALNEQIKRDYGAEMEISYMDAEGDKVSIRTQEDLLLALK
>Chattonella_subsalsa_CCMP2191_CAMPEP_0187170092|3_83
NSLKVCFGGEIRRIRVIPSDVSHLRALLEERFNLPSNYSITYKDDENDTITIQTDEELSEAIQITSED
DRKTLRVEVIST
>Chattonella_subsalsa_CCMP2191_CAMPEP_0187173358|8_87
MIVKLHYESQLSLAEIDENTYECLVHKICEIFSLKADQLLLHFKDNEKAWVSLKSTEDMLLGMKICEA
RKPLHILVKQK
>Chrysochromulina_polylepis_CCMP1757_CAMPEP_0193731940|7_86
KLSFGDEMRRFKINTEDITYQTIHTRAEALFGFAPDTKFKLQYKDDEGDVITMATDEEMFEAVALAVS
IEPAILRVSVK
>Dinobryon_sp_UTEXLB2267_CAMPEP_0187276494|279_359
NCKRFCFDEAMIFDDHRNISFDKMRQQIIEYFPSLRERTFKLMYEDNEGDVITVSSDCELVEAFRIMN
SMNVISFHVILL
>Dinobryon_sp_UTEXLB2267_CAMPEP_0187279124|1_84
TAVKVYFQEETRRLQLDDSDLNFKSLFRAIENIFPSLNGNFCIKWKDEEGDLISFTSDVEFAEAVRIM
SDVSDSLLRFHIQEA
>Ditylum_brightwellii_GSO103_CAMPEP_0187300826|16_108
VVLKLSLGEGDKAQVRRITLSRLWENGKVSYNRLISIVREHSLEEAEVTITYDDEDGDVITISTDDEL
ADAFGQFVESQPPVLRVKGHVKEV
>Ditylum_brightwellii_GSO103_CAMPEP_0187314946|6_102
LILKITLGENETHRTTCNSIWDESESKVSYDLLVDLVTKYAFPTHNATGLPKDYEVVVTYVDEDGDKV
NISSDVELNDAVEQFVSCKPPVLRIKAE
>Ditylum_brightwellii_GSO104_CAMPEP_0193941848|622_700
FKVVDGEGHTYRIRCECKLDKLVSVVISKVGGDLEANSVKLKFIDDEGDTILIGSDDCLAEAIGLARN
AGSEVVKLSL
>Dunaliella_tertiolecta_CCMP1320_CAMPEP_0187378638|270_355
GANNAGANGAAPQGSHSATVHLGGLTSFDDLRAAVQAAFAPGLLPPQPQAKLVYLDADGDWMMITRDD
HWASFCATARKLLVTDR
>Durinskia_baltica_CSIRO_CS_38_CAMPEP_0199947424|399_475
DFKVTDSAGLLHKIRCTVESLSALRSAVAQKFLAPAEHVLLKYIDEEGDEIVISADMVLREAVESVRH
MGGVSLKL
>Emiliania_huxleyi_374_CAMPEP_0187584202|89_170

VIVKISGNNQTRRWTSRADHLTFASIQKRVADNFQTPKFYLTYKDDEDDDINVSTDEELQEAVSLALK
TEPAVLRLKLVEG
>Euplotes_focardii_TN1_CAMPEP_0187816052|5_80
TIKTFFGADIRRFQLVEGAPLFDTFSDWVATTYNLPNGEATFKYVDDEGDMCLITSEEELQEAIRLSA
EVLKITV
>Euplotes_focardii_TN1_CAMPEP_0187823868|20_83
YDDLSELRTIIEGSFKSLEPNSYTLKYLDNENDWLYIFDNTDLQALKEYHQEKNGKSIKLVVE
>Euplotes_focardii_TN1_CAMPEP_0187833828|30_89
QDTSTFEEVKSEVCKGLDIKDMEIFYTDDEGDTIRVCNDQEIAEAFAVARADNSITFYV
>Eutreptiella_gymnastica_like_CCMP1594_CAMPEP_0200384762|30_100
EQIRRFSVQKESKLEAIKDRLNEMSGMDNGYQIHYVDPDGDQVLISSQEEWEECLNVLGDPLTLRLHV
SE
>Eutreptiella_gymnastica_like_CCMP1594_CAMPEP_0200419268|11_93
IHVKAYLHRTRGDQIRRFGVPQNVTWAAMIDQLIQLYGAGGYEVRYVDPDGDQVLVSSQEEWEECLSL
VKVSAVVYLHLNKQ
>Extubocellulus_spinifer_CCMP396_CAMPEP_0200448384|166_264
AVMKLQLGERGGTRRINFSRLWDVSRNSVSYSKLFSLAVEYSFPEDAPIDLSNYDIVITYLDEDGDTI
TVSSDDELSEAFMQFVGKSPPILRATAKVK
>Favella_taraikaensis_FeNarragansettBay_CAMPEP_0199846172|1_81
QAKKAVEGLLQGETKRLKLSGDYQELIERTRSAFKNLPPSYKFFYLDEDNEIISISSQSDFTEALEIE
DLSALRLTVAVK
>Fragilariopsis_kerguelensis_L26_C5_CAMPEP_0188079814|2215_2301
VVFKIVDPDGHTHRFHSETKIVCLLEAFVKKLAGRLRAKNVSLKFFDEEGDAILISTDNDLYEAVVLA
RNASQGSKVVVKLICEVK
>Gephyrocapsa_oceanica_RCC1303_CAMPEP_0188224140|60_146
KEDGDHHREDLRQQPDPALDDTGDHLTFLSIQKRVADNFQTPKFYLTYKDDEDDDINVSTDEELQEAV
SLALKTEPAVLRLKLVEG
>Gephyrocapsa_oceanica_RCC1303_CAMPEP_0188226736|457_543
SAFLFKLLDAGGAVHRIRHASPSLEELRAAAFGRLSAGPPPAGSLLLYEDEDDDRVVLSSDAELADAV
QSARAAGRDRLVLHVVKR
>Gloeochaete_witrockiana_SAG46_84_CAMPEP_0193988646|4_82
VAFKAKFGDELRRFTHDVSSGNQLTFDWLLQTVSELFRVPGRKLRVQYKDEDGDLIEISSDNELAEAV
RIAQLNKQGL
>Gloeochaete_witrockiana_SAG46_84_CAMPEP_0193988778|5_83
KFIFNQDIRRVSLHNEELSLQKLRNVAVQLFGSALPESFVFQYTDSDGDAITIASDIELEEALKFSSS
PSGLLRLTIV
>Gloeochaete_witrockiana_SAG46_84_CAMPEP_0194024110|1143_1228
FTFKAAFREEIARFKFLSSQTLSDLVAEVRSAFGHIITSEQHIALKYKDEEGDFISLVRDSDLAECLL
VGRSANNRCSIAVAISS
>Goniomonas_pacifica_CCMP1869_CAMPEP_0188456340|51_132
IAFKARLREELRRFSLNVPFDLPALVAQVARVFQLDVGRVNLTYLDEDNDFITLSNNEDLRELLRLHG
ESPQAVRLFVNRK
>Goniomonas_pacifica_CCMP1869_CAMPEP_0188459158|4_85
IAFKARLREELRRFSLNVPFDLPALVAQVARVFQLDVGRVNLTYLDEDNDFITLSNNEDLRELLRLHG
ESPQAVRLFVNRK
>Goniomonas_pacifica_CCMP1869_CAMPEP_0188481442|18_91
KFAFLDDIRRVSLPDEARLSDVQRVVCERFDVENVKLMYRDDEGDTITVGSDEELREAFRLASSLSQC
LRFDV
>Goniomonas_pacifica_CCMP1869_CAMPEP_0188482608|86_166
VKCSFKQAEDEEVIRTLGLPRQISFQTLRTSLEREFGPVVAKYVDEDGDMVLLASSADLVECLGALPE
SGRPMLRLHLVP
>Goniomonas_pacifica_CCMP1869_CAMPEP_0188483422|1_85
QVVKVVYSEGEREDIRRIVFAPDVSFSDFVDVITKRFGFEGVRLTYRDDEGDLINLESEDELREAFRI
AQTFSSQTLRVMVAQG

>Goniomonas_pacifica_CCMP1869_CAMPEP_0188534744|1_78
LRSGARGGESXLRAAASYSEICAALFGLFGEPSSQGMRLEWRDGEGDWIRIDSEAEWHEAVRCTPPNH
TLRLRTQEK
>Goniomonas_pacifica_CCMP1869_CAMPEP_0188538062|25_104
FSIKATHAATRDVRRATIQEENINSLQEKLKALFNSPQPFDLLYKDDEGDLVRFSTNEELNEGLRLCP
SRVLHLVVCEE
>Heterosigma_akashiwo_CCMP2393_CAMPEP_0188545816|2_80
ILVKSNYMNDLRVFELLDSILFESLQQEVENQYGLRRNEFSVKFQDSEGEDVTIASDTDLKLALRLTK
KIFRIKICPV
>Heterosigma_akashiwo_CCMP2393_CAMPEP_0188556866|5_80
LKVCCNGEVRRLRDVPTEFEELRKLLQDRFALPSNYSIVYKDEDGDLITITTSDELLEAIRVCEENGM
KSLRVEV
>Heterosigma_akashiwo_CCMP2393_CAMPEP_0188564074|397_475
FKVTDAAGHMHRITAAADNLAGLTRAVAAKLAAEADAVVLKYTDDEGDEVAITCQQSLREAVNHAREA
GNKALRLKVV
>Heterosigma_akashiwo_CCMP2393_CAMPEP_0188579542|155_223
RVKFALGNGEVRAARLPMEMTLKKLRKQIRKDYGAQKMLISYKDSEGDQVAIRTQKDLRLALRLQGKL
>Heterosigma_akashiwo_CCMP3107_CAMPEP_0188607612|3_83
DTLKVCCNGEVRRLRDVPTEFEELRKLLQDRFALPSNYSIVYKDEDGDLITITTSDELLEAIRVCEEN
GMKSLRVEVLTP
>Isochrysis_galbana_CCMP1323_CAMPEP_0193632134|101_182
IIVKIAGNDQIRRWTAKADQVTFAALEQRVSETFKTPKFTLTYKDDEEDDITLGCDEELADAIALSLK
SEPSVLRLKLTDL
>Isochrysis_galbana_CCMP1323_CAMPEP_0193679308|390_474
FLFKIADDGTVHRVQNAGTSIASLRAAVLARLNVPPAMAASTVLLYEDDEADRVILTTDTELRDAVQA
ARGVGRDRIVLHVKRT
>Karenia_brevis_CCMP2229_CAMPEP_0188978606|4_83
LKLKCEGEVRRVHLKEDQELTYEVICENIRTAFPEIGPHITKYPDDEGDLCTLCSTTFADFLSLADGA
TGQKVMKLEIV
>Karlodinium_micrum_CCMP2283_CAMPEP_0200759400|7_93
LTLKIEYQGEVRRFRDWPANDAPASVEEMRASVCQLFGLHLTENDALTLKYRDDEGDLCTIAEPSLSD
ALMFASESGILRLTASLV
>Lingulodinium_polyedra_CCMP1738_CAMPEP_0189995912|1_79
KLTFNGDIRRTRLSGLTWEELQRAVAISFPELSKEDLGLKYRDDEGDLCLLTPASLSDLVALSGQATT
VKLELVLRRP
>Lotharella_globosa_CCCM811_CAMPEP_0190145524|22_116
VTVSHGEEHGLLDIEENECVNSLHQKILVRFPELKSEDFVVKYKDDEGDWITVGTTSDLREALEFLRE
TNEDVPNSAARYRESRVSQGTLQLII
>Lotharella_globosa_CCCM811_CAMPEP_0190147092|43_121
IKLEFEGVIKRREAPDDFSELLQILPKVFPGSSLEISGTAPEVAYVDSEGDVLAVQSDDDLLDACEWS
STSGKALKLK
>Lotharella_globosa_CCCM811_CAMPEP_0190151114|5_90
RNVKVVYGGDIRRAFLLVDDETDLYSELEALIVKLFGSSAGKDTKIRYKDEDGDMILVCSTDELADAF
AQAKAQGKTLKLFVQKA
>Lotharella_globosa_CCCM811_CAMPEP_0190151218|27_106
TICKIVFGNDIRRMPLPKDKPFNHLLLKLEAIHPGKFDGGIAVRYKDDEGDLVTVTSNLELMEAFEMS
DSTLRLLISAK
>Lotharella_globosa_CCCM811_CAMPEP_0190166318|3_89
THVKLIFEDGPKRDVRRFRLAQDATGQYKLICGMVKSLFGIRESFLQYIDEEGEKVVVGSEYEWEEAV
AFARRQKSKTVKLFLRKG
>Micromonas_sp_CCMP2099_CAMPEP_0190189722|551_625
KLFLEDDIRFLEVDPDASYEELIIAVGKVFIETYVIKFEDQDNHHITLKTSDDLKIAIRQSEKAESPY
LRLVLE
>Micromonas_sp_CCMP2099_CAMPEP_0190192152|159_244

YKGGDTRLVSISRGVSFTNFVELLDLSLASAKLHRIIYRDPNDANALIRVTNDADIDELFQEWDRYVE
GLMTGGVPAAKLKVYVV
>Micromonas_sp_NEPCC29_CAMPEP_0190201886|205_267
KLMLKDDMIRMRLLNEVSHRALIKRVAEVMDIPEHQVRLKYKDDEDDWCVLATDADLKDAYG
>Micromonas_sp_NEPCC29_CAMPEP_0190216150|1053_1134
LTVKVSVNTGDNVRFKLLPGMSYVDVQSRLKESIGGQVGDLRLRYQDEVDEWCALNGDSDLAECIHVC
QSRGIVRMQAADD
>Micromonas_sp_RCC472_CAMPEP_0190221402|559_634
VKLFLEEDIRFLELDPDVSYEELIIAVGKVFIETYVIKFEDEDKHHITLKTSEDLKIAVRQYEKNESQ
YLKLILE
>Ochromonas_sp_CCMP1393_CAMPEP_0190269606|12_71
RFKCRYEGEIRTVIVTPETDMSVLKRRLSSDYGFDVALKYEDKEGDFIVLSSQNDLNDL
>Oxyrrhis_marina_CAMPEP_0190378796|30_108
TVKITGNNQIRRWTTTADQMNFVALQKRCAEFFQTSKFSLTYNDDEGDDVTLGCDEELREAVAIALQS
EPPVLRLKLT
>Paraphysomonas_imperforata_PA2_CAMPEP_0190464200|33_115
TALKLCFNGEIRRLSVSTSALTFDELVKTTLRLFPHLSSIQFSWVDDEDDKVIVGSDDELFEALRVMA
SEKKGYLRFEVCVG
>Paraphysomonas_imperforata_PA2_CAMPEP_0190480052|8_67
RLKCRYDNEIRIVIISADYSFTQLYNRLTLDYGFDISLKYEDQDGDLITLTSQNDFEDL
>Pavlova_sp_CCMP459_CAMPEP_0190507586|69_152
TVLKIVYGKDIRRCTLSEGSATWSALQRRVIDLLGVDLSGLRVTYKDDEGDVVTITTDGELEEALRLA
AQSTPAILRLTVLKK
>Pelagomonas_calceolata_CCMP1756_CAMPEP_0199681208|110_180
LKFHHRRAIRVVNVRRGESVSFRALKQRLTSDYGFELALAYQDADGDLITLASQNDLNELLDGFSSTN
SQ
>Peridinium_aciculiferum_PAER_2_CAMPEP_0190656522|5_80
TLKVDLSGDVRRCRLEDGSEHTLENFLQAICVLFGMPAGSLFLQYRDDEGDLCSLVEATLGDALAHAR
APGVLRL
>Pleurochrysis_carterae_CCMP645_CAMPEP_0190763832|44_127
IVVKVTNGKEFRRLTQIDTYEVLAHRAMKGLSSSPNFALHYTDDEGDLVLMSSEAEWQEAVRQAKAQA
ETTKTNPPILRVTLT
>Prymnesium_parvum_Texoma1_CAMPEP_0191233966|3_87
ITCKLSYGDEIRRITFDKSSEVTYERMCNCVNDFFGLAPQSFKLQYTDDEGDTITMSTESEMKDAVKL
VKLLDPIILRLYVKLR
>Pseudo_nitzschia_fradulenta_WWA7_CAMPEP_0199776800|806_895
GYATFKVQDPDGHTHRIRSSTRIAALRKSLAAKLKGRAYCEKNLSLKYFDEEGDAILISTDEDLVEAV
NLARTTSRESKFIVKLIAEEE
>Pseudo_nitzschia_fradulenta_WWA7_CAMPEP_0199804386|3_80
VLKVAFGSDIRRKRFSSASEVTLQAVDAFVRETMGAEGGSGNYVAKYCDDEGDLCTLADHSLEDALQL
AAEKQLLRL
>Pseudopedinella_elastica_CCMP716_CAMPEP_0191413862|20_100
IKFDFIASGEKRRVPRSTLGSYAELRALLAEYGADEPCVVSYLDEDGDAVLVTSDLELEEAFAVAKSM
NAKVFKFTVHPS
>Pseudopedinella_elastica_CCMP716_CAMPEP_0191414076|2_82
VKIIFKNETRRLQLSESSLNFHVLKKTITDVIPSLRGQDFLIKWLDDEDDLIALSSDIELSEAVRVME
NLKSNKLLRFQV
>Pseudopedinella_elastica_CCMP716_CAMPEP_0191457318|799_879
APFTFKIEDPNGNLHKFISASENFDQLVRVVSERLYTDIKTLKLKYKDEDGDEVLLTTDASLAEAVDQ
ARAKGDTSVRVR
>Pteridomonas_danica_PT_CAMPEP_0193806382|3_92
RSLKLSYHETIRRFSMKSMNIEEGKLSFEDLKDVVCKVFTELQNKEFVLTYKDSDNDIITLCNTDELN
EAFSVMNSLCLDVLRLMVDDC
>Pyramimonas_parkeae_CCMP726_CAMPEP_0191469370|607_686

PESPCVYVGGETRMVKIRSGSSYLELISQLGIFCKGPPLVRYALPGEEVMVSLRTDNDVRLMFEEWHL
WRRSDQSRNPE
>Pyramimonas_parkeae_CCMP726_CAMPEP_0191480616|4_84
ITVKSDLQGDKRRFQLDVRGGFDALRLAVETAYGECNLLIKYEDDEKDLVTLVSDDDLQEAIRVLNSM
NLASPLRLSVFN
>Rhodella_maculata_CCMP736_CAMPEP_0191525956|3_88
LHFKATFNSDIRRFSIPSTPDAISALLSTVHALFSPEAPAAALAVRYVDDEGDLVSLRTQADLDEALR
LVPPDPSKPIKLQVSLP
>Rhodomonas_sp_CCMP768_CAMPEP_0191552814|19_98
VTFEETTRRFVVNEPCTYAELWEIVAKRFEVPTNVTGSLRYTDPDGDLISLSTDDELKEALHLVSDSD
PPLLRMKLLLD
>Rhodomonas_sp_CCMP768_CAMPEP_0191560200|504_600
TPVKVTLNSSGVDSVIRRLSLGLRLRSDGTPVGGFQTVMASVSAAFKADLQPAAPTTLTYRDEDGDAV
VISSDHELGIALQHRAGGPLLRMTLSSR
>Rhodomonas_sp_CCMP768_CAMPEP_0191568790|5_85
YSLKVAFKDQYRRFLLNDLDYGNLLAMAADRFGFEDSDALYFHHRDPDGDLVTISSSEELAEAMRLLA
PEATLRLILAER
>Schizochytrium_aggregatum_ATCC28209_CAMPEP_0191599156|605_690
LVLKVKDQGDAKVYRLSDRSVRSLDDLLAQLERATGRPAEALDLVYLDEDGDEVQLANDLALREARSL
AAQQGWKKVDVLIRPPR
>Symbiodinium_sp_C15_CAMPEP_0192425966|35_114
FVLKTSYQGDIRRLTFDSVEQVSFESIRTFVAKSYDVQDFLAKYQDEEGDWCTLTAATLTDAMDLVKD
KHLLRLEVQAA
>Symbiodinium_sp_C1_CAMPEP_0199641460|140_224
VGVKTSLGHDKRRFQLAPGSRIEDLHRIVAAVHALDTASATLRTCWTDEDGDRITIATTDDLVEAVRW
AAATDTRLLRIDVAAE
>Symbiodinium_sp_C1_CAMPEP_0199642500|3_87
VTLKLIFGDDRRRARVPSTDGFTFAALEKLVLASFPEVNEPITIQYVDDEDEVITVATDVELAEAFRV
ASEEGRKVLRITVSGA
>Symbiodinium_sp_CCMP2430_CAMPEP_0192498336|2_94
AVLKATYRAEVRRCLLDKDDLCFQGLSRRVSELFPELPQYTAKYMDEEGDVCTLCEASFADFLQVSTQ
TKSASGNMPGSDTKVILKLELQAV
>Symbiodinium_sp_Mp_CAMPEP_0192602324|7_80
LKVCYKEEVRRLKDWPQGEPSFEDLQRSALALFDLSEQHCVDYQDGEGHLVSLTEEKLPDVLQLAAQF
GLLRL
>Tetraselmis_striata_LANL1001_CAMPEP_0200905450|308_397
FTAKCTLEGDTRVLYLSHNTTYSELVAKVKQMFPNAGAFTIKYTDKENDTITVTSRNDIHTAMAESLK
QMEKRPGSTVIPPLRFEIVKD
>Tetraselmis_striata_LANL1001_CAMPEP_0200907914|480_547
KFTLRKDTRRWTPAERPRLAELLDKVAAIYDLPEGTKTKLKYTDVDGDMVTLASETDVEELFRQALP
>Tetraselmis_striata_LANL1001_CAMPEP_0200926804|560_641
DGIECRLEGTDSVEFVRLSEHCNASSLTSYLRQIFSQKLAEEDCNFGVVYQDASGDWMLLQPGEPWTL
VVNVSRRLLVTTK
>Thalassiosira_antarctica_CCMP982_CAMPEP_0200132098|1094_1185
PVQTAPFEDLVAYKVVDGAGQTYVIRAGKTLDSILKVLEGKVSNLDPSTTVFKYFDDEGDEILIKSDE
CVEEAVRSSVQSGNKNVKLSMKA
>Thalassiosira_miniscula_CCMP1093_CAMPEP_0201084540|417_501
DTVAYKVVDDAGHTYVIRAGKTLDSIVKVLEGKVSDLDPSSVVFKYFDEEGDEILIRSDDCVEEAIRS
SAQAGNKNVKLSMKVS
>Thalassiosira_oceanica_CCMP1005_CAMPEP_0192893646|69_157
LKLKLEKGAAVQVRRIRVSELFVQGKNLSYDKLMGVAAGFLDDIGKTTLAAKSQVTYIDSDGDKINIS
SDEELNDSFEGILKKLPIIT
>Vaucheria_litorea_CCMP2940_CAMPEP_0199162090|642_731
DDVMFVYKVTGPEGDTHRVRSSAENFSMLQMAVAEVLNFSLDSVNLKLHYLDDDGDRCMLTGDETLHE

AIDMARAAGWTVLKLTALFDS

>NP_000622.2
NWLRCYYYEDTISTIKDIAVEEDLSSTPLLKDLLELTRREFQREDIALNYRDAEGDLVRLLSDEDVAL
MVRQARGLPSQKRLFPWKLHITQK
>NP_001040687.1
QVKSKFDSEWRRFSIPMHSASGVSYDGFRSLVEKLHHLESVQFTLCYNSTGGDLLPITNDDNLRKSFE
SARPLLRLLIQR
>NP_001040901.1
QFDAEFRRFALPRTSVRGFQEFSRLLCVVHQIPGLDVLLGYTDAHGDLLPLTNDDSLHRALASGPPPL
RLLVQKR
>NP_001093521.2
EVKSKFEGEYRRFALKKNTGGFQEFYQLLQTIHRIPGVDVLLGYADIHGDLLPINNDYNFHKALSSAN
PLLRIIVQK
>NP_001107789.1
IRIKTPVGDMDSPVDSPSHLNFNDLLAAIRDAMPEATVTAFEYEDEVGDRITVRSDDELKAMLSYYCN
TVMEQRFNGLLPEPLQIFPRAG
>NP_001124131.1
VKSKYGAEFRRFSVDRYEPGRYKDFYRLIVRLHQLWHTDVFIGYADVHGELLPINNDDNFCKAVSSTQ
SLLRIFIQ
>NP_001182408.1
LIIKAQLGEDIRRIPIHNEDITYDELVLMMQRVFRGKLLSNDEVTIKYKDEDGDLITIFDSSDLSFAI
QCSRILKLTLFVN
>NP_001193733.1
MEGHFPQSDVIGQVLPEATTTAFEYEDEDGDRITVRSDEEMKAMLSYYYSTVMEQQVNGQLIEPLQIF
PRAC
>NP_001239372.1
KAQLGEDIRRIPIHNEDITYDELVLMMQRVFRGKLLSNDEVTIKYKDEDGDLITIFDSSDLSFAIQCS
RILKLTLF
>NP_001252262.1
LKARHVDFVHKTSIHHTNDLTLIDLVLNVQWRLALPSDANFVLKYKNKQGDLVTLVVDSDLLMVLHTS
GA
>NP_001299842.1
MTVKAYLIGKEDCNKEIRRFAVDQDVSTSFEYLQRKVLDVFVGLRTAPFQMYYKDEDGDMIAFSSDDE
LMMGLALVKDDTFRLFIKQR
>NP_002731.4
VRVKAYYRGDIMITHFEPSISFEGLCNEVRDMCSFDNEQLFTMKWIDEEGDPCTVSSQLELEEAFRLY
ELNKDSELLIHVFPC
>NP_002735.3
RLKAHYGGDIFITSVDAATTFEELCEEVRDMCRLHQQHPLTLKWVDSEGDPCTVSSQMELEEAFRLAR
QCRDEGLIIHVFP
>NP_003891.1
LTVKAYLLGKEDAAREIRRFSFCCSPEPEAEAEAAAGPGPCERLLSRVAALFPALRPGGFQAHYRDED
GDLVAFSSDEELTMAMSYVKDDIFRIYIKEK
>NP_006061.2
LIIKAQLGEDIRRIPIHNEDITYDELVLMMQRVFRGKLLSNDEVTIKYKDEDGDLITIFDSSDLSFAI
QCSRILKLTLFVN
>NP_006600.3
VRVKFEHRGEKRILQFPRPVKLEDLRSKAKIAFGQSMDLHYTNNELVIPLTTQDDLDKAVELLDRSIH
MKSLKILLVIN
>NP_009359.1
ILFRISYNNNSNNTSSSEIFTLLVEKVWNFDDLIMAINSKISNTHNNNISPITKIKYQDEDGDFVVLG
SDEDWNVAKEMLAENNEKFLNIRLY

>NP_032703.2
NWLRCYFYEDTGKTIKDIAVEEDLSSTPLFKDLLALMRREFQREDIALSYQDAEGDLVRLLSDEDVGL
MVKQARGLPSQKRLFPWKLHVTQ
>NP_032883.2
RVKAYYRGDIMITHFEPSISFEGLCSEVRDMCSFDNEQPFTMKWIDEEGDPCTVSSQLELEEAFRLYE
LNKDSELLIHVFP
>NP_032886.2
RLKAHYGGDILITSVDAMTTFKDLCEEVRDMCGLHQQHPLTLKWVDSEGDPCTVSSQMELEEAFRLVC
QGRDEVLIIHVFP
>NP_035148.1
RFSFCFSPEPEAEAQAAAGPGPCERLLSRVAVLFPTLRPGGFQAHYRDEDGDLVAFSSDEELTMAMSY
VKDDIFRIYIK
>NP_035970.1
IRIKIPNSGAVDWTVHSGPQLLFRDVLDVIGQVLPEATTTAFEYEDEDGDRITVRSDEEMKAMLSYYY
STVMEQQVNGQLIEPLQIFPRAC
>NP_036077.1
VRIKFEHNGERRIIAFSRPVRYEDVEHKVTTVFGQPLDLHYMNNELSILLKNQDDLDKAIDILDRSSS
MKSLRILLLSQ
>NP_058644.1
VEVKSKFDAEFRRFALPRASVSGFQEFSRLLRAVHQIPGLDVLLGYTDAHGDLLPLTNDDSLHRALAS
GPPPLRLLVQKR
>NP_062652.1
LIIKAQLGEDIRRIPIHNEDITYDELVLMMQRVFRGKLLSNDEVTIKYKDEDGDLITIFDSSDLSFAI
QCSRILKLTLFVN
>NP_067384.2
MEVKSKFGAEFRRFSLERSKPGKFEEFYGLLQHVHKIPNVDVLVGYADIHGDLLPINNDDNYHKAVST
ANPLLRIFIQKK
>NP_115899.1
VEVKSKFGAEFRRFSLDRHKPGKFEDFYKLVVHTHHISNSDVTIGYADVHGDLLPINNDDNFCKAVSS
ANPLLRVFIQKR
>NP_493462.1
TILKARHADVVRKTSLHHANDLTLIDLVLNVQRLLALPSDANFVLKYKDEEGDLVTLAEDSDLLLALH
TSGATLDVTVVVD
>NP_495011.1
KLKTRFQGQVVVLYARPPLILDDFFALLKDACKQHKKQDITVKWIDEDGDPISIDSQMELDEAVRCLN
SSQEAELNIHVFV
>NP_524892.2
ITVKTAYNGQIIITTINKNISYEELCYEIRNICRFPLDQPFTIKWVDEENDPCTISTKMELDEAIRLY
EMNFDSQLVIHVF
>NP_573238.1
EVKSKFDAEFRRWSFKRNEAEQSFDKFASLIEQLHKLTNIQFLILYIDPRDNDLLPINNDDNFGRALK
TARPLLRVIVQR
>NP_593744.1
CKVKVRLGDETFALRVPSDISFEDFCERLTNKLGECEHLSYRDTNANKVLPLNNVDDLRKACSQESGV
LLFAE
>NP_594030.1
SQFTIKYRSIAGRVHRLRLDGINSVSDLRTAVEEREKEQLVTLTYIDDEGDVVELVSDSDLREAILLA
RRRGLPRLE
>NP_594221.1
VKIRLRLHEVSLVLVVAHDITFDELLAKVEHKIKLCGILKQAVPFRVRLKYVDEDGDFITITSDEDVL
MAFETCTFELMDPVHNKGMDTVSLHVVVYF
>NP_660143.1
IRIKIPNSGAVDWTVHSGPQLLFRDVLDVIGQVLPEATTTAFEYEDEDGDRITVRSDEEMKAMLSYYY
STVMEQQVNGQLIEPLQIFPRAC

>NP_956309.2
KAQLGDDIRRIPIHNEDITYDELLLMMQRVFRGQLQSSDEVTIKYKDEDDDLITIFDSSDLSFAIQCS
RILKLTLFVN
>NP_991274.2
LSIQPSYEELLSRMRDVFHVDDIALNYRDAEGDLIRILDDEDVVLMVQESKRTESKVK
>XP_001922991.1
LKVTFRGNAKSFLLSGSETKSWESMEAMVKRSFGLCNLQLTYFDEENEEVSVNSQLEYEEALKSAARQ
GNRLQMNV
>XP_005162724.1
LRVKLEHEREKRIIPFQRPLKFKDLLQKVTEAFGQQMDLYFTEKEMLVALKCQEDLDRAIQGLSSSSG
MNNLLRVILKTP
>XP_005168897.1
VDSPSHLNFNDLLAAIRDAMPEATVTAFEYEDEVGDRITVRSDDELKAMLSYYCN
>XP_005257433.1
RIKFEHNGERRIIAFSRPVKYEDVEHKVTTVFGQPLDLHYMNNELSILLKNQDDLDKAIDILDRSSSM
KSLRILL
>XP_006511208.1
IRIKIPNSGAVDWTVHSGPQLLFRDVLDVIGQVLPEATTTAFEYEDEDGDRITVRSDEEMKAMLSYYY
STVMEQQVNGQLIEPLQIFPRAC
>XP_006522044.1
LIIKAQLGEDIRRIPIHNEDITYDELVLMMQRVFRGKLLSNDEVTIKYKDEDGDLITIFDSSDLSFAI
QCSRILKLTLFVN
>XP_006532493.1
TLNVTFKNETQSFLVSDPENTTWADVEAMVKVSFDLNTIQIKYLDEENEEISINSQGEYEEALKMANI
KQGNQLQMQVHE
>XP_006532495.1
VTLNVTFKNETQSFLVSDPENTTWADVEAMVKVSFDLNTIQIKYLDEENEEISINSQGEYEEALKMAN
IKQGNQLQMQVHEG
>XP_006532496.1
VTLNVTFKNETQSFLVSDPENTTWADVEAMVKVSFDLNTIQIKYLDEENEEISINSQGEYEEALKMAN
IKQGNQLQMQVHEG
>XP_006721966.1
VTLNVTFKNEIQSFLVSDPENTTWADIEAMVKVSFDLNTIQIKYLDEENEEVSINSQGEYEEALKMAV
KQGNQLQMQVHEG
>XP_009294714.1
SCLHCYFLQPEGIETRDICVQEDLSIQPSYEELLSRMRDVFHVDDIALNYRDAEGDLIRILDDEDVVL
MVQESKRTESKVKRPVNQFPWELLVTHA
>XP_009295652.1
RVKLEHEREKRIIPFQRPLKFKDLLQKVTEAFGQQMDLYFTEKEMLVALKCQEDLDRAIQGLSSSSGM
NNLLRV
>XP_009298791.1
KFRVILDHEIKKLCVPEMPRTVKELEATIKQTFEIYVDISLQYKDQDFDDFITLSSTDDLK
>XP_009302121.1
KIKAHYGGDMLISDLDLALTYTEVCKEVREMCGVRKETPITLKWIDDEGDPCTISSQMELEEAFRIYS
RNRHSGLLLHVFP
>XP_009305028.1
VTVKVNFRGNVKKFPVLDTNKAQWETVEAWIKTTFGLSHFQVKYFDEDNEEVCINSQDEYTEALKSAF
KQANQLHMNV
>XP_011241042.1
IRIKIPNSGAVDWTVHSGPQLLFRDVLDVIGQVLPEATTTAFEYEDEDGDRITVRSDEEMKAMLSYYY
STVMEQQVNGQLIEPLQIFPRAC
>XP_011247339.1
VRIKFEHNGERRIIAFSRPVRYEDVEHKVTTVFGQPLDLHYMNNELSILLKNQDDLDKAIDILDRSSS
MKSLRILLLSQ

>XP_011520089.1
IRIKIPNSGAVDWTVHSGPQLLFRDVLDVIGQVLPEATTTAFEYEDEDGDRITVRSDEEMKAMLSYYY
STVMEQQVNGQLIEPLQIFPRAC
>XP_011540075.1
RLKAHYGGDIFITSVDAATTFEELCEEVRDMCRLHQQHPLTLKWVDSEGDPCTVSSQMELEEAFRLAR
QCRDEGLIIHVFP
>XP_016878750.1
PDSIVEFDAEFRRFALPRASVSGFQEFSRLLRAVHQIPGLDVLLGYTDAHGDLLPLTNDDSLHRALAS
GPPPLRLLVQKR
>XP_016884297.1
NWLRCYYYEDTISTIKDIAVEEDLSSTPLLKDLLELTRREFQREDIALNYRDAEGDLVRLLSDEDVAL
MVRQARGLPSQKRLFPWKLHITQK
>XP_635325.1
ITYKSNFEGDVRRFSSDHPLTYTRLQDKLVNLYNLYEISFGITYLDDGDNITIADAKDLEEAHNLLGN
EILRLTITRK
>XP_638776.1
TTLNVTLGSEMKSITVPKSSTYKDMISTIKDKFGVNSKSTLCIKCENKDGEMFSLASDCHVKKAYNQQ
PENQPKELRLVVKEI
>XP_639165.2
IRIKCILGDDIRIIKFNSNISYGGLMNQLEQDFQCPISIHQYEDYEGDKVTVKSKDDIMEALTMYFEL
KALNPTKIISTKF
>XP_644298.1
ILIKSELENDKRRFRLKECSFSCLCYTLASIYSFYNDMIYSIFYLDNENEWITLASTDDLKESYSLCP
SLIRIKIIVLDLT
>XP_646538.1
LILKIQHNDDTRRVSMERDPTFLELRKMTVTFFKINSFLIKYFDEDKDLITITSDNDLKEAFSIATTS
PRTVRLFVSKT
>Amoebozoa_A0A0A1UD85
FKLEYMNDIRVVDLSTLDLNSLYTAFQNKFNLTNMRLRPRFLLKYLDAEGDWVTLCDDDDLVLALKQA
NKYTLRLKLVD
>Amoebozoa_A0A151ZAG5
NDNKSVSIPKNSSYKDLTTTIKDMFGMNNKSTLCIKYENKNGDMLSVATDCQLKKAYRQTPSADGGDS
ELRLVVKE
>Amoebozoa_A0A151ZC01
KVQYGDDTRRVSLDREPTYSELKKMTMNFFNLQENNFQIKYFDDEGDKITVTSDIEIREAFNFARKKT
PALLKLFI
>Amoebozoa_A0A151ZC78
ILIKSELGEDKRRFRLKECSFSCLCYNLTSLYQLSNDSIYSILYLDNENEWITLASTDDLKEAYSISP
LLIKIKIN
>Amoebozoa_A0A151ZHK7
TDRRLIQLTYPYTYQQLKQKIELKFEISLRNITIKYQSSDEQYYPITNQGDLDYAISY
>Amoebozoa_A0A2P6N9K2
IVFKIRLTNEIRRLSVPSTISFSHLLNLIAHAFADPNLTQTHLLRYKDDEQDWVTMSSDLELREAIRI
YKKDPSVPLQLQQS
>Amoebozoa_A0A2P6NEE1
LTVKVTHKDQVRRSTIPVHTPYSEFKSRIGQLVGVKHELFNLQYEDDEGDLISLTNDEDLREAYNVGS
KSTSLRLRTILL
>Amoebozoa_A0A2P6NIU0
FSCHYKDKRYVRLLSHKDKTVSLLDLNQFIEEKFGISNLIIRYKDDENDLITIVKQEDLDIAILDF
>Amoebozoa_A0A2P6NJ77
LIINATVLGEQEKPIENRISSIIAKDNEITFDAIVENLSKTFQYLNTEGHTISLKWRDEEQDLITITS
DPELQEALSHCQTIGCHWLLRLFLVVR
>Amoebozoa_B1N2I4
TLFKLEYMNDIRIVELQNLTIVSLYDAFTKKFNLKDVRLRPRYLMKFLDTDGDWVTLTDDTDILLAMK

QMKNITLRLKLVDS
>Amoebozoa_D3B745
LSIKVTYNGDTRRISFERTPSYLELFSTLKSLFSLTSLLLKYTDDDGDLISVTSEVEFKEALSLIGYK
TPRLLRLTVLAK
>Amoebozoa_D3B9M1
YTFKVCFEGDVRRFSFEGQPAFASLSQKLTELFSLVESFVITYLDEVGDNITISSDEELEEAFIFLQE
TQQSILRLSVVPK
>Amoebozoa_D3BCZ9
TVIKCKLYDQESYKKVDISLYNGFIPLRETLSAMFSIGHSFVICYQDSENDIIEINDQEDDWFDFYSI
PVWKPAS
>Amoebozoa_D3BP85
VRIKCILGGDIRIIRVSVNVEYEALLMQLESEFGQSVQVYQYEDYEGDLITVRGKSDLMEAIYICIEL
KNKPISLRFFLKP
>Amoebozoa_F0ZEH2
LVLKIQYGDDTRRISMEREPTFLELKKMTITFFKLSGFLIKYYDEDNDLITITSDSDLKEAFTVKRNP
PNVLKLFVSKS
>Amoebozoa_F0ZG42
YKSNFEGDIRRFSSDSELSYQRLKDKVSNLYSLYDIEFIITYLDDGDNITIADNKDIEEAYKTIIDNE
ILRLTVTR
>Amoebozoa_F0ZVT4
LRIKCILGDDIRIIKVNANVSYDGLMNQLEQDFNTPILIHQYEDYEGDKVTVKSEYDIMEAISMYFEL
KSLNPSKIISTKFFLKQG
>Amoebozoa_F0ZY34
VTLKIFYIDRRLIQVPVPCTISTLKQKIELKFELKLDNQIEISFILDGQKELLNNQVQLDKMLCMEVN
EVHLRDR
>Amoebozoa_F4PHK7
LFKSQFGEDKRRFRIKEPSFPCLCYTISSLYCFDNNDFIYTIFYLDDENEWITLSSTEDLKEAFTISP
KIVRLLVNKL
>Amoebozoa_F4PJ30
RIKCKLGEDDIRIIKVPSTVTYVELMDQLEEDFGQPVAVHQFEDHEGDRVTVRGKNDIIDAINMFDEF
RSMYP
>Amoebozoa_F4PSP5
LIIKATHQGDNRRVSFERDPTFEELYGTLKNLFKIDPFNITYTDEDGDQITLSSEMEMKEAISMISAN
KPRLLRLAVTKP
>Amoebozoa_K2H6P4
TLFKLEYMNDIRIVELQNLTIVSLYDAFTKKFNLKDVRLRPRYLMKFLDTDGDWVTLTDDTDILLAMK
QMKNITLRLKLVDS
>Amoebozoa_L8GDU9
KAFYNEEIRRLVLPEPSWDHLKQRLAELFHFPSTSSFQVKWKDDEDDYITLDSDEELAQALRYTDPQQ
LLRLFIF
>Amoebozoa_L8GF76
WVKLVFDDDISILRFKRRTLTVKRLQREINRIHQCQPRLRWIDSDGDCIEIKTTEFLNYALEDWQRR
>Amoebozoa_L8GGA9
VVIKFQYGNDVRRITLPAEGITYEQLLSLIESLYSQAGPGDHPALPQEFKLKYADEENDFIGLTCDNE
LRDALSLLPEGKVLRIMIVDV
>Amoebozoa_L8GGR6
KCLYNKETRCVRIREGSSYKDLQKQLEKEFKFVPVIKYKDPEGDKVSMKKQGDFEYALSLINGNALRL
YI
>Amoebozoa_L8GKF5
ITLKCVFKDDISLLHVSPLVEFAELGKKIKKEYGRMMEIKYKDAEGDLIRVKTTKGLRSAIKNWSGEG
SLKLLLSEK
>Amoebozoa_L8GMM9
VLIKAYLGNDARRVTVEDDIPFADLRAMLMKLFAVSGSGEPITPPAPTQFWIRYKDDEDDLVSISTDS
DLREAIAYSASNADLLRLYIQQG

>Amoebozoa_L8GNA6
VLVKCYHGDDIRVVEVPRDVRYQDFLSELQGKYVGPLTVTYKDEEGESITIDSQGSLETAIHKHLKEA
AKTLKLRISIR
>Amoebozoa_L8GNC1
IRCKLSFNGELRCCALALPPSWDALQADVARLFNIGPAVGRAFVLKYVDDEGDSVTFDSPLELHDAVH
AAAQATTPNPCLKLLVQLV
>Amoebozoa_L8GNI9
HFVELRDTIRTAYGRNLRVKYSDRDGDTINIDSDASLNYAFNDWRGQIGLVDHHHQKATTSWRLHLHD
>Amoebozoa_L8GQZ1
GSRQLQYADLVEEVARVHGSVATKKMKIRYTDGEGDEVTILGDDALRYAFRDWR
>Amoebozoa_L8GRV3
TRTVVTASASFAELERQVRKSFGLSERHQLDLRYIDDEQDRVCVSSDEELRYALNSFQDKSLHLFL
>Amoebozoa_L8GTK3
LKCFYGEDVTVLRVKRDITFTKLNSKLKAEYKKDMEVAYKDSDGDVIPITKSAHLKAAMKEADGRAVR
LQLK
>Amoebozoa_L8GY70
TRLKCFFGEEISVLTVPSDIKYRHLKAKIEEIFGEGVAIHKYEDHVGDLITVHSKHDVREAFTLYAKA
IRKQSKRDAVEPYLKLHLARE
>Amoebozoa_L8GYB1
LDAHRPLLYADLLREIERVHGPLVAAAASGEVGVRVRYEDKDGDVVNVHSDDGLCYAY
>Amoebozoa_L8GZZ7
VAIKCVVDDDISLVRLNLETDLSLSGLHRAIERVYGQRMRAHYFDSEGDKIRITTDSSLRYAYEDWCQ
ALERSGATISWRLHLKPR
>Amoebozoa_L8H6B1
VHFIPPPRYTDLVELMRTSPLRIPSTALIQFRDSDGDLLPLRHQDDLNYFLDNHKRSHQPTAISGLSL
FV
>Amoebozoa_L8H883
MTIVATNENRFIALPSPPAFALLVEKISTVFDQVNKAAVRIHYVDDESDRVAVSSSDELACALSLLPA
GRQLRLFVS
>Amoebozoa_L8HI52
KKAESTYHDDISLLRLDARRPLLYADLMAEVHRLHGPLLLAILMQEGAKGVVRISYRDVEGDLVGIQS
DDALRYAYEDWRKQARRRPR
>Amoebozoa_L8HJM7
SLKVKLDEDIRRFSVPKGITFAELHSLVSKHFVLAPENVQLKYIDDEEELITLGSDLELQEAKRLQPT
VLRLN
>Amoebozoa_L8HKC2
VVLKVTYGSEIRRITVDDTQAFSYKDLRVLLKKLHHNTLPNHFEIKYLDDENDKVTISSDRELADALQ
FVKTAKQPLLRIMLSDP
>Amoebozoa_Q54Q21
TTLNVTLGSEMKSITVPKSSTYKDMISTIKDKFGVNSKSTLCIKCENKDGEMFSLASDCHVKKAYNQQ
PENQPKELRLVVKEI
>Amoebozoa_Q867T7
ITLKVFYKDRRLIQIPVPCNLSTFIQKIELKFEITISDKFSLSFQLDGEENEINSQVQLDKMICMEIN
EINVKDI
>Apusozoa_A0A0L0D6J2
LRVKLVYVGDDGGSVRVHAMSLPHDVHFDEVLQTASAGAGLDAHVALSYVDEDGDVVGLASDLDWAAA
LDWFMAQAEVRTLRIQVHVV
>Apusozoa_A0A0L0DC02
VVLKCKFKGAFKLLRIERTDPFTKLQTQMDEAYKTAVYLSWQRADGTLIDVINADDFAYMLDEHASKK
KIDLIVQ
>Apusozoa_A0A0L0DJ90
RVVKITFGAETRRFRLNPETTTFTRLRELVKQSFAPLAGDATEFVLKWTDEFSEVNVLGSNLELTEAF
HVAGTALNPPILRLMVEAS
>Apusozoa_A0A0L0DL51

LKVVRGQDRKLLTVARTVQLAELFAKLGAEYATVIQEFKFKDAEGDMITVRSSSDLQEAFHEYDL
>Apusozoa_A0A0L0DM96
TKIKLYFGDDVRKVVLMAPFSFGELLEGVEARYGMSGLQLAYKDEDGDIIVMVDDGDIPELMEQEKIA
VYVSQ
>Apusozoa_A0A0L0DQI7
YFFKFKHGGSTFRASASGESYIELIESVRTRIKAPRGAGIELAYKDDEDDDIELGSDSELRDAVQLAE
ELGWRTIKLSVVVR
>Apusozoa_A0A0L0DVD6
VKVKCHFHDTRVIVVTSDIPFTELEQAVFRKFNENNLILKFRDEDGDLVTMSCQDDLDLALEENPKLE
LFIDQ
>Apusozoa_A0A0L0DVU4
IRLKCKFGDEMRLVSVTPSMAFDDVAAKMEAMFGVACLMQYSADDDTITIRSDEDLREAVEFVADSGS
TTLKVFLT
>Cryptophyta_L1JV30
LPLKATFSSQDADSVIRRLSVPVELDEEEVPVEGFSAVTDALVYAFRNELKMGCDDVKISYTDEDGDE
VMISCDEELGLAMQQFRDASVMRVKLHA
>Cryptophyta_L1K2L0
LKFKVSLTQPGKQAVFRRLKWNFEFDHFGNPVQGYEMIMTGLRNAYREELLGVPTSALQLSYVDEDGD
NILISSDSELGMLLQDKSQSAGIRLYLHAL
>Euglenozoa_A0A0N0DU41
ARLKIHFDAQTRVMPVADAAQAKFTDVYDQLHAHCQAQLAQSSCSGAAVAVGTSSSPEERRLRIRYED
VDGDFISLLNQQDWDVMLSELLSQEENGSGSSSGGGKIELFCDYP
>Euglenozoa_A0A0S4J229
VSFKIVLQDKKRPETKHRFRGDFTYAQLTDLLSHLWGGKKKLVVEYTDEDGDDILVLTEVEWQECVRL
HLKRLEESR
>Euglenozoa_A0A1X0NI49
LKLKVHFEAQLRILRVPEPENITFEQIYEKVHAFCMQQAAVLPRPVGQRLRIRYRDHDGDSISLITQD
DWNAFLVEEVPGGLSGAKLELFCDFP
>Euglenozoa_E9AVC0
LRLKIHFDAQTRVLPVADAAQASFHNVYKQIYEFCQTQPQRAAGAERRLRIRYEDAEGDFISLLDQQD
WHMMLSELAPHGCGGVKIELFCDYP
>Euglenozoa_E9BFE6
RLKIHFDAQTRVLPVADAAQASFHDVYKQIHDFCQTQLQQPQRAAGAERRLRIRYEDTEGDFISLLDQ
QDWHMMLSELAPRGCGGVKIELF
>Euglenozoa_Q4DB71
LKVHFDAQLRVVLIANPENATFDEIYKKVSELCSRQEGGGASNSAGRKLRLRYQDSDGDCISLLTQED
WDVF
>Euglenozoa_S9VA21
QRTIEFNDLRKLVLERWELDINGPSQLYYKDDEGDRIKIDSSEFKRLLQRHFATSDIVRIHID
>Euglenozoa_W6KL49
LKLKVHFEAQTRIVHVHEADKTSFEDVSKRLYKQCCELFAIMHRSQSSFPPDPRRLRIRYEDTDGDFI
TLLNDEDWRVMLSELRLSSNLRLSCSGG
>Heterolobosea_D2UX43
KFKVYRGSDIYLITQKRSQLSFTSLKLEIIERYQCVKPTILYKTRDGDFIIINRDEDLARVISSYHQ
>Heterolobosea_D2V5H2
FKTIYKAEMRVFAKVPRDVTYLELKETISKSYGFPVVLKYVELDDEKDEITIGSDRELRDFIECFEEM
GKGLRLEL
>Heterolobosea_D2V7N4
KEEKPKSVHRLILDQSGITFEEFVQKLKTKSIVTDTSFKLFYLDDQNDWVSLSDEEDWKILNSILASE
TVIKIVCQNI
>Heterolobosea_D2VCK7
LSVKVKLNDTIRRFSVPSSCTLIEFYQQVSKAFQLNIDITTHLVQYKDNEDDLIVTTSEDEWNYAKKS
FSTPCMQVTLKEK
>Heterolobosea_D2VL54

VIIKLYLSKEHIRRFTIPKIASYEDLKSRIIQFYDEVGQTINPSEDLDLQYCDDDNEWTVLRNEDDWY
CCKAICPTNIKIRKPSMWMKAQ
>Heterolobosea_D2VP18
KVKLSYKEETTIVSVFLGMSMDGLFEIISSHYNNFLACDNLHDKINIKYADDEDKNELISIRTQTDLD
HFVES
>Heterolobosea_D2VT71
LVFKCYHAEDVHLIALKRDRATFSELKLKIIEEYNCLKPIIKYRDTEGDFINIRKEEDFNICKQNVPV
DSAIRLYITTE
>Heterolobosea_D2W1M7
KEEKPKSVHRLILDQSGITFEEFVQKLKTKSIVTDTNFKLFYLDDQNDWVSLSDEEDWKILNSILASE
TVIKIVCQNI
>Heterolobosea_D2W469
IMIKCYDSLENQSDATILRFSPEDELTLQKVKMEVLSRKPILSSTNQIQLKYKDEDDDIITLKKEDDF
KLCLANYRNSGQRYLKLYISSN

>g17664_CHBRA211g00240
LFCKIFRGGELVGRAVDLGKFSNYDQLCAELSVMYGIDRHELQKNMAYRDSEGHWVLVGDEPYRHFAG
TARKAMIMST
>g19475_CHBRA233g00310
LVIKVQYQTNLRRFIVPRPPASKFADVKARIREALSIADGVEFSLTYRDKDDDLVTMAGDDDLYDAFT
IQNLNPLRVFVTVE
>g19697_CHBRA238g00220
AHITAIFEGWPIGRRVNLREHSSYEGLSKTLREMFKARIAEEKWTCSKAGGGGGGGGGGEGETCNSLN
ALPGHVIAYEDADGDLMLAGDIPWEQFIVTAKKIKIVKT
>g34700_CHBRA42g00490
VPLKLYYGHDIRLAEMPVNAGFGELRDVVRKRFPSSKAVLIKYRDSEGDLVTITSRQELRLAEACASV
ASSVATASSTAPPLSSVGE
>g38148_CHBRA490g00130
VTVKVTFGQDTARFRLPSGHCFDDLQQEVAQRLKLDTTTLVMKYLDDEGEYVLLSSNEDLMECIDVSR
NAGNNTIKLTARCE
>g39692_CHBRA529g00140
AHITAIMEGWPIGRRISLREHYNYEGLSTTLRHMFEFMIPAEQRYWIRPGCHPPCNSLNALPGHVIAY
EDADGDLMLAGDIPWEHFVTTAKRIKIVQT
>g40427_CHBRA554g00190
FTFKIEDKTGRMHRFTCGIESLDELVMAVANRTGLDMKSTNAPHIMYEDDEGDMVLLNNDSDLIAAVN
LARASGWKGIRLYLDVD
>g53552_CHBRA981g00020
NRLTYVGGDTRIQTVNRDVSYGELMKKLTDFYGQPLILKYQLSNEGLDTLVTVSCEEDLVNMMEEYDA
EVRSERTAARLKVFLFP
>g53553_CHBRA981g00030
NRLTYVGGDTRIQTVNRDVSYAELMKKLTDFYGQPLILKYQLSNEGLDTLVTVSCEEDLVNMMEEYDA
EVRSERTGARLKVFLFP
>kfl00098_0270
IPLKLVYQGHDIRKTELPANGGLKTLREVVQKRYPGSKAVLVKYQDEDGDWITITSNEELRGALELCG
YAGKSADAKNPEEGAKKGPVRLEITEV
>kfl00358_0060
TVFTFKVEDNLGRKHRFSCGCQSMAELVVALSSRLDLPPAAIPPISYIDDEGDRVLLSGDGDLSSAVN
VARTAGLKNLRLYLEFA
>kfl00615_0010
SVVKVSFKGIMKRLTYGTEEANFDRLVKTLKESFSIAEGAKLRILYTDSDGDKVLLSDNEDLKDALKN
QKLNPLYLEVTIIKEAKS
>kfl00667_0030
DGKLRYVGGETRIFTVSRDITYSELMFKLTEYYSEALSLKYQLPDMDLDTLLSVSNDEDLNSMMEEYD

EIERNSAEGKAQRLRLFLFP

>CDC24_YEAST
IPESSILFRISYNNNSNNTSSSEIFTLLVEKVWNFDDLIMAINSKISNTHNNNISPITKIKYQDEDGD
FVVLGSDEDWNVAKEMLAENNEKFLNIRLY
>BEM1_YEAST
AQSTSGLKTTKIKFYYKDDIFALMLKGDTTYKELRSKIAPRIDTDNFKLQTKLFDGSGEEIKTDSQVS
NIIQAKLKISVHDI
>NoxR_H6CB14_EXODN
GSNRPEMRKIRVKCHFNEDTRYLMIGAVIDFGDFESKIREKFSIKDPMKIRMQDDGDMITMGDQDDLE
MLLSSVRNQARKERNEMGKMEVWISSSRLSSMS

>Fungi_Basidiomycota|Pucstr1|23979|evm.model.scaffold_6.94|3_81
HQAPLSIKIKHATVIHKVQVNQHVPIWSTFLAAIAQRFGMPEEQPIGLQYLDPEGDTITISTQADCDE
LWHKVLSVTA
>Animalia_Echinoderm_tr|W4XXL6|W4XXL6_STRPU|5_91
DGMIPIKAAFNGSILCIRIDPNVTLEDFLQDMRGICNFPEKQNFTVKWLDEDGDPCTISSQEELEEAI
RLYELNKDSNINIHGNYI
>Fungi_Mucoromycota|RhiirA1_1|414327|
fgenesh1_kg.219_#_22_#_remain_c11464|59_123
FSIVITSVKPTWDQLSTKIRSQFNIPANFKFGLTYLNSDNNEIIISSQKELNNYFFQHKDDELS
>Fungi_Basidiomycota|Pucstr1|4115|evm.model.scaffold_13.89|3_82
HQAPLSIKIKHATVIHKLQVNQHVPIWSTFSAAIAQRFGMPEEQPIGLQYLDPEGDTITISTQADCDE
LWHKVLSVTAI
>Protozoa_Amoebozoa_Entamoeba_tr|B1N2I4|B1N2I4_ENTHI|1_88
TLFKLEYMNDIRIVELQNLTIVSLYDAFTKKFNLKDVRLRPRYLMKFLDTDGDWVTLTDDTDILLAMK
QMKNITLRLKLVDSYQTAT
>Fungi_Mucoromycota|RhiirA1_1|419773|
fgenesh1_kg.493_#_1_#_ACTTGA_L001_R1_(paired)_contig_17190|50_129
MRTSTFKISNQAITRRFTLSIEQPTWLELELKVREIFSIPSSVSPGLTYIDEGDDNITISSQIELEDY
YKQVKCHEYSD
>Fungi_Mucoromycota|Morel2|829797|Morel1.CE669783_1526|12_103
LGDKPVIIKFKCAGTNLRVPISQVPSYNELCFIVQRLFRSDVSSDLDNLVLRYEDEDGDLITIQDDAD
ISHAISLSSLLKLTVNDKITHPV
>Chromista_Stramenopile_Phytophthora_tr|D0P0V1|D0P0V1_PHYIT|1_78
ASQQVALKIAFGGDIHRTRVELRCLSLAELTQLMAQTFRLPVGDFVVQYRDPEGDSVNVTTDAEFQEA
LRVFLSSSE
>Chromista_Hacrobia_tr|L1IAZ6|L1IAZ6_GUITC|164_241
LEVDGEGRVVRGHGALVRALVEEYELVLGCCEESVVVSYEDSDGDRVEMRSDEDLAQAMESYPFGYVR
VYVQGRGGG
>Chromista_Hacrobia_tr|L1JXV3|L1JXV3_GUITC|21_86
LEGPPPDTMLELEEKTRHALGYGAGAKFCFKWIDEEEDMIMLGSDEELKESLSSVSKSQRKLVFV
>Protozoa_Choanozoa_Monosiga_tr|A9UPR4|A9UPR4_MONBE|3_99
TRASTVILKVRLGNDIRKIPIHNADLTYDDLVLMLQRVFGPTGRLPKDDFTIKYEDEADAAVSSIDSA
SIDNIHRQLQNVCVSVNDLVRRLDQVAG
>Fungi_Mucoromycota|Morel2|31788|Morel1.estExt_Genemark1.C_270029|
1_90
AATVKAWHRDDCRRLSIDIEQIQYQDLKKKLQNTFQLENEFTVKYADLDGDLCTIAGTEEFEDMINSE
LVKERKATLRLQVIPNSRRPS
>Chromista_Hacrobia_tr|L1IKT8|L1IKT8_GUITC|236_296
EPFQGYRKLVEAVIKEFGLDVAFADTMRLFYQDADGDNIIVSSDHELGIAMKHFPAGLYV
>Plantae_Rhodophyta_Chondrus_tr|R7QD18|R7QD18_CHOCR|218_308

GMSSLYAVKAEYRREIHRLTISRSASFLELRRKMEEVFQLSEKVCLAYRDNEDDYITISTESDMRELF
GLADKYSLNPIRVRLVERIEKR
>Animalia_Echinoderm_tr|W4Y123|W4Y123_STRPU|2_95
AQPLVIRIKTQHGEDVDWTVQLGQLSFHEVLEVIAQVLQDNTPAAFEYEDEDGDRITVRSDDEMEAMI
NYYLSLLAECEARRVMAPPLNIYPQ
>Animalia_Nematode_sp|Q9NAN2|PAR6_CAEEL|10_100
HHSTLQVKSKFDSEWRRFSIPMHSASGVSYDGFRSLVEKLHHLESVQFTLCYNSTGGDLLPITNDDNL
RKSFESARPLLRLLIQRRGESW
>Fungi_Mucoromycota|RhiirA1_1|511591|MIX24024_34_34|227_310
MKEDEILLKCHVGRDIRILNFILPIRYEEIKRKIQEAFGIAEISKLKYKDEDGEFITIIDNVDFDTAS
KLYKRENKLEIWVIE
>Chromista_Stramenopile_Phytophthora_tr|D0N7Q1|D0N7Q1_PHYIT|2_76
AKQTALKINYKGELHRLRVDLSTFSLEALTALFDETFNLSPGSFVVQYTDADGDCLNVTSQAEYEEAC
RVFLSG
>Chromista_Hacrobia_tr|L1IEZ2|L1IEZ2_GUITC|211_282
PVGGYRMVMEGLQSAFSEDLKGIRTSCLQLSYIDDDGDKILINSDQELGILLQDKPQAARIRLYLHAV
SGS
>Fungi_Chytridiomycota|Rhihy1|717761|
fgenesh1_kg.26_#_47_#_Locus5745v1rpkm39.22|1_91
STSETVIKISYNGSLRNVPLTSEEQISWDAFETKIRTLFTIPDSTALTVNYVDSDGDRIIFDTDQELA
ELFRGVGRPSKLFVTGKDSTPV
>Animalia_Frog_tr|A0A1L8GH71|A0A1L8GH71_XENLA|229_331
EQYQVSLLRCYFHDHDICLIRDISLEEDVGKCPSYKELLDLIRNQFPDAEVALNMRDTDGEMIRLMDN
SEMELLITRGKRTPRAKNYFPWELHVTHKDDFEV
>Animalia_Frog_tr|Q3SYN3|Q3SYN3_XENLA|233_329
VSLLRCYFHDHDRCLIRDISLEEDVRKCPSYKNLLGLIRNQFPDAEVALNMRDTDGELIRLMDNSDME
LLITRGKLTPRAKNYFPWELHVTHQDDL
>Animalia_Nematode_tr|Q9GRV1|Q9GRV1_CAEEL|134_214
TTSTVLKARHVDFVHKTSIHHTNDLTLIDLVLNVQWRLALPSDANFVLKYKNKQGDLVTLVVDSDLLM
VLHTSGATFDVT
>Fungi_Mucoromycota|RhiirA1_1|499194|MIX11627_521_96|1_83
ANIKVTYGSISRRLTISSTTTWSEIETQFRNLFNIPKEFQIIVSYTDDEGDVITLSSDLELQEVLSNN
SIKFILTTSIMNNL
>Animalia_Dog_tr|E2RIM1|E2RIM1_CANLF|232_334
EEDPTNWLRCYYYEDTISTTKDIAVEEELSSTPLFKDLMQLMRREFQREDIALNYRDAQGDLVRLLSD
EDVGLMVKQAQGLPSQKHLFPWKLHITQKDDYRV
>Animalia_Human_sp|Q15080|NCF4_HUMAN|232_333
EDDPTNWLRCYYYEDTISTIKDIAVEEDLSSTPLLKDLLELTRREFQREDIALNYRDAEGDLVRLLSD
EDVALMVRQARGLPSQKRLFPWKLHITQKDNYR
>Animalia_Mouse_sp|P97369|NCF4_MOUSE|232_334
DEDTTNWLRCYFYEDTGKTIKDIAVEEDLSSTPLFKDLLALMRREFQREDIALSYQDAEGDLVRLLSD
EDVGLMVKQARGLPSQKRLFPWKLHVTQKDNYSV
>Animalia_Pig_tr|K7GN17|K7GN17_PIG|234_335
EEDPTNWLRCCFYEDSVSTTKDIAVEEDLTSTPLYKDLLELVRREFQREDLALNYRDAEGDLVRLLSD
EDVQLMVRQAQGLPSQKRLFPWKLHITQKDNYK
>Animalia_Pig_tr|A0A286ZZ01|A0A286ZZ01_PIG|11_99
PDSIVEVKSKFDAEFRRFALPRASVSGFQEFSRLLRAVHQIPGLDVLLGYTDAHGDLLPLTNDDSLHR
ALASGPPPLRLLVQKRAEAD
>Animalia_Mouse_sp|Q9Z101|PAR6A_MOUSE|10_100
SPDSIVEVKSKFDAEFRRFALPRTSVRGFQEFSRLLCVVHQIPGLDVLLGYTDAHGDLLPLTNDDSLH
RALASGPPPLRLLVQKRAEGDS
>Animalia_Human_sp|Q9NPB6|PAR6A_HUMAN|11_99
PDSIVEVKSKFDAEFRRFALPRASVSGFQEFSRLLRAVHQIPGLDVLLGYTDAHGDLLPLTNDDSLHR
ALASGPPPLRLLVQKRAEAD

```
>Animalia_Chicken_tr|A0A1D5NVK9|A0A1D5NVK9_CHICK|232_334
QEDTVNKIRCYYYDETVSTIRDISVEENLSSIPLFKDLMELIKQEFDQHDIVLNYRDLDGDLIRLLSD
QDVELMVSQSRKRSSEKHFFPWKLHITHKDDFSV
>Animalia_FruitFly_tr|O97111|O97111_DROME|14_106
SDTNLIEVKSKFDAEFRRWSFKRNEAEQSFDKFASLIEQLHKLTNIQFLILYIDPRDNDLLPINNDDN
FGRALKTARPLLRVIVQRKDDLNE
>Animalia_Echinoderm_tr|W4XKQ8|W4XKQ8_STRPU|1_84
FIFQLDAEFRRFTINPNKVGTYEDFYAFLERMHHLGDVPFLVGYTDPQGDLLPINNDDNYLKALTSSK
PPLKIVLQKRDEVSE
>Animalia_ZebraFish_tr|Q1LUW2|Q1LUW2_DANRE|244_350
AQGSYSCLHCYFLQPEGIETRDICVQEDLSIQPSYEELLSRMRDVFHVDDIALNYRDAEGDLIRILDD
EDVVLMVQESKRTESKVKRPVNQFPWELLVTHAKDLTV
>Animalia_Chicken_tr|F1P0U8|F1P0U8_CHICK|11_99
AEPVIEVKSKFDAEFRRFAMKRSSAGTFQDFYRLLQSVHQIPCVDVLLGYTDVHGDLLPINNDDNYHK
ALSSANPLLRVIIQKKAESD
>Chromista_Stramenopile_Phytophthora_tr|D0MRD9|D0MRD9_PHYIT|1_106
TENVVLKLSYSGETHRVTVPLRAVDPKKDLSYELVLAKVRETFPRLSSNLRWTLVYRDDEGDVVTLSH
AFEFDEACHVLLAMTPENDDKLRTLHFCVLQRVSFRE
>Animalia_Pig_tr|I3LSR2|I3LSR2_PIG|14_102
DCSAVEVKSKFGAEFRRFSLDRQKPGKFEDFYKLVLHTHHISNTEVTIGYADVHGDLLPINNDDNFCK
AVSSANPLLRVFIQKREEAE
>Fungi_Mucoromycota|RhiirA1_1|452740|gm1.5221_g|1_90
ASIKVTFGQTSRKFTIPSNTTWSQFESQLHDLFNIPSDTSFSISYIDEDGDVITLSTDTELQQILSDQ
ESFGTNVKFNIYTSENSDNND
>Animalia_Chicken_tr|Q0PVE4|Q0PVE4_CHICK|11_100
RGLGTMEVKSKFGAEFRRFSLERSKPGKFEEFYGLLQHVHKIPNVDVLVGYTDVHGDLLPINNDDNYH
KAVSTANPLLRIFIQRKEDAD
>Animalia_Mouse_sp|Q9JK83|PAR6B_MOUSE|11_101
GCLGTMEVKSKFGAEFRRFSLERSKPGKFEEFYGLLQHVHKIPNVDVLVGYADIHGDLLPINNDDNYH
KAVSTANPLLRIFIQKKEEADY
>Animalia_Human_sp|Q9BYG5|PAR6B_HUMAN|11_101
GCLGTMEVKSKFGAEFRRFSLERSKPGKFEEFYGLLQHVHKIPNVDVLVGYADIHGDLLPINNDDNYH
KAVSTANPLLRIFIQKKEEADY
>Animalia_Dog_tr|F1PJN8|F1PJN8_CANLF|13_103
RFGDCLKIRGFFGAEFRRFSLDRHKPGKFEDFYKLVVHTHHIANTDVTIGYADVHGDLLPINNDDNFC
KAVSSANPLLRVFIQKREEAEH
>Plantae_Rhodophyta_Chondrus_tr|R7Q6J1|R7Q6J1_CHOCR|283_375
SVENLVNVKLSFILENEEGCRRWGMHPSDSLSSLLNSIAKKLNCQEVVDLEYIDDDEDTIALKNDEDV
KEMFSIVKKSRIDPLRMNTTLKQD
>Animalia_Frog_tr|A0A1L8ESP6|A0A1L8ESP6_XENLA|12_101
RSLGTVEVKSKFGAEFRRFALEKTKPGKFDEFYGLLQHVHKIPNVEVLVGYADIHGDLLPINNDDNYL
KAITTANPLLRIFLQRKEEAD
>Animalia_Human_sp|Q9BYG4|PAR6G_HUMAN|14_102
DCSAVEVKSKFGAEFRRFSLDRHKPGKFEDFYKLVVHTHHISNSDVTIGYADVHGDLLPINNDDNFCK
AVSSANPLLRVFIQKREEAE
>Animalia_Frog_tr|A0A1L8ELH2|A0A1L8ELH2_XENLA|12_102
RSLGTVEVKSKFGAEFRRFALEKTKPGKFEEFYGLLQHVHKIPNVEVLVGYADIHGDLLPINNDDNYL
KAMTTANPLLRIFLQRKEEADY
>Animalia_ZebraFish_tr|F1R754|F1R754_DANRE|10_100
RTLNAVEVKSKFGAEFRRFSLDRSKPGRFDEFYGLLQHVHRIPNVDLLVGYADVHGDLLPINNDDNYH
KAISMATPLLRLFLQRKEEADY
>Fungi_Mucoromycota|RhiirA1_1|528749|estExt_Genemark1.C_210064|1_84
TSIKVAYQSTVRRFPIPNNITWLDLESKIQELFSLPSTSKFSLSYTDDEGDVIILSTDLELQELLSSN
SMLKFVLVPDEQQPL
```

```
>Animalia_Mouse_sp|Q9JK84|PAR6G_MOUSE|13_103
YDCSAVEVKSKFGAEFRRFSLDRHKPGKFEDFYQLVVHTHHISNTEVTIGYADVHGDLLPINNDDNFC
KAVSSANPLLRVFIQKREEADH
>Animalia_Chicken_tr|E1BY76|E1BY76_CHICK|13_103
LECSAVEVKSKFGAEFRRFSLDRYKPGKFEDFYKLILHIHHIANLEVMIGYADVHGDLLPINNDDNFF
KAVSSAHPLLRVFIQRQDEVDY
>Animalia_Frog_tr|A0A1L8FY89|A0A1L8FY89_XENLA|13_103
VASSSVEVKSKYGAEFRRFSLNRYKPGTFDEFYNLILHIHNISSMDVMLGYADVHGDLLPINNDDNFF
KAVSSANPLLRVFIQKQEEVDY
>Animalia_Frog_tr|Q6GPA4|Q6GPA4_XENLA|13_103
VASSSVEVKSKYGAEFRRFCLNRYKPGTFDEFYNLILHIHHISSMDVMLGYADVHGDLLPINNDDNFL
KAVSSANPLLRVFIQKQEEVDY
>Animalia_ZebraFish_tr|A0A2R8Q7H2|A0A2R8Q7H2_DANRE|10_99
NEESVVEVKSKFEGEYRRFALKKNTGGFQEFYQLLQTIHRIPGVDVLLGYADIHGDLLPINNDYNFHK
ALSSANPLLRIIVQKRDVDDT
>Fungi_Mucoromycota|Morel2|524347|Morel1.CE364333_5140|6_81
LSPCKANFNGSLRRFLISRPAVWSDFEHKLRNVYSLPASAAIDVQYKDEEGDVISLNTDSELEDVLAT
HALFTQI
>Chromista_Hacrobia_tr|L1IE99|L1IE99_GUITC|273_339
RMVMEGLQSAFSEDLKGLPTSCLQLSYIDDDGDKILINSDQELGILLQDKPQAARIRLYLHAVSVS
>Animalia_Chicken_tr|E1C0T1|E1C0T1_CHICK|5_96
DLSGKLIIKAQLGEDIRRIPIHNEDITYDELVLMMQRVFRGKLLSNDEVTIKYKDEDGDLITIFDSSD
LSFAIQCSRILKLTLFVNGQPRP
>Chromista_Hacrobia_tr|L1IDW1|L1IDW1_GUITC|279_345
RMVMEGLQSAFSEDLKGLPTSCLQLSYIDDDGDKILINSDQELGILLQDKPQAARIRLYLHAVSGS
>Animalia_ZebraFish_tr|Q6P0G5|Q6P0G5_DANRE|5_96
DLSGKLIIKAQLGDDIRRIPIHNEDITYDELLLMMQRVFRGQLQSSDEVTIKYKDEDDDLITIFDSSD
LSFAIQCSRILKLTLFVNGQPRP
>Animalia_Mouse_tr|Q9Z1A1|Q9Z1A1_MOUSE|5_96
DLSGKLIIKAQLGEDIRRIPIHNEDITYDELVLMMQRVFRGKLLSNDEVTIKYKDEDGDLITIFDSSD
LSFAIQCSRILKLTLFVNGQPRP
>Animalia_Human_sp|Q92734|TFG_HUMAN|8_96
GKLIIKAQLGEDIRRIPIHNEDITYDELVLMMQRVFRGKLLSNDEVTIKYKDEDGDLITIFDSSDLSF
AIQCSRILKLTLFVNGQPRP
>Animalia_Pig_tr|A0A287AB01|A0A287AB01_PIG|8_96
GKLIIKAQLGEDIRRIPIHNEDITYDELVLMMQRVFRGKLLSNDEVTIKYKDEDGDLITIFDSSDLSF
AIQCSRILKLTLFVNGQPRP
>Animalia_Dog_tr|E2REP2|E2REP2_CANLF|8_96
GKLIIKAQLGEDIRRIPIHNEDITYDELVLMMQRVFRGKLLSNDEVTIKYKDEDGDLITIFDSSDLSF
AIQCSRILKLTLFVNGQPRP
>Animalia_Pig_tr|A0A286ZXK0|A0A286ZXK0_PIG|43_129
DSGLVRTRFGAEFRRFSLERSKPGKFEEFYGLLQHVHKIPNVDVLVGYADIHGDLLPINNDDNYHKAV
STANPLLRIFIQKKEEAD
>Animalia_Frog_tr|A0A1L8HGI0|A0A1L8HGI0_XENLA|8_96
GKLIIKAQLGEDIRRIPIHNEDITYDELVLMMQRVFRGKLLTNDEVTIKYKDEDGDLITIFDSSDLSF
AIQCSRILKLTLFVNGQPRP
>Animalia_Frog_tr|A0A1L8H8F9|A0A1L8H8F9_XENLA|5_96
DLSGKLIIKAQLGEDIRRIPIHNEDITYDELVLMMQRVFRGKLLTNDEVTIKYKDEDGDLITIFDSSD
LSFAIQCSRILKLTLFVNGQPRP
>Animalia_Echinoderm_tr|W4XUZ7|W4XUZ7_STRPU|1_96
SMTVKAYLKRGENANAEIRRFVIDVAVSSNYEYLSKKVAQVFPSLGDPDYFSLSWKDSEGDNITFSSD
DELVEALGQINDDTFRIYVKEKKRCRR
>Animalia_Frog_tr|A0A1L8H1A4|A0A1L8H1A4_XENLA|13_114
CQALVIRIRIPDGGAVDWTVQPATQLLFRDILDVIGQVIPDATTTAFEYEDEDGDRITVRSDEEMKAM
```

LSYYCSMVLEQQGSGPLMEPLQIYPRACKPPGK
>Fungi_Mucoromycota|RhiirA1_1|412835|
fgenesh1_kg.155_#_28_#_step3_rep_c504|1_88
VNIKVIHKKAVRRFTLPSNATWIELEAKLRTLFNIPALSPFTLSYTDEDNDVITLSTDLELQEIFSSA
PTVKFDLKFSTSDDSSDSE
>Animalia_ZebraFish_tr|Q6TNS1|Q6TNS1_DANRE|13_103
LNMNAVEVKSKYGAEFRRFSVDRIKPGKFEEFYKLIMTIHRIANMEVMIGYADIHGDLLPINNDENFS
KAVSTAHPLLRIFIQRQEEVDY
>Animalia_Echinoderm_tr|W4YES8|W4YES8_STRPU|11_102
DMSNKLIIKAQLGDDIRRIPIHNEDITYDELVLMMQRVYRGKLNPSDEVVIKYKDEDGDLITIFDSTD
LSFAIQCSRILKITLFVNGQPRP
>Animalia_Human_sp|Q13501|SQSTM_HUMAN|36_105
AAAGPGPCERLLSRVAALFPALRPGGFQAHYRDEDGDLVAFSSDEELTMAMSYVKDDIFRIYIKEKKE
C
>Animalia_Mouse_sp|Q64337|SQSTM_MOUSE|1_107
ASFTVKAYLLGKEEATREIRRFSFCFSPEPEAEAQAAAGPGPCERLLSRVAVLFPTLRPGGFQAHYRD
EDGDLVAFSSDEELTMAMSYVKDDIFRIYIKEKKECRR
>Fungi_Mucoromycota|RhiirA1_1|499196|MIX11629_2584_100|1_83
ATIKVTYNSISRRFNIASPNWVELESKLRSLYNIPTTSSLIVSYNDEDGDVITLSSDLELKEILDQQS
SSRPIKLILSTVEN
>Animalia_ZebraFish_tr|F8W4T2|F8W4T2_DANRE|63_149
CSMLEVKSKYGAEFRRFSVDRYEPGRYKDFYRLIVRLHQLWHTDVFIGYADVHGELLPINNDDNFCKA
VSSTQSLLRIFIQLREEA
>Animalia_Dog_tr|J9P3W4|J9P3W4_CANLF|14_95
QVLVIRIKIPNSGAVDWTVHSGPQLLFRDVLDVIGQVLPEATTTAFEYEDEDGDRITVRSDEEMKAML
SYYYSTVMEQQVN
>Animalia_Human_sp|Q13163|MP2K5_HUMAN|14_95
QVLVIRIKIPNSGAVDWTVHSGPQLLFRDVLDVIGQVLPEATTTAFEYEDEDGDRITVRSDEEMKAML
SYYYSTVMEQQVN
>Animalia_Mouse_sp|Q9WVS7|MP2K5_MOUSE|14_95
QVLVIRIKIPNSGAVDWTVHSGPQLLFRDVLDVIGQVLPEATTTAFEYEDEDGDRITVRSDEEMKAML
SYYYSTVMEQQVN
>Animalia_ZebraFish_tr|A9JRD0|A9JRD0_DANRE|26_91
DMDSPVDSPSHLNFNDLLAAIRDAMPEATVTAFEYEDEVGDRITVRSDDELKAMLSYYCNTVMEQ
>Animalia_ZebraFish_tr|F1Q5Z8|F1Q5Z8_DANRE|1_95
SMTVKAYLIGKEDCNKEIRRFAVDQDVSTSFEYLQRKVLDVFVGLRTAPFQMYYKDEDGDMIAFSSDD
ELMMGLALVKDDTFRLFIKQRKEHKR
>Animalia_Chicken_tr|A0A1D5PLN4|A0A1D5PLN4_CHICK|1_98
AALTVKAYLLGKEDATREIRRFSLMPPVRYQAVHDRVAELFQGLLRAGPPPAFRMHYKDEDGDLIAFS
SDEELDLAMPYVQDGVFRVYIKEKKECRR
>Animalia_Frog_tr|Q6PGS1|Q6PGS1_XENLA|1_104
TVTVKAYLLGKDESHKEIRRFQLELPVAGKGKAASSGSSCEILANKVTDVFQGLKGGAFQMFYKDEEG
DLVAFSTDEELHMGLSLLNEDVFRIYIKEKKECKR
>Animalia_Pig_tr|A0A287A5U6|A0A287A5U6_PIG|13_114
NQVLVIRIKIPNSGAVDWTVHSGPQLLFRDVLDVIGQVLPEATTTAFEYEDEDGDRITVRSDEEMKAM
LSYYYSTIMEQQVNGQLIEPLQIFPRACKPPGE
>Protozoa_Choanozoa_Salpingoeca_tr|F2UE70|F2UE70_SALR5|46_110
DTAFLRYRIEESFEAALGGKEWQLKWKDEDDDMITIGDAEDLLVAVQSAQDDTLRLFVTTTTTT
>Animalia_Pig_tr|F1S445|F1S445_PIG|55_143
REIRRFSFCFSPEPEAEAEAAAGPGPCERLLSRVAALFPVLRPGGFQAHYRDEDGDLVAFSSDEELTM
AMSYVKDDIFRIYIKEKKEC
>Chromista_Stramenopile_Phytophthora_tr|D0MQN8|D0MQN8_PHYIT|29_96
DGRVASAGNELSFRDLRDYVLLVFPELKDVELLLYYIDDDSEQVRITNDAELDEAFRLMRELAVAAG
>Plantae_Rhodophyta_Chondrus_tr|R7Q2Z3|R7Q2Z3_CHOCR|1_86

DQSIILKVTFEGTIRRLPVTRDISFADFHQQLAERFSITVPFVTQYEDLDKDIITFDSDAELNDLFST
IEDTSKPLRISLLTLEE
>Fungi_Basidiomycota|Pucstr1|7191|evm.model.scaffold_16.223|2_89
SHQAPLSIKIKHADVIHKLQVNQHVPIWSTFLAAIAQRFGMPEEQPIGLQYLDPEGDTITISTQADFD
ELWHEVLSVTAQSSQTGGV
>Animalia_Nematode_tr|Q9U1W1|Q9U1W1_CAEEL|8_94
TSTILKARHADVVRKTSLHHANDLTLIDLVLNVQRLLALPSDANFVLKYKDEEGDLVTLAEDSDLLLA
LHTSGATLDVTVVVDSRA
>Plantae_Rhodophyta_Chondrus_tr|R7QME1|R7QME1_CHOCR|1_86
APLSVKACYKHTKRRFSLEPSSTFADFQAKLAAIFSIPAPRTILYKDDEDDLVAVSSDSELAELFAIA
TSASITPLRVYLYDTAE
>Plantae_Rhodophyta_Chondrus_tr|R7Q5J2|R7Q5J2_CHOCR|415_503
AMSFGVKASCTYGWQTQRRFVSVAYGLRFLSFKQVLCEIFDMDGVAFTVCYVDDDGDEIGISEDSDMR
PMFDLAKGKQGALRVRLRPP
>Animalia_Dog_tr|J9P3Q8|J9P3Q8_CANLF|178_268
SPDNIVEVKSKFDAEFRRFALPRASVSGFQEFSRLLRAVHQIPGLDVLLGYTDAHGDLLPLTNDDSLH
RALASGPPPLRLLVQKREADSS
>Plantae_Rhodophyta_Chondrus_tr|R7QC62|R7QC62_CHOCR|431_522
SGGSLIGVKAEFNGDIRRLRILPDASRASFMSQLQTLFEVPEILVLKYLDEEDDFVTVANETDMTEMI
YMVREHNLSPLRVHLQVERTECA
>Fungi_Basidiomycota|Agabi_varbur_1|112480|
estExt_fgenesh1_kg.C_40166|444_535
YDLTKIRVKIHYKDDVRGMALTPEMTYKEFMSKLASKFNKSVNGLGLKFQDEDGGKVTLADETDFELA
VETARTSTQGKAEGRLVIWCTDR
>Fungi_Ascomycota|Aspnid1|4239|AN6046|442_519
EVRRFRVKVHSFEDTRYILIPPTIEFAEFETRIREKFGFQMALKIKMQDEGDMITMVDQEDLDLLLMA
SREIARREG
>Fungi_Basidiomycota|Pucstr1|10491|evm.model.scaffold_2.653|443_536
SEMTKIRVKLHHGSDTRGMAVSIDMEFEDFLGKVIKKFSLRQGSVSMKYKDEEGSMVSIMDVDDWESA
IETARAYANGRAEGKVEIWVEDGYL
>Fungi_Basidiomycota|Pucstr1|21991|evm.model.scaffold_50.27|443_536
SEMTKIRVKLHHGSDTRGMAVSIDMEFEDFLGKVIKKFSLRQGSVSMKYKDEEGSMVSIMDVDDWESA
IETARAYANGRAEGKVEIWVEDGYL
>Fungi_Ascomycota|Schpo1|1006|scd2|454_536
TAGSTCKVKVRLGDETFALRVPSDISFEDFCERLTNKLGECEHLSYRDTNANKVLPLNNVDDLRKACS
QESGVLLFAERRRF
>Fungi_Mucoromycota|Morel2|235636|Morel1.CE75622_11690|1_88
ADSNQRPCKVSYNGSLRRFLIARPAIWQDFENKIRNVYSIPGNLALDVQYKDDEGDLITLNTDSELDD
VLAMHTIFTPLAPVKFDVV
>Chromista_Stramenopile_Phytophthora_tr|D0MYM0|D0MYM0_PHYIT|435_513
ADCYGNNHRFTSSAESVKQLLVDVQNRLGDNTIRRILYVDDEGDHVLLSEDSDLKDAVNRARTWGNKY
IRLIVPHYRL
>Fungi_Ascomycota|Sacce1|462|YBR200W|473_551
SGLKTTKIKFYYKDDIFALMLKGDTTYKELRSKIAPRIDTDNFKLQTKLFDGSGEEIKTDSQVSNIIQ
AKLKISVHDI
>Animalia_Frog_tr|A0A1L8ESG6|A0A1L8ESG6_XENLA|39_128
NKQADVRIKFEHSGERRILQFSRPVKYEEVEQKVKNVFGQQLDLYYMNNELSIPLRSQDDLDKAVDLL
DRSSSMKSLRILLLSHDRNHI
>Animalia_Chicken_tr|F1NUM0|F1NUM0_CHICK|103_192
HQVLVIRIKIPDGGAVDWTVHSAPQLLFRDVLDVIGQVLPDATTTAFEYEDEDGDRITVRSDEEMKAM
LSYYYSTVMEQQVNGQLIEPL
>Fungi_Mucoromycota|Morel2|140392|
Morel1.fgenesh1_kg.57_#_16_#_Locus3168v1rpkm66.61|495_576
GLRDKLRVKCHYIDTRAVLVRVDTSLQELLQKVQEKFQADRPLKLKYKDEDHHMLSMIDDEDWLMAQQ

VQMETQGTLDRME
>Animalia_Frog_tr|A0A1L8GAK2|A0A1L8GAK2_XENLA|13_104
DHSHQVRVKAYFKGDIMITHFEPSITFDGLCNEVRDMCSFENDQPFTMKWIDEEGDPCTVSSQLELEE
AFRLYELNKDSELLIHVFPCIPE
>Animalia_ZebraFish_sp|Q90XF2|KPCI_DANRE|13_106
ENPHQVRVKAYYRGDIMITHFEPSISYEGLCNEVRDMCSMDNDQLFTMKWIDEEGDPCTVSSQLELEE
ALRLYELNKDSELIIHVFPCVPEKP
>Animalia_Frog_tr|A0A1L8FG56|A0A1L8FG56_XENLA|9_102
DLSKDIKIKAHYKGDILITRLGALMNYEQLCEEVREMCYLNQRQPITLKWIDDEGDPCTISSQMELEE
AFRLYSIYKEEGLSIHVFPSIPEKP
>Animalia_Frog_tr|A0A1L8FM28|A0A1L8FM28_XENLA|10_101
LSKDIKIKAHYKGDILITRLEAGMNYEQLCEEVREMCYLNQRQPITLKWIDDEGDPCTISSQMELEEA
FRLYSIYKEEGLSIHVFPSIPEK
>Animalia_Dog_tr|F6XMN6|F6XMN6_CANLF|11_102
SGCRVRLKAHHSGDILITSLDAAMTFEELCDEVREMCSLRQGHPLTLKWVDNEGDPCTVSSQMELEEA
FRLACQRRDEGLIIHVFPSTPEQ
>Protozoa_Amoebozoa_Dictyostelium_tr|Q54EB5|Q54EB5_DICDI|1_85
GITYKSNFEGDVRRFSSDHPLTYTRLQDKLVNLYNLYEISFGITYLDDGDNITIADAKDLEEAHNLLG
NEILRLTITRKLNENK
>Animalia_Mouse_sp|Q02956|KPCZ_MOUSE|10_103
RSGGRVRLKAHYGGDILITSVDAMTTFKDLCEEVRDMCGLHQQHPLTLKWVDSEGDPCTVSSQMELEE
AFRLVCQGRDEVLIIHVFPSIPEQP
>Animalia_Chicken_tr|E1BQN6|E1BQN6_CHICK|11_102
SKDIIRVKAHYSGDILITNLDAYISYDELCDEVREMCNLQQEQPITLKWIDDEGDPCTISSQMELEEA
FRLYCQNREEGLIIHVFPSIPEK
>Animalia_Human_sp|Q05513|KPCZ_HUMAN|11_102
SGGRVRLKAHYGGDIFITSVDAATTFEELCEEVRDMCRLHQQHPLTLKWVDSEGDPCTVSSQMELEEA
FRLARQCRDEGLIIHVFPSTPEQ
>Animalia_Pig_tr|A0A286ZSY3|A0A286ZSY3_PIG|11_102
NGDRVRLKAHYSGDILITSLDSATTFEELCEEVREMCCLSRDHPLTLKWVDSEGDPCTVSSQMELEEA
FRLSSQRRDEGLIIHVFPSIPEE
>Plantae_Chlorophyta_Ostreococcus_tr|A0A096P8H7|A0A096P8H7_OSTTA|
193_260
VTCEGEVKSIKLPVQLRYSDLVNAIKSEFGIEKYVAVKYRDFDGDFVTITSRMDLRTALTNFAAVAE
>Animalia_Mouse_sp|Q62074|KPCI_MOUSE|20_113
DHSHQVRVKAYYRGDIMITHFEPSISFEGLCSEVRDMCSFDNEQPFTMKWIDEEGDPCTVSSQLELEE
AFRLYELNKDSELLIHVFPCVPERP
>Animalia_Human_sp|P41743|KPCI_HUMAN|20_113
DHSHQVRVKAYYRGDIMITHFEPSISFEGLCNEVRDMCSFDNEQLFTMKWIDEEGDPCTVSSQLELEE
AFRLYELNKDSELLIHVFPCVPERP
>Animalia_Nematode_sp|Q19266|KPC3_CAEEL|8_99
EDGDIKLKTRFQGQVVVLYARPPLILDDFFALLKDACKQHKKQDITVKWIDEDGDPISIDSQMELDEA
VRCLNSSQEAELNIHVFVGKPEL
>Fungi_Zoopagomycota|Coere1|81886|fgenesh1_kg.21_#_28_#_isotig03066|
62_154
QPVVPFRIKFVNNKEEATMADIYPYEEFNDIVQRITAKLRMTRHAEYVLMYKDNDDEEIGVACSDNLR
EMFAIFEPGSRLQLRIVPFNINNS
>Animalia_ZebraFish_tr|B0S4T7|B0S4T7_DANRE|11_102
PSEYVKIKAHYGGDMLISDLDLALTYTEVCKEVREMCGVRKETPITLKWIDDEGDPCTISSQMELEEA
FRIYSRNRHSGLLLHVFPSIPEK
>Animalia_Chicken_tr|E1BWA3|E1BWA3_CHICK|23_114
DSAHQVRVKAYYKGDIMITYFEPSISFEGLCSEVRDMCSFDNEQLFTMKWIDEEGDPCTVSSQLELEE
AFRLYELNKDSELLIHVFPCVPE
>Fungi_Mucoromycota|Morel2|133812|

```
Morel1.fgenesh1_kg.8_#_170_#_Locus2404v1rpkm90.59|459_549
GNEFTFKFQESSGQAHRFSSSTRSLAELRNKIIEKIGGLEAGTDVVLSYCDEEGDNVMILQDSDIVDA
VQMAYRSQSSLVRLSVTISSKE
>Plantae_Chlorophyta_Ostreococcus_tr|A0A090M2T8|A0A090M2T8_OSTTA|
193_263
PPALQIKAVFGQDVRMFSVFSTIGFKDLVTSIATKFNFAGQFSIKYEDEEGVMRNVQSKSDFQKSIYA
TS
>Fungi_Chytridiomycota|Rhihy1|712469|
fgenesh1_kg.5_#_132_#_Locus3843v1rpkm60.52|463_539
GKLVRFTLMTPTTLADLRAIASGRLGVAPTSQDLRLSYEDDDGDWVLVSSDADLDDAISMARRLGEKL
SLRVGDDV
>Protozoa_Amoebozoa_Dictyostelium_sp|Q867T7|NCFA_DICDI|304_389
IQDVKITLKVFYKDRRLIQIPVPCNLSTFIQKIELKFEITISDKFSLSFQLDGEENEINSQVQLDKMI
CMEINEINVKDIIPSPS
>Fungi_Ascomycota|Canalb1|60187|CAALFM_C112250CAT0|440_544
MTVDEIPFIFKFKSPGIEGRVHRITLKASDGIIKLRELMNEKLHDKDFAFLNVPKPDGNSSTQETYAI
SYVDDEGDVVSITSDSDLAECIRINLNLQNEKADLY
>Animalia_FruitFly_sp|A1Z9X0|APKC_DROME|25_118
NTPNSITVKTAYNGQIIITTINKNISYEELCYEIRNICRFPLDQPFTIKWVDEENDPCTISTKMELDE
AIRLYEMNFDSQLVIHVFPNVPQAP
>Animalia_Frog_tr|A0A1L8ENK5|A0A1L8ENK5_XENLA|27_113
KQNAVRVKFEHRGEKRILQFSRPVLLEDLYAKAKVAFGQCMDLHYSNNELVIPLKTQDDLDKAVELLD
RSVHLKSLKILLVLHGHT
>Fungi_Chytridiomycota|Rhihy1|725792|
fgenesh1_kg.106_#_37_#_Locus1385v1rpkm158.10|457_548
LDPSQFGYKLRDKRSGKVHRFTSSSMNVSDVYAVIRAKTGGAPGVVSYEDEEGDMILLGSNADLEEAV
GMARRFGWERLVLHVGEQPKEKE
>Fungi_Ascomycota|Aspnid1|3158|AN7030|534_616
SSGGALKVKVNFQDDLIAIRVPSDINVQQLKEKLMDRLKINDEIVVQYKDEASGAYVDLISDSDLDTA
IQRNSKLTLYVGLA
>Fungi_Mucoromycota|RhiirA1_1|367976|estExt_Genewise1.C_8910004|
462_556
RGHDENFFTFKFKAPNGKSHRFTVDYTSFEHIRATVASKLPSNVRDFTIYYVDEDDDHVSMSNDDDVI
DAVRIAQRQGMSRVILHIQEIEKKSL
>Animalia_Frog_tr|A0A1L8FQI5|A0A1L8FQI5_XENLA|33_118
QQDARIKFEYNGEKRIIEFRRPIKLKDVQQKITDAFGQTMDLHYANNELLIPLMCQEDMDRALEYLDT
CPALKSLRILVKSPKKM
>Animalia_Human_sp|Q9Y2U5|M3K2_HUMAN|38_127
KKQNDVRVKFEHRGEKRILQFPRPVKLEDLRSKAKIAFGQSMDLHYTNNELVIPLTTQDDLDKAVELL
DRSIHMKSLKILLVINGSTQA
>Animalia_Mouse_sp|Q61083|M3K2_MOUSE|39_126
KQNDVRVKFEHRGEKRILQVTRPVKLEDLRSKSKIAFGQSMDLHYTNNELVIPLTTQDDLDKAVELLD
RSIHMKSLKILLVVNGSTQ
>Animalia_Dog_tr|E2RN16|E2RN16_CANLF|40_126
KQNDVRVKFEHRGEKRILQFSRPVKLEDLRSKAKIAFGQSMDLHYTNNELVIPLTTQDDLDKAVELLD
RSIHMKSLKILLVINGST
>Animalia_Chicken_tr|Q5ZLW4|Q5ZLW4_CHICK|36_125
KQQRNVRIKFEYEGEKRIIQFPRPVKFKEVVQKVTDAFGQTMDLVCMSNELLIPLKSQEDLDKAMEQL
ELSPSLKSLRILVSAPKKANL
>Animalia_Pig_tr|I3LNQ4|I3LNQ4_PIG|40_126
KQNDVRVKFEHRGEKRILQFPRPVKLEDLRSKAKIAFGQSMDLHYTNNELVIPLTTQDDLDKAVELLD
RSIHMKSLKILLVMNGST
>Animalia_ZebraFish_tr|E7F683|E7F683_DANRE|38_123
NQNDVRVKFEYRGEKRILQFPRPVSLEDLSAKAKVAFGQSMDLHYTNNELVIPLSTQDDLDKAVELLD
```

RSVHMKSLKILLVLPSW
>Animalia_ZebraFish_tr|A0A0R4ITX2|A0A0R4ITX2_DANRE|57_141
YRQDLRVKLEHEREKRIIPFQRPLKFKDLLQKVTEAFGQQMDLYFTEKEMLVALKCQEDLDRAIQGLS
SSSGMNNLLRVILKTP
>Animalia_Frog_tr|A0A1L8EL30|A0A1L8EL30_XENLA|39_128
NKQADVRIKFEHSGERRILQFSRPVKYEEVEQKVKTVFGQPLDLYYMNNELSIPLRSQDDLDKAVDLL
DRSSSMKSLRISLFSHDRNHI
>Animalia_Dog_tr|F1P750|F1P750_CANLF|39_128
NRQSDVRIKFEHNGERRIIAFSRPVRYEDVEHKVTTVFGQPLDLHYMNNELSILLKNQDDLDKAIDIL
DRSSSMKSLRILLLSQDRNHT
>Animalia_Human_sp|Q99759|M3K3_HUMAN|39_128
NRQSDVRIKFEHNGERRIIAFSRPVKYEDVEHKVTTVFGQPLDLHYMNNELSILLKNQDDLDKAIDIL
DRSSSMKSLRILLLSQDRNHN
>Animalia_Mouse_sp|Q61084|M3K3_MOUSE|39_128
NRQSDVRIKFEHNGERRIIAFSRPVRYEDVEHKVTTVFGQPLDLHYMNNELSILLKNQDDLDKAIDIL
DRSSSMKSLRILLLSQDRNHT
>Animalia_Frog_tr|A0A1L8EVE8|A0A1L8EVE8_XENLA|45_134
KKQNDVRVKFEHRGEKRILQFSRPVLLEDLYAKAKVAFGQCMDLHYSNNELVIPLKTQDDLDKAVELL
DRSVHMKSLKILLVLHGHTEV
>Animalia_ZebraFish_tr|E7EXX1|E7EXX1_DANRE|49_133
LQEDVRIKFEFCGERRILMFGRPVQFEEVQQKVKTFFGQQLDLHYMNNELSIPLRDQDDLDKAIDLLD
RSSNMKSIKIMLLTQE
>Plantae_Chlorophyta_Ostreococcus_tr|A0A090M3U8|A0A090M3U8_OSTTA|
496_585
DDTEKIQVKLFLEEDIRFLEIEPDVSFEELVIAVGKVFIESYSIKFEDADKHHITLRSTDDVRIACRQ
HERTAATYLKLLLDKASKEKS
>Chromista_Hacrobia_tr|L1JV30|L1JV30_GUITC|529_630
CKKSVLPLKATFSSQDADSVIRRLSVPVELDEEEVPVEGFSAVTDALVYAFRNELKMGCDDVKISYTD
EDGDEVMISCDEELGLAMQQFRDASVMRVKLHA
>Fungi_Chytridiomycota|Rhihy1|852374|estExt_Genemark1.C_320167|
552_623
TITIRSVVKGVDLFRLDVPRDVRYDELVDGVERILGKRVGGLEYLDESGMMVTLHGDEDLGLMVRTWG
VLE
>Animalia_Pig_tr|F1SH28|F1SH28_PIG|60_151
DHSHQVRVKAYYRGDIMITHFEPSISFEGLCNEVRDMCSFDNEQLFTMKWIDEEGDPCTVSSQLELEE
AFRLYELNKDSELLIHVFPCVPE
>Protozoa_Choanozoa_Salpingoeca_tr|F2TVT2|F2TVT2_SALR5|2_89
VQPGIRFKCCLGQENRNIFLDSPVSLDQLKQKIVAYFRKPNLNILVVDTEGKRKRLRTQDDLDQCLKS
VDAQSSSTAVRLVLTSSTD
>Protozoa_Amoebozoa_Dictyostelium_tr|Q55CE3|Q55CE3_DICDI|1_87
VNLILKIQHNDDTRRVSMERDPTFLELRKMTVTFFKINSFLIKYFDEDKDLITITSDNDLKEAFSIAT
TSPRTVRLFVSKTEEESS
>Plantae_Chlorophyta_Chlorella_tr|A0A2P6U062|A0A2P6U062_CHLSO|
159_253
TSGRQVVFPAKLSMGDDTRFLQLVPGVTYLELMEHVRQLYPAAGPFVLKFVDKEGDLVTIAGARDIQR
AMQEAVETAQRGSRNVQLTQQSLPPI
>Fungi_Mucoromycota|RhiirA1_1|455712|gm1.8193_g|559_646
QKPTKIKIKCYHKDTRVVLVSANIRYPDLIKRIQEKFAIGASLQLKYKDEDGTMVTMIDQDDLEMAIS
MVPDAATDTGRMEIWIEME
>Fungi_Basidiomycota|Agabi_varbur_1|81204|
fgenesh1_kg.1_#_90_#_1866_1_CFAF_CFAG_CFAH_EXTA|471_568
PSGRTHRFQSRHDDVAHLREIVAGKLATDPFFTELQLEQDGGPPADPNDFHLSYTDADGDTVLITSDD
DVTDAVKIARTAGLDRVVLYVQGGKGWKD
>Fungi_Basidiomycota|Agabi_varbur_1|110685|

```
estExt_fgenesh1_kg.C_10692|1_87
PTHFKLKFDSLTRRATFAHHPSWGQLSAKVASLFSIPQHQVAVTYVDADDEEITLSTEEELQDYYQSS
LPGDSFKLSVRDLSRRNQ
>Animalia_Pig_tr|A0A287AGW3|A0A287AGW3_PIG|70_159
KFLSDVRVKFEHNGERRIIAFSRPVRYEDVEHKVTTVFGQPLDLHYMNNELSILLKNQDDLDKAIDIL
DRSSSMKSLRILLLSQDRNHT
>Fungi_Mucoromycota|Morel2|35585|Morel1.estExt_Genemark1.C_760063|
586_662
ADEFIKLKVSYLDDISAMRIPVSVTFQSLQQKIFERLDCDPKPLSYRDSRGDFAALQTDIEVREAIDH
CGGKLVIY
>Fungi_Ascomycota|Aspnid1|10154|AN5716|479_584
SPFPFKFKAPSGRVHRVNILPAAGIAELVAQVTAKLGPEVEAVGGAASCADGVLSNTGYALSYVDNEG
DTVSITTDQDLVDAVYIARHARRDKVDLFVHDPAQPP
>Fungi_Mucoromycota|Morel2|131755|
Morel1.fgenesh1_kg.1_#_160_#_Locus6161v1rpkm26.08|578_666
GLRNKLRVKCHYIDTRAVLVRADTPLEELIQRIQEKFQADRPLKLKYKDEDQHMLSMVDDEDWLMAQQ
VHMETTGSLDRMELWCFDEE
>Animalia_Echinoderm_tr|W4Y4I9|W4Y4I9_STRPU|66_159
SHSTEIRVKFEFNGEKRILQIPRPLKYDDILLKARTTFGQPVDMFLTNHNTTFSLLIPIRDQTDLNHA
VEITDNNPNAKSLRLHLELATPGSS
>Fungi_Zoopagomycota|Coere1|80618|fgenesh1_kg.10_#_71_#_isotig02782|
590_674
KKDSMKVKVHFGDDIINLMVPKLAAFETLRAKISAKIANAVASGQPSRAALRIQYLDEDGEAVLMTDE
DDFELAKAYAGGDMSA
>Plantae_Chlorophyta_Chlamydomonas_tr|A0A2K3D9U2|A0A2K3D9U2_CHLRE|
191_280
LQQGVFVAKATLGDDTKLVHLSLSNSYADVLAAVQQKFPSAGAFLLKYVDKNGDLITLTCKADMHTAL
GELVQQYQRQVQGQGAHGPKL
>Animalia_Dog_tr|F1PG28|F1PG28_CANLF|127_218
DHSHQVRVKAYYRGDIMITHFEPSISFEGLCNEVRDMCSFDNEQLFTMKWIDEEGDPCTVSSQLELEE
AFRLYELNKDSELLIHVFPCVPE
>Animalia_Chicken_tr|A0A3Q2TZP2|A0A3Q2TZP2_CHICK|69_158
KIPNDVRIKFEYHGERRIIPFVRPVRYEDVQQKVKTAFGQPLDLRYVNNELSIPLKNQDDLDKAVDLL
DRSSNMKSLRILLLSQDRNHL
>Fungi_Mucoromycota|Morel2|124649|Morel1.fgenesh1_pg.22_#_71|639_718
SSEEMIKIKISYHEDIMAMRIPVSISFRSLQQKIFERLHSDHKDLSYRDDRGDFATIQNDGDVRDAID
RSGGKLMIYVD
>Fungi_Basidiomycota|Pucstr1|16142|evm.model.scaffold_31.154|2_106
SGQSPLFIKIKYAGSTRKVRVPNQPAPVWSKLSKAIIDRFGIPENQPIGLQYVDSDGDAITISSQVEF
DELWHEITLAARPSQKNGANQRALSLELVLIDPITP
>Fungi_Basidiomycota|Pucstr1|3214|evm.model.scaffold_12.295|2_106
SGQSPLFIKIKYAGSTRKVRVPNQPAPVWSKLSKAIIDRFVIPENQSIGLQYVDSDGDAITISSQVEF
DELWHEITLAARPSQKNGTDQRALSLELVLIDPINP
>Fungi_Ascomycota|Schpo1|1292|mug70|567_654
NPQSPSQFTIKYRSIAGRVHRLRLDGINSVSDLRTAVEEREKEQLVTLTYIDDEGDVVELVSDSDLRE
AILLARRRGLPRLEVRGVA
>Fungi_Mucoromycota|Morel2|1817815|fgenesh1_pg.19_#_89|653_742
NGVPQLKVKVNYQEDSYLIVVPVQIGYSELIERVERKIRLCGSRRTESQPLRLRYKDEDNDYINMEDD
EDISLAMESCFADGVMNLHVI
>Plantae_Rhodophyta_Chondrus_tr|R7QPC9|R7QPC9_CHOCR|372_470
GDVALASFKFKDLNHEYRRIKMPMPPAPGAFDQFVVDIRRRFAGSANVGPIKIKYVDEDGDEVLLSND
EDLASCYEDFLESKNRTIHLRVYDTERPTS
>Protozoa_Amoebozoa_Dictyostelium_tr|Q86KD2|Q86KD2_DICDI|1_88
NILIKSELENDKRRFRLKECSFSCLCYTLASIYSFYNDMIYSIFYLDNENEWITLASTDDLKESYSLC
```

PSLIRIKIIVLDLTINNNN
>Fungi_Zoopagomycota|Coere1|81531|fgenesh1_kg.17_#_68_#_isotig02469|
726_836
GQSKAVKVKVHFQNDIFVVIIRNNVTFDDLVSRVDRKIKICAGPHVGISSSNEHDIASAPVLNIRMRY
QDEDGDMILIGSDEDVQLAFESASAAKKDDTSMSTLNLFVSI
>Fungi_Ascomycota|Canalb1|56321|CAALFM_C501970CAT0|744_831
PTFDVAIKLLYKSTELSEPLIVNAQIEYNDLLQKIISQIITSNLVADDVNISRLRYKDDEGDFVNLNS
DDDWGLVLDMLTSEDFYQT
>Fungi_Ascomycota|Sacce1|36|YAL041W|781_845
TLLVEKVWNFDDLIMAINSKISNTHNNNISPITKIKYQDEDGDFVVLGSDEDWNVAKEMLAENN
>Protozoa_Choanozoa_Salpingoeca_tr|F2UE56|F2UE56_SALR5|1_86
AQTVEWVKVSLGDDLRRMRVSQTFEQLRSRVQEAFHLAGTAFTLKWVDEDGDLITLADDTDLIAMLAH
PKACVRAYVQLDGSGND
>Fungi_Chytridiomycota|Rhihy1|720875|
fgenesh1_kg.45_#_45_#_Locus2061v1rpkm110.78|13_89
WEDIVRVKVEHVGTVRQIVFDEAEVPWTLFASKVKEVFRIPTATSISAFYIDEDGDSIQIDSDVEIQY
LLKLPRVP
>Fungi_Ascomycota|Schpo1|1489|scd1|770_872
NTTNVKIRLRLHEVSLVLVVAHDITFDELLAKVEHKIKLCGILKQAVPFRVRLKYVDEDGDFITITSD
EDVLMAFETCTFELMDPVHNKGMDTVSLHVVVYF
>Fungi_Mucoromycota|Morel2|124606|Morel1.fgenesh1_pg.22_#_28|800_889
AGVPQLKIKVNFQEDAYLIVVPIQIGYNELIERVEKKIRLCGCRRSDSQPLRLRYKDEDNDYIIIKDN
DDILLAFESCYAEGVMNLHVS
>Fungi_Mucoromycota|RhiirA1_1|409512|
fgenesh1_kg.59_#_31_#_ACTTGA_L001_R1_(paired)_contig_7603|803_894
NFSTTMKIKVNYAEDIFVIVVPQNIEYKELCDRVERKIRLCTTQRDESIPLRIKYQDEDGDHITINSD
EDVLMAFEGRLAAGGNFVNLYVS
>Plantae_Chlorophyta_Ostreococcus_tr|Q015Z7|Q015Z7_OSTTA|816_896
VVANAISVKASVEDDVVRFKLTSDMTFAALLAKLHLNGKKEVTLHYLDDGDEWIRLGGDLDLLEARIC
GDAAGALRIKCM
>Fungi_Mucoromycota|RhiirA1_1|429261|fgenesh1_kg.2247_#_2_#_TR18143|
c0_g1_i1|5_88
INFKVSCTSDPTVLRKLNFKNSDLTYSSLYAKLSALFKLDNFIIRYTDEDEDAICIDNDKELKDAIDH
ALNLGKNNARVVIRL
>Fungi_Ascomycota|Aspnid1|10298|AN5592|810_913
FMPTQLKAKVNFDENYVTLVISSNIGFRTLTDRVDAKLARFTNRSIGSKTVRLRYQDEDGDFVTIDSD
EAVQLAFVEWKEQHREELARGQVGEIQLFCQPIEN
>Animalia_Chicken_tr|E1C6F8|E1C6F8_CHICK|1_87
EPQVNLRVSYRGETQSFLVSDTAHTTWADVEAMVKVSFDLDDIQIKYIDEDNDEVSVNSKEEYEEALK
IAVKQGNQLQMNVYEESS
>Protozoa_Amoebozoa_Dictyostelium_sp|Q54R82|MKKA_DICDI|10_101
FVDPFIRIKCILGDDIRIIKFNSNISYGGLMNQLEQDFQCPISIHQYEDYEGDKVTVKSKDDIMEALT
MYFELKALNPTKIISTKFFLKQL
>Protozoa_Choanozoa_Monosiga_tr|A9UPL7|A9UPL7_MONBE|6_80
GDESVVVKFRWQDSVRRLHFERLPDIQELVAQLEAVTKASADQITYVDSDGDRVTIDTTIELREARRE
LGQKLV
>Animalia_Dog_tr|E2RL96|E2RL96_CANLF|1_90
EPQVTLNVTFKNETQSFLVSDPENTTWADVEAMVKVSFDLTTIQIKYLDEENEEVSINSQGEYEEALK
MAVKQGNQLQMQVHEGSRVDE
>Fungi_Basidiomycota|Agabi_varbur_1|54553|estExt_Genewise1.C_30413|
3_114
CPDKPLLVRCAFNGRSKRITFQSARNCNYDLLRRKVDQCFSLYGTPYLIAWKDDDGETTNITTDNDLV
ETIKYFHDGGEAPLSSAASILSGRSFSSRKITIHVDVIMEYDG
>Animalia_Frog_tr|A0A1L8EMC4|A0A1L8EMC4_XENLA|1_90

ESQVNLFVSCNGESQNFLVSNSENTTWADVETMVLVSYDLNYIQIKYIDEDNEEVSVNSQGEYEEALK
SAVKQGGLLRINVYDKQQPSK
>Animalia_Human_sp|Q14596|NBR1_HUMAN|1_90
EPQVTLNVTFKNEIQSFLVSDPENTTWADIEAMVKVSFDLNTIQIKYLDEENEEVSINSQGEYEEALK
MAVKQGNQLQMQVHEGHHVVD
>Animalia_Frog_tr|A0A1L8ETV5|A0A1L8ETV5_XENLA|1_90
ESQVNLCVSCNGESQNFLVSDSENTTWADVETMVLVSYDLNDIQIKYMDEDNEEVSVNSQGEYEEALK
SAVKQGGLLRMNVYEKQQPSK
>Animalia_Mouse_sp|P97432|NBR1_MOUSE|1_91
EPQVTLNVTFKNETQSFLVSDPENTTWADVEAMVKVSFDLNTIQIKYLDEENEEISINSQGEYEEALK
MANIKQGNQLQMQVHEGYHVVD
>Fungi_Basidiomycota|Agabi_varbur_1|65097|estExt_Genewise1.C_390041|
900_988
DQSPPVKVKVHFHEDIFVIQVPRVTEFDDLVEKVGKKIRLCGPRRDDGPLKVKYRDEDGDLVSLGSTE
DVQIAFESFRPGGQVTLFVT
>Animalia_ZebraFish_tr|E9QEG9|E9QEG9_DANRE|1_90
NLPVTVKVNFRGNVKKFPVLDTNKAQWETVEAWIKTTFGLSHFQVKYFDEDNEEVCINSQDEYTEALK
SAFKQANQLHMNVYKMKGQAE
>Animalia_Pig_tr|A0A287A9L9|A0A287A9L9_PIG|1_90
EPQVTLNVTFKNETQSFLVSDPENTTWADVEAMVKVSFDLNTIQIKYLDEENEEVSINSQGEYEEALK
MAVKQGNQLQMQVHEGYSVVD
>Animalia_ZebraFish_tr|F1R2I7|F1R2I7_DANRE|1_90
DFYINLKVTFRGNAKSFLLSGSETKSWESMEAMVKRSFGLCNLQLTYFDEENEEVSVNSQLEYEEALK
SAARQGNRLQMNVYETRGSRA
>Plantae_Rhodophyta_Chondrus_tr|R7Q3Y4|R7Q3Y4_CHOCR|915_1011
EKKKTFIVKASCRYGWANSTRRLSVSMKTAFPAFKTRLGDAFDMIKPFTITYRDEEGDFVKVSSEAEM
AHLLNMAGRHVRPVQVKLIPPHGLFKTS
>Protozoa_Amoebozoa_Dictyostelium_tr|Q54Q21|Q54Q21_DICDI|48_141
NNLDSTTLNVTLGSEMKSITVPKSSTYKDMISTIKDKFGVNSKSTLCIKCENKDGEMFSLASDCHVKK
AYNQQPENQPKELRLVVKEIPQKKC
>Protozoa_Euglenozoa_Trypanosoma_tr|Q38CG8|Q38CG8_TRYB2|606_710
SDVDGFKLKIHFEAQLRVLRIQGTETCTFDEVYKRVHDLCTQQTALHHRPAGQKLRLRYEDAEGDCIS
LLTQEDWNVFISEQAPSGLRGAKLEIYCDFPPVPSQ
>Fungi_Basidiomycota|Ustma2_2|9640|mRNA_UMAG_02422|1014_1102
SSAIKLNISFAEDRYVVVVLSSTPFSTLLEKVTKKIRLCSGKNLEHTLRMRYIDEDGDAVLISDDDDV
QMAFDSARASPAGEVELVVN
>Animalia_Echinoderm_tr|W4ZC17|W4ZC17_STRPU|1_88
EENIALEVRFEGETQIYSPAYKTKWQDLLAMLKCSFDLEDIVVTYIDDEEDEIAVDTEEEYDQAKALA
SKCNNVLRLRVSRVIILDT
>Fungi_Basidiomycota|Pucstr1|23587|evm.model.scaffold_58.102|
1085_1192
NGPTSIRFKLKAGEDTYVIVTLSTVTYQELIAKILKKLKNCGVSSSKDPAHSISSSSSGSAGYAGKIK
IRYEDEEGDLILICNDEDVGMAIDWMKSVGISHLMLFVD
>Fungi_Basidiomycota|Pucstr1|6457|evm.model.scaffold_150.47|
1085_1192
NGPTSIRFKLKAGEDTYVIVTLSTVTYQELIAKILKKLKNCGVASSKDPAHSISSSSGGSAGYAGKIK
IRYEDEEGDLILICNDEDVGMAIDWMKSVGISHLMLFVD
>Plantae_Chlorophyta_Chlamydomonas_tr|A0A2K3D924|A0A2K3D924_CHLRE|
17_132
KKDDRIRVRLHYGGKFAQDAPNLWRYVGGEVFNESFPLEAKYADVCLRLNDKFGDTVSFKYLCPGDDL
DPDNLVQVQGDDDLTEMKDEYAHAVGDTRSRTVRLKIYVFRAVIFER
>Protozoa_Choanozoa_Salpingoeca_tr|F2U486|F2U486_SALR5|681_765
LHRLPHPFTAHHISVIMEVAAHARWEEVQSDLQRTLGFALSRVQYVDDEDDWITISSDEDWTSAVTLY
PAKLFLQVDKASACVS

```
>Plantae_Chlorophyta_Chlorella_tr|A0A2P6TY08|A0A2P6TY08_CHLSO|
810_911
RFAQGEDGAWRYLGGDHFLESVPESTKYADLMFSLVEKVDGAVSVKYQMPGEELDPEALISVNDDGDI
KELFAEYQHALRLPGTPIKTFRLRLFLFPAAEE
>Protozoa_Euglenozoa_Leishmania_tr|Q4QC60|Q4QC60_LEIMA|1204_1298
RLLDTLRLKIHFDAQTRVLPVADAAQASFHDVYKQIHDFCQTQLQQPPRAAGAERRLRIRYEDTEGDF
ISLLDQQDWHMMLSELAPHGCGGVKI
>Plantae_Chlorophyta_Chlamydomonas_tr|A0A2K3D147|A0A2K3D147_CHLRE|
1914_2016
FSACRATGHWEYGGGETRLVAVEEAAGFERFRDSVFAACKGRVRSPDEASLRYELPSCPGTLVDVRSD
GDVGMMWEELADFAESSGKPSYKLHVYVCHHLLL
>Chromista_Stramenopile_Phytophthora_tr|D0N4K8|D0N4K8_PHYIT|
3091_3187
KNMDDNTFVFKVSDGAQGHFHRIMCRFNSMGPLLEQIRFKMGMDDNEALRLKYEDDEGDLALLTSDES
LVEAVHMAQRAGWKRLVLVVDVVKRPQH
```
