## Additional File 6 for "Deep Evolutionary History of the Phox and Bem1 (PB1) Domain Across Eukaryotes"

```
# Load required libraries
library("protr")
library("Peptides")

# Load FASTA files from protr
landplant_raw <- readFASTA("PlazaOrthologues_selectedPB1.fa")
length(landplant_raw)

# Load class/clade information
landplant_metadata <- read.table("PlazaOrthologues_metadata.txt",
sep = "\t", row.names = 1,
                                 col.names = c(NA, "Species",
"GeneFamily", "OrthoGroupID"))

# Amino acid type sanity check and remove the non-standard sequences
from protr
landplant <- landplant_raw[(sapply(landplant_raw, protcheck))]
length(landplant)

# Sequence length from Peptides
landplant_length <- data.frame(Length=(sapply(landplant,
lengthpep)))

# Molecular weight from Peptides
landplant_mw <- data.frame(MolWt=(sapply(landplant, mw)))

# Net charge from Peptides
landplant_charge <- data.frame(NetCharge=(sapply(landplant, charge,
pH = 7, pKscale = "EMBOSS")))

# Isoelectric point (pI) from Peptides
landplant_pI <- data.frame(pI=(sapply(landplant, pI, pKscale =
"EMBOSS")))

# Aliphatic index from Peptides
landplant_aIndex <- data.frame(AliphaticIndex=(sapply(landplant,
aIndex)))

# Hydrophobicity index (H) from Peptides
landplant_hydrophobicity <-
data.frame(Hydrophobicity=(sapply(landplant, hydrophobicity, scale =
"Eisenberg")))

# Hydrophobic moment index (mH) for Alpha helices and Beta sheets
from Peptides
landplant_mHalpha <- data.frame(mHalpha=(sapply(landplant, hmoment,
angle = 100, window = 11)))
landplant_mHbeta <- data.frame(mHbeta=(sapply(landplant, hmoment,
angle = 160, window = 11)))

# Aminoacid composition from protr
landplant_AAC <- data.frame(t(sapply(landplant, extractAAC)))

# Aminoacid composition type from Peptides
```

```
subdata <- function(x){ x[,2] }
landplant_aaComp <-
as.data.frame(t(data.frame(sapply(sapply(landplant, aaComp),
subdata))))


# merge all the descriptor data
landplantdf <- data.frame(landplant_length, landplant_mw,
landplant_charge, landplant_hydrophobicity, landplant_pI,
                          landplant_aIndex, landplant_mHalpha,
landplant_mHbeta, landplant_AAC)

landplantdatafull <- merge(landplant_metadata, landplantdf, by =
"row.names", all.x = FALSE)
rownames(landplantdatafull) <- landplantdatafull[,1];
landplantdatafull[,1] <- NULL
str(landplantdatafull)
landplantdata <- landplantdatafull[, -c(1,3)]   # Remove unnecessary
columns
str(landplantdata)
table(landplantdata$GeneFamily)
colnames(landplantdata)
# [1] "GeneFamily"     "Length"         "MolWt"          "NetCharge"
"Hydrophobicity" "pI"             "AliphaticIndex" "mHalpha"
# [9] "mHbeta"         "A"              "R"              "N"
"D"              "C"              "E"              "Q"
# [17] "G"             "H"              "I"              "L"
"K"              "M"              "F"              "P"
# [25] "S"             "T"              "W"              "Y"
"V"


# Run RandomForest
library("randomForest")
set.seed(123)
rf <- randomForest(GeneFamily ~ ., data=landplantdata,
importance=TRUE, proximity=TRUE)
print(rf)
# attributes(rf)
# rf$confusion # Confusion matrix
plot(rf)  # Error rate of Random Forest

# Variable importance
varImpPlot(rf)
varImpPlot(rf, sort = T, n.var = 10, main = "Top 10 - Variable
importance")
importance(rf)
varUsed(rf)


## ~~~~~~~~~~~~~~~~~~ RandomForestExplainer ~~~~~~~~~~~~~~~~~~ ##
library("randomForestExplainer")
explain_forest(rf, interactions = TRUE, data = landplantdata)
```

```
# ~~~~~~~~~~~~~~~~~~ Plots ~~~~~~~~~~~~~~~~~ #
x <- landplantdata[,c(1,2:9)]  # Length, MolWt, NetCharge, pI,
AliphaticIndex, Hydrophobicity, mHalpha, mHbeta
y <- landplantdata[,c(1,10:29)] # Amino acids from AAC


df.m <- melt(y, id.var = "GeneFamily")
p <- ggplot(data = df.m, aes(x=variable, y=value))
p <- p + geom_violin(aes(fill = GeneFamily))
p <- p + facet_wrap( ~ variable, scales="free")
# p <- p + xlab("Category") + ylab("Value") + ggtitle("Amino acids")
# General properties, Size and characteristics, Amino acids
p <- p + guides(fill=guide_legend(title="Gene family"))
p
```
