## Additional File 1 for "Deep Evolutionary History of the Phox and Bem1 (PB1) Domain Across Eukaryotes"

**TABLE OF CONTENTS**

**SUPPLEMENTARY FIGURES**

**Fig. S1:** Presence of various PB1 domain containing proteins identified in only one sequence and/or one species.

**Fig. S2:** Complete (A) and simplified (B) illustration of the unrooted tree with the PB1 domains from all five kingdoms.

**Fig. S3:** Violin plots showing the descriptive stats of the 28 descriptors/variables used Random Forest (RF) classification.

**SUPPLEMENTARY TABLES**

**Table S1:** List of domains (present in Fig. 1) with their short name, full name and a link to the InterPro domain database.

**Table S2:** List of identifiers of the PB1 domain containing proteins from four species of land plants used for the sequence alignment and logo construction.

**Table S3:** Confusion matrix from the Random Forest (RF) model.

**SUPPLEMENTARY FIGURES**

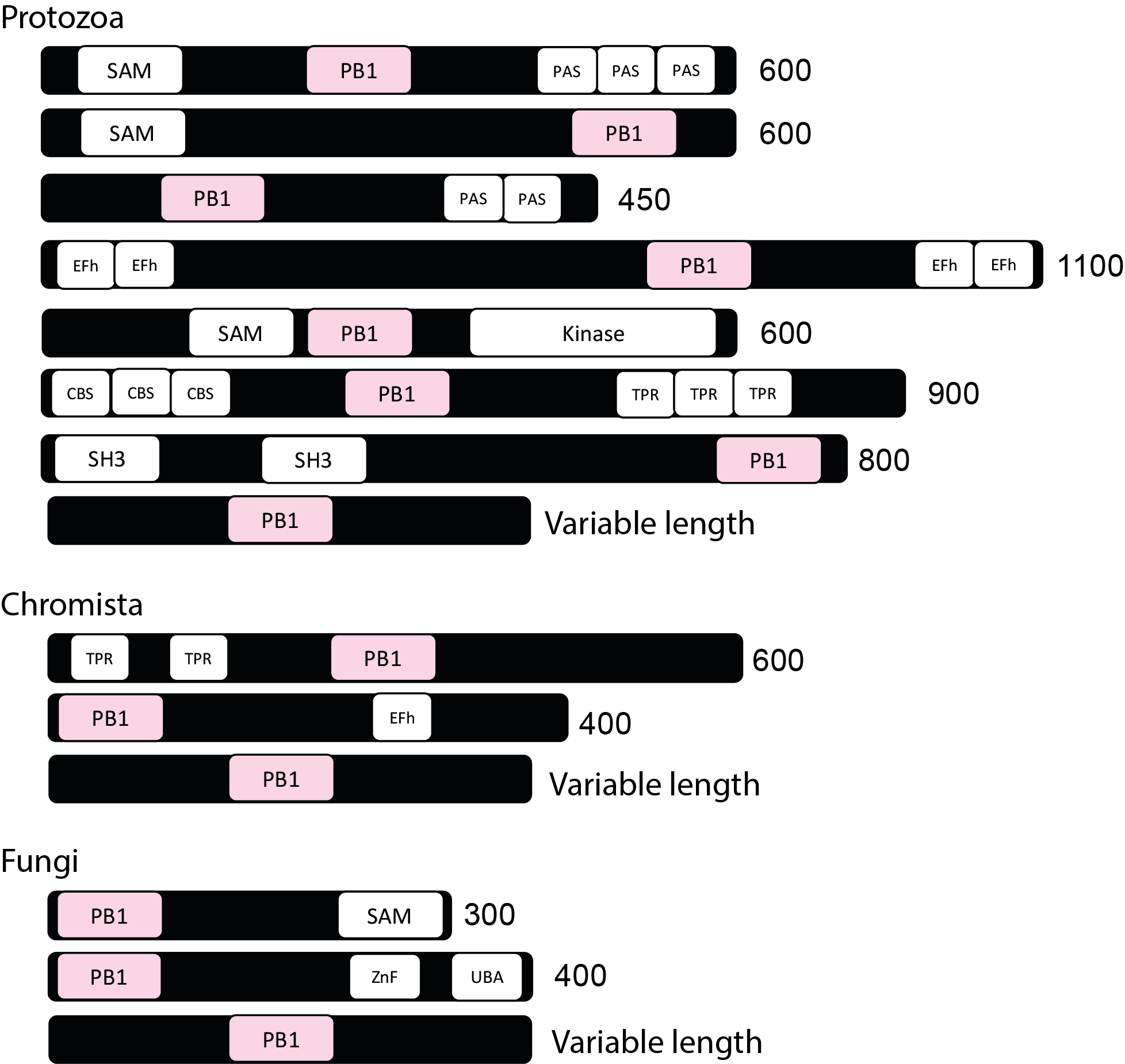

**Fig. S1:** Presence of various PB1 domain containing proteins identified in only one sequence and/or one species. The numbers at the end of each row is the approximate length of the protein. The complete information about the domains and their respective InterPro database links are provided in Table S1.

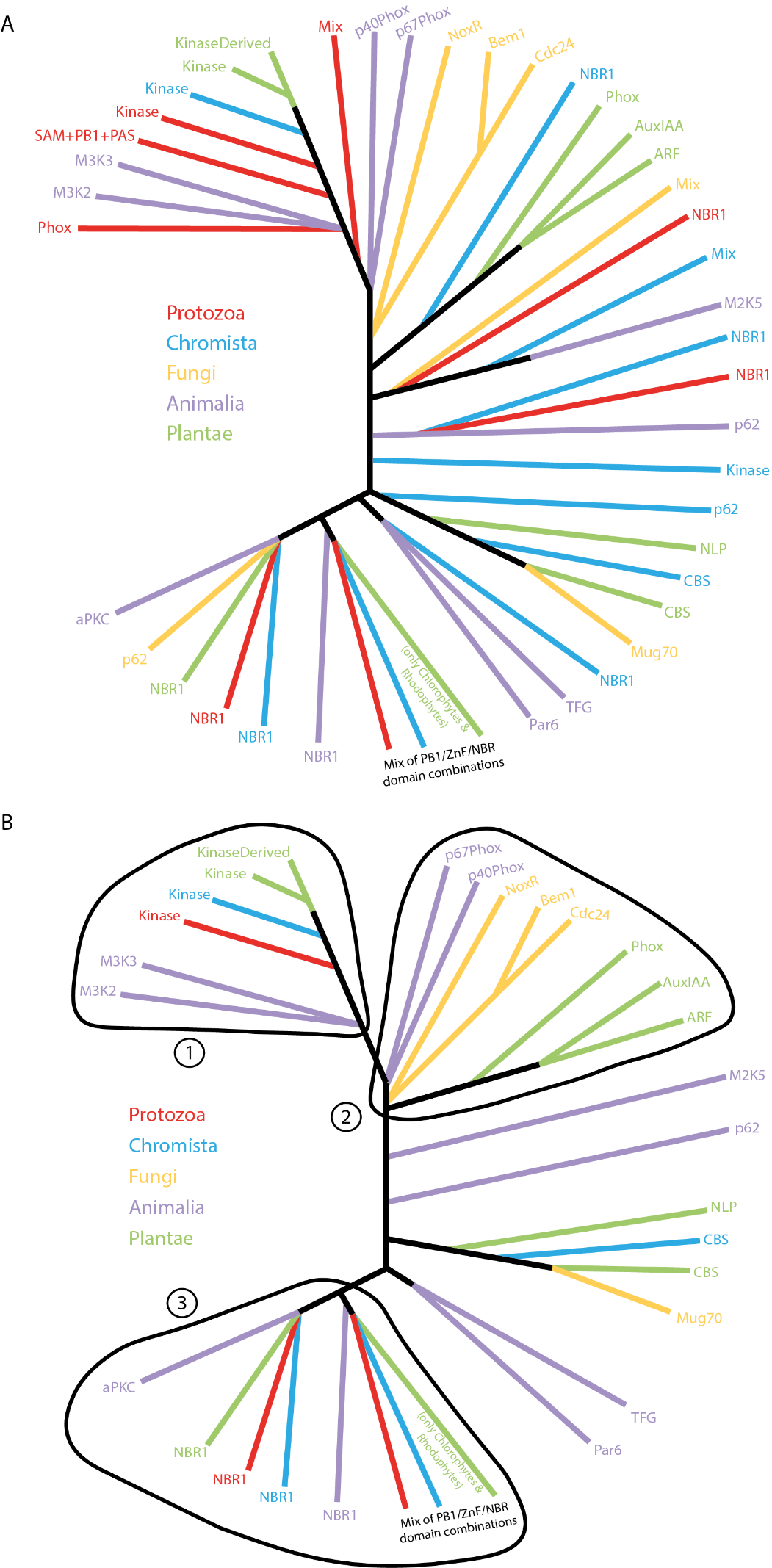

**Fig. S2:** Complete (A) and simplified (B) illustration of the unrooted tree with the PB1 domains from all five kingdoms. Orthologs from each kingdom are represented with each colour as indicated: Protozoa in ‘red’, Chromista in ‘blue’, Fungi in ‘orange’, Animalia in ‘purple’ and Plantae in ‘green’. The groups outlined with continuous lines indicated with numbers 1, 2 and 3 represent the probable ancestral copies in LECA corresponding to Kinase, Phox and NBR1 groups respectively. ‘Mix’ indicates a combination of (partial) PB1 domains with other domains in random. Full version of the tree with taxa names and domain information can be found at iTOL: https://itol.embl.de/shared/dolfweijers.

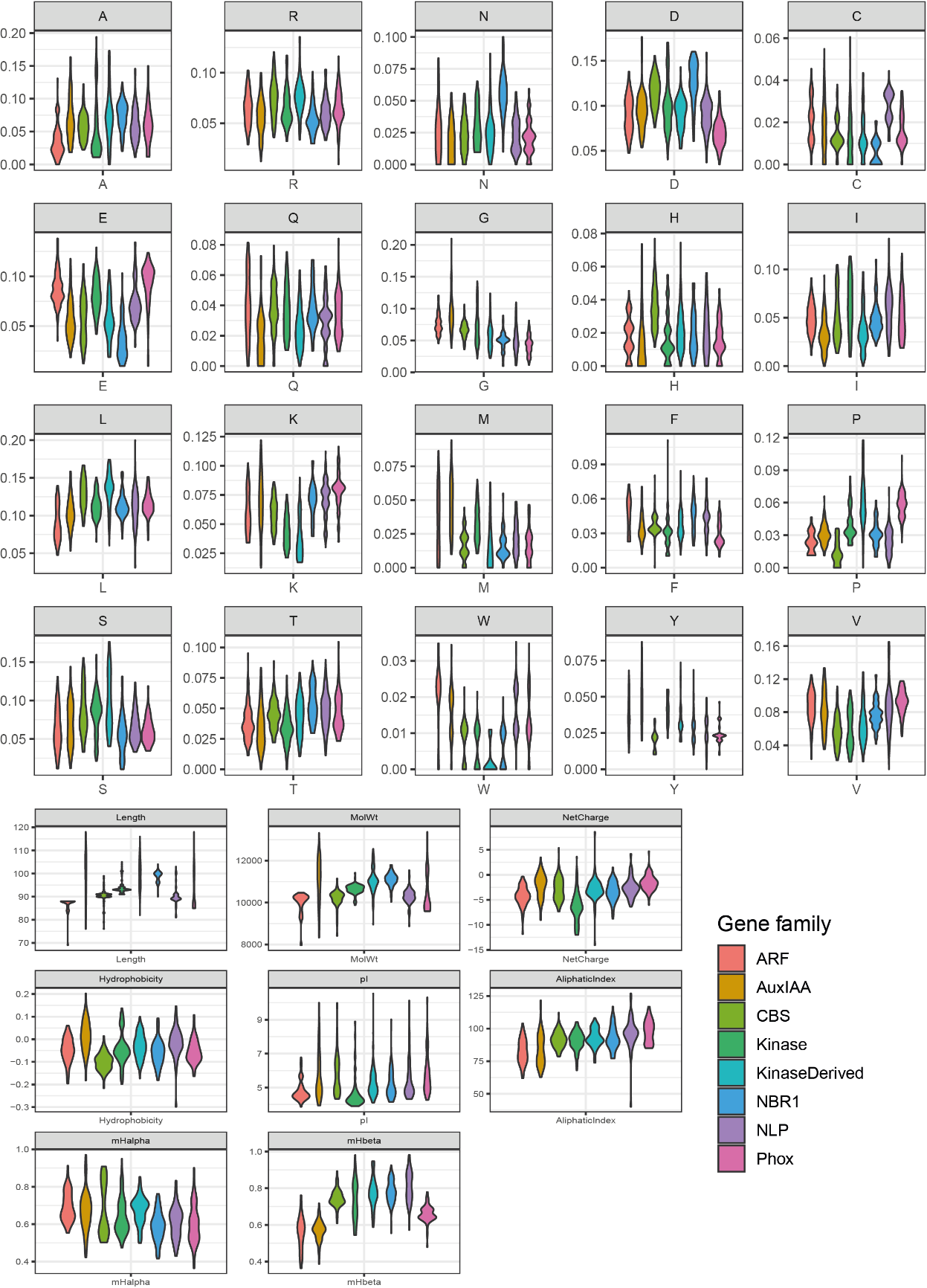

**Fig. S3:** Violin plots showing the descriptive stats of the 28 descriptors/variables used Random Forest (RF) classification. Colours in each plot represent one gene family as shown in the legend.

**SUPPLEMENTARY TABLES**

**Table S1:** List of domains (present in Fig. 1) with their short name, full name and a link to the InterPro domain database.

| **ShortName** | **FullName (InterProID)** | **InterProDB link** |
| --- | --- | --- |
| PB1 | PB1 domain (IPR000270) | <https://www.ebi.ac.uk/interpro/entry/IPR000270> |
| Kinase | Protein kinase domain (IPR000719) | <https://www.ebi.ac.uk/interpro/entry/IPR000719> |
| WD40 | WD40 repeat (IPR001680) | <https://www.ebi.ac.uk/interpro/entry/IPR001680> |
| TPR | Tetratricopeptide repeat (IPR019734) | <https://www.ebi.ac.uk/interpro/entry/IPR019734> |
| ZnF | Zinc finger, ZZ-type (IPR000433) | <http://www.ebi.ac.uk/interpro/entry/IPR000433> |
| CentralDomain | Next to BRCA1, central domain (IPR032350) | <http://www.ebi.ac.uk/interpro/entry/IPR032350> |
| UBA | Ubiquitin-associated domain (IPR015940) | <http://www.ebi.ac.uk/interpro/entry/IPR015940> |
| CBS | CBS domain (IPR000644) | <http://www.ebi.ac.uk/interpro/entry/IPR000644> |
| RWPRK | RWP-RK domain (IPR003035) | <http://www.ebi.ac.uk/interpro/entry/IPR003035> |
| DH | Dbl homology (DH) domain (IPR000219) | <http://www.ebi.ac.uk/interpro/entry/IPR000219> |
| PH | Pleckstrin homology domain (IPR001849) | <http://www.ebi.ac.uk/interpro/entry/IPR001849> |
| SH3 | SH3 domain (IPR001452) | <http://www.ebi.ac.uk/interpro/entry/IPR001452> |
| PX | Phox homologous domain (IPR001683) | <http://www.ebi.ac.uk/interpro/entry/IPR001683> |
| kDAG | Phorbol ester/diacylglycerol-binding domain (IPR002219) | <http://www.ebi.ac.uk/interpro/entry/IPR002219> |
| PDZ | PDZ domain (IPR001478) | <http://www.ebi.ac.uk/interpro/entry/IPR001478> |
| ARF-DBD/B3 | B3 DNA binding domain (IPR003340) | <http://www.ebi.ac.uk/interpro/entry/IPR003340> |
| ARF-DBD/ARF | Auxin response factor (IPR010525) | <http://www.ebi.ac.uk/interpro/entry/IPR010525> |
| Aux/IAA-I | Domain-I or EAR motif | NA |
| Aux/IAA-II | Domain-II or DEGRON motif | NA |

**Table S2:** List of identifiers of the PB1 domain containing proteins from four species of land plants (Marchantia, Physcomitrella, Amborella and Arabidopsis) used for the sequence alignment and logo construction.

| **GeneFamily** | **Marchantia** | **Physcomitrella** | **Amborella** | **Arabidopsis** |
| --- | --- | --- | --- | --- |
| ARF | Mapoly0019s0045.1 | Pp3c1_14480V3.1 | evm_27.model.AmTr_v1.0_scaffold00007.382 | AT1G59750 |
|  | Mapoly0011s0167.1 | Pp3c1_40270V3.1 | evm_27.model.AmTr_v1.0_scaffold00016.128 | AT5G62000 |
|  | Mapoly0075s0050.1 | Pp3c2_25890V3.1 | evm_27.model.AmTr_v1.0_scaffold00021.210 | AT5G60450 |
|  |  | Pp3c4_12970V3.3 | evm_27.model.AmTr_v1.0_scaffold00025.251 | AT1G19850 |
|  |  | Pp3c4_13010V3.3 | evm_27.model.AmTr_v1.0_scaffold00029.187 | AT1G30330 |
|  |  | Pp3c5_9420V3.1 | evm_27.model.AmTr_v1.0_scaffold00057.126 | AT5G20730 |
|  |  | Pp3c6_21370V3.1 | evm_27.model.AmTr_v1.0_scaffold00092.36 | AT5G37020 |
|  |  | Pp3c13_4720V3.1 | evm_27.model.AmTr_v1.0_scaffold00148.24 | AT4G23980 |
|  |  | Pp3c14_16990V3.10 | evm_27.model.AmTr_v1.0_scaffold00155.56 | AT2G28350 |
|  |  | Pp3c16_6100V3.1 | evm_27.model.AmTr_v1.0_scaffold00211.4 | AT2G46530 |
|  |  | Pp3c17_19900V3.1 |  | AT1G34310 |
|  |  | Pp3c27_60V3.1 |  | AT1G34170 |
|  |  | Pp3c9_21330V3.1 |  | AT1G35540 |
|  |  | Pp3c15_9710V3.1 |  | AT1G35520 |
|  |  |  |  | AT4G30080 |
|  |  |  |  | AT3G61830 |
|  |  |  |  | AT1G19220 |
|  |  |  |  | AT1G35240 |
|  |  |  |  | AT1G34410 |
|  |  |  |  | AT1G34390 |
| AuxIAA | Mapoly0013s0010.1 | Pp3c24_6610V3.1 | evm_27.model.AmTr_v1.0_scaffold00002.512 | AT4G14560 |
|  | Mapoly0034s0017.1 | Pp3c8_14720V3.1 | evm_27.model.AmTr_v1.0_scaffold00002.514 | AT3G23030 |
|  |  |  | evm_27.model.AmTr_v1.0_scaffold00019.282 | AT1G04240 |
|  |  |  | evm_27.model.AmTr_v1.0_scaffold00039.160 | AT5G43700 |
|  |  |  | evm_27.model.AmTr_v1.0_scaffold00045.141 | AT1G15580 |
|  |  |  | evm_27.model.AmTr_v1.0_scaffold00056.118 | AT1G52830 |
|  |  |  | evm_27.model.AmTr_v1.0_scaffold00109.120 | AT3G23050 |
|  |  |  | evm_27.model.AmTr_v1.0_scaffold00122.5 | AT2G22670 |
|  |  |  | evm_27.model.AmTr_v1.0_scaffold00184.12 | AT5G65670 |
|  |  |  |  | AT1G04100 |
|  |  |  |  | AT4G28640 |
|  |  |  |  | AT1G04550 |
|  |  |  |  | AT2G33310 |
|  |  |  |  | AT4G14550 |
|  |  |  |  | AT3G04730 |
|  |  |  |  | AT1G04250 |
|  |  |  |  | AT1G51950 |
|  |  |  |  | AT3G15540 |
|  |  |  |  | AT2G46990 |
|  |  |  |  | AT3G16500 |
|  |  |  |  | AT4G29080 |
|  |  |  |  | AT5G25890 |
|  |  |  |  | AT4G32280 |
|  |  |  |  | AT3G62100 |
|  |  |  |  | AT3G17600 |
|  |  |  |  | AT2G01200 |
|  |  |  |  | AT5G57420 |
|  |  |  |  | AT1G15050 |
| CBS | Mapoly0179s0023.1 | Pp3c1_14290V3.1 | evm_27.model.AmTr_v1.0_scaffold00013.41 | AT5G63490 |
|  |  | Pp3c1_14310V3.1 | evm_27.model.AmTr_v1.0_scaffold00017.72 | AT2G36500 |
|  |  | Pp3c11_15160V3.1 |  | AT3G52950 |
|  |  | Pp3c2_26110V3.1 |  | AT5G50640 |
|  |  | Pp3c7_10070V3.1 |  |  |
| Kinase | Mapoly0013s0150.1 | Pp3c15_24250V3.1 | evm_27.model.AmTr_v1.0_scaffold00004.293 | AT1G04700 |
|  |  | Pp3c9_25280V3.1 | evm_27.model.AmTr_v1.0_scaffold00019.236 | AT1G16270 |
|  |  |  | evm_27.model.AmTr_v1.0_scaffold00026.90 | AT1G79570 |
|  |  |  | evm_27.model.AmTr_v1.0_scaffold00039.196 | AT2G35050 |
|  |  |  | evm_27.model.AmTr_v1.0_scaffold00081.26 | AT3G24715 |
|  |  |  |  | AT3G46920 |
|  |  |  |  | AT5G57610 |
| KinaseDerived |  |  | evm_27.model.AmTr_v1.0_scaffold00007.258 | AT1G25300 |
|  |  |  | evm_27.model.AmTr_v1.0_scaffold00046.178 | AT1G70640 |
|  |  |  | evm_27.model.AmTr_v1.0_scaffold00049.221 | AT2G01190 |
|  |  |  | evm_27.model.AmTr_v1.0_scaffold00109.93 | AT3G18230 |
|  |  |  |  | AT3G26510 |
|  |  |  |  | AT3G48240 |
|  |  |  |  | AT4G05150 |
|  |  |  |  | AT5G09620 |
|  |  |  |  | AT5G16220 |
|  |  |  |  | AT5G49920 |
|  |  |  |  | AT5G63130 |
|  |  |  |  | AT5G64430 |
| NBR1 | Mapoly0100s0042.1 | Pp3c11_16970V3.1 | evm_27.model.AmTr_v1.0_scaffold00049.238 | AT4G24690 |
|  |  | Pp3c7_8990V3.1 |  |  |
| NLP | Mapoly0083s0040.1 | Pp3c12_2070V3.1 | evm_27.model.AmTr_v1.0_scaffold00058.115 | AT2G17150 |
|  |  | Pp3c15_9180V3.1 | evm_27.model.AmTr_v1.0_scaffold00066.150 | AT4G35270 |
|  |  | Pp3c17_4370V3.1 | evm_27.model.AmTr_v1.0_scaffold00080.66 | AT4G38340 |
|  |  | Pp3c17_4375V3.1 |  | AT1G20640 |
|  |  | Pp3c19_2670V3.1 |  | AT1G76350 |
|  |  | Pp3c19_2720V3.1 |  | AT1G64530 |
|  |  | Pp3c22_6360V3.1 |  | AT4G24020 |
|  |  | Pp3c22_6370V3.1 |  | AT2G43500 |
|  |  | Pp3c9_14600V3.1 |  | AT3G59580 |

**Table S3:** Confusion matrix from the Random Forest (RF) model. Diagonal values represent the correctly classified PB1’s, and others represent the mis-classified category. The column ‘class.error’ represents the classification error for that particular class of PB1’s as shown in Fig. 5a.

|  | **ARF** | **AuxIAA** | **CBS** | **Kinase** | **KinaseDerived** | **NBR1** | **NLP** | **Phox** | **class.error** |
| --- | --- | --- | --- | --- | --- | --- | --- | --- | --- |
| **ARF** | 99 | 0 | 1 | 0 | 0 | 0 | 0 | 0 | 0.01 |
| **AuxIAA** | 4 | 95 | 1 | 0 | 0 | 0 | 0 | 0 | 0.05 |
| **CBS** | 2 | 1 | 92 | 1 | 0 | 2 | 2 | 0 | 0.08 |
| **Kinase** | 0 | 1 | 0 | 95 | 4 | 0 | 0 | 0 | 0.05 |
| **KinaseDerived** | 1 | 0 | 2 | 7 | 87 | 0 | 1 | 2 | 0.13 |
| **NBR1** | 0 | 2 | 2 | 1 | 1 | 72 | 0 | 0 | 0.07 |
| **NLP** | 0 | 0 | 3 | 0 | 1 | 0 | 95 | 1 | 0.05 |
| **Phox** | 0 | 2 | 0 | 0 | 0 | 0 | 2 | 96 | 0.04 |
