## Supplementary figures and images for "Deep Evolutionary History of the Phox and Bem1 (PB1) Domain Across Eukaryotes"

### Additional File 2

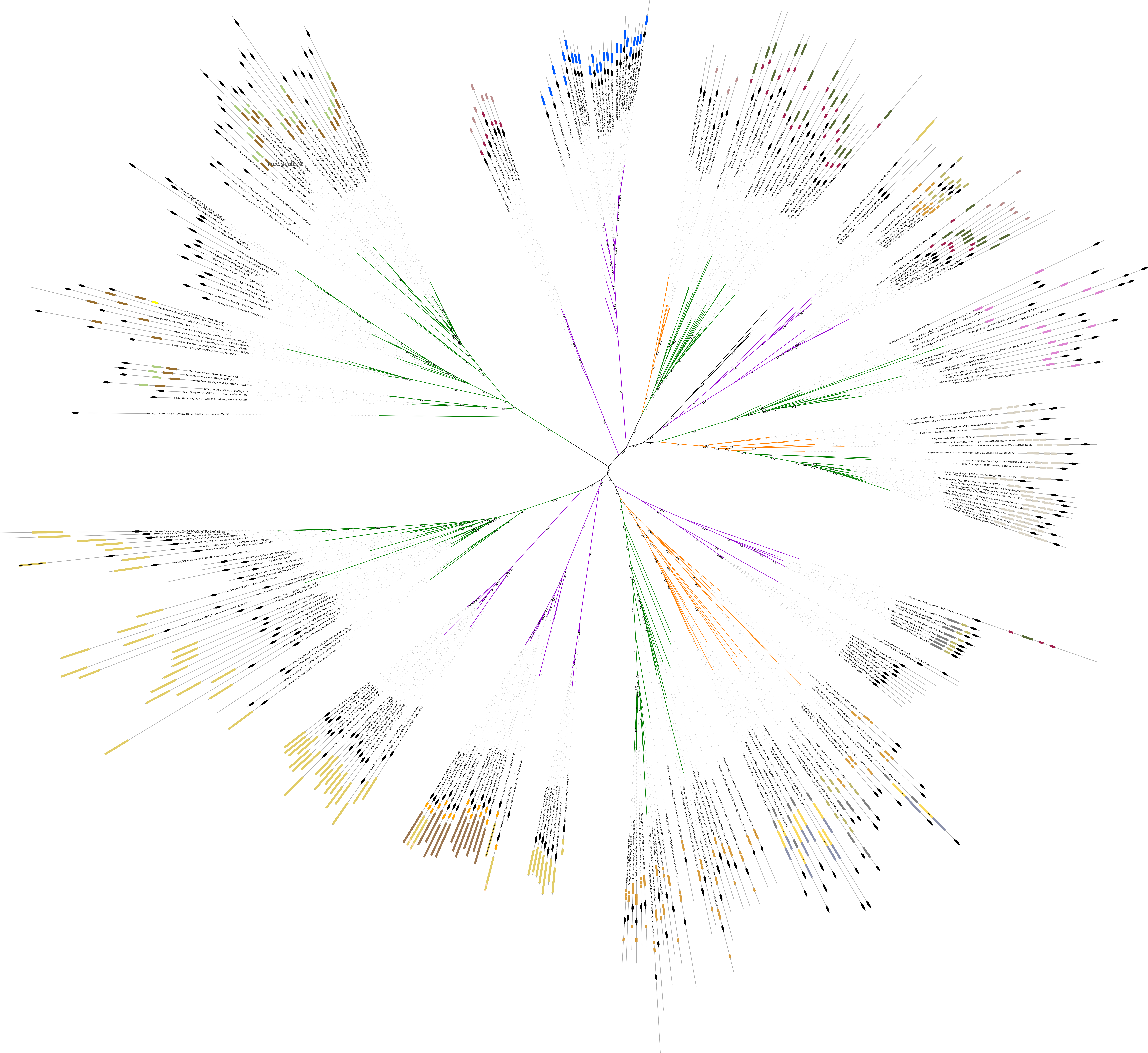

### Additional File 3

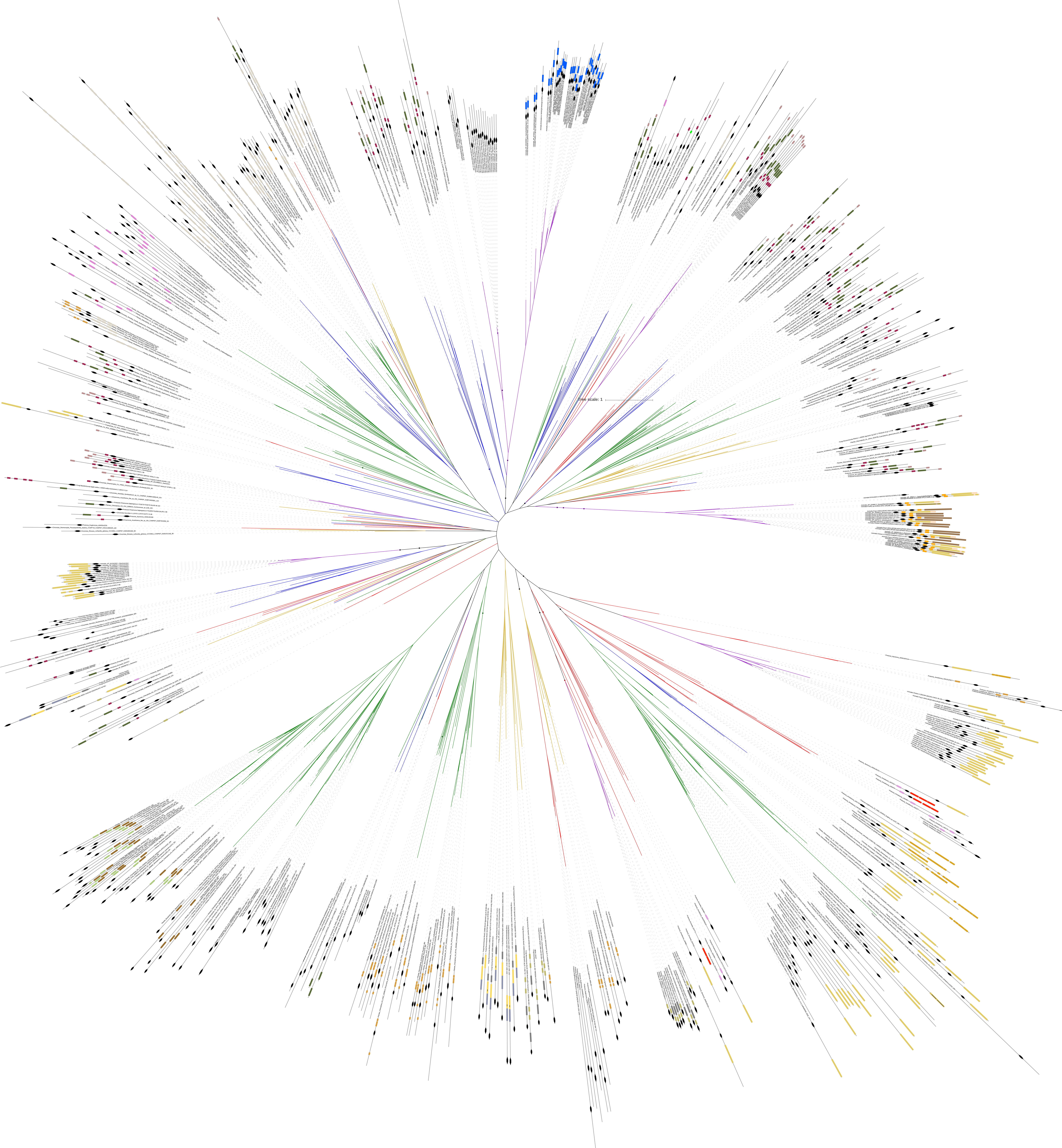
